# Small non-coding RNA CjNC110 influences motility, autoagglutination, AI-2 localization, hydrogen peroxide sensitivity and chicken colonization in *Campylobacter jejuni*

**DOI:** 10.1101/2020.01.02.893479

**Authors:** Amanda J. Kreuder, Brandon Ruddell, Kathy Mou, Alan Hassall, Qijing Zhang, Paul J. Plummer

## Abstract

Small non-coding RNAs are involved in many important physiological functions in pathogenic microorganisms. Previous studies have identified the presence of non-coding RNAs in the major zoonotic pathogen *Campylobacter jejuni*, however, few have been functionally characterized to date. CjNC110 is a conserved ncRNA in *C. jejuni,* located downstream of the *luxS* gene which is responsible for the production of the quorum-sensing molecule autoinducer-2 (AI-2). In this study, we utilized strand specific high-throughput RNAseq to identify potential targets or interactive partners of CjNC110 in a sheep abortion clone of *C. jejuni*. This data was then utilized to focus further phenotypic evaluation of the role of CjNC110 in motility, autoagglutination, quorum sensing, hydrogen peroxide sensitivity and chicken colonization in *C. jejuni*. Inactivation of the CjNC110 ncRNA led to a statistically significant decrease in autoagglutination ability as well as increased motility and hydrogen peroxide sensitivity when compared to wild-type. Extracellular AI-2 detection was decreased in ΔCjNC110, however, intracellular AI-2 accumulation was significantly increased, suggesting a key role of CjNC110 in modulating the transport of AI-2. Notably, ΔCjNC110 also showed a decreased ability to colonize chickens. Complementation of CjNC110 restored all phenotypic changes back to wild-type levels. The collective results of the phenotypic and transcriptomic changes observed in our data provide valuable insights into the pathobiology of *C. jejuni* sheep abortion clone and strongly suggest that CjNC110 plays an important role in regulation of energy taxis, flagellar glycosylation, cellular communication via quorum sensing, oxidative stress tolerance and chicken colonization in this important zoonotic pathogen.

## INTRODUCTION

*Campylobacter jejuni* is a major foodborne and zoonotic pathogen that causes enteritis in humans (1). A sheep abortion (SA) clone, with IA3902 as the prototype strain, has recently emerged as the predominant cause of ovine abortion and as an important pathogen in foodborne outbreaks of human gastroenteritis in the United States (2, 3). *C. jejuni* clone SA is also widely distributed in the cattle population in the U.S. (4). The genome of IA3902 is quite syntenic with that of *C. jejuni* type-strain NCTC 11168 although it is much more virulent than NCTC 11168 in inducing systemic infection and abortion in animals (5). Despite its hypervirulence, *C. jejuni* IA3902 does not harbor any virulence factors known to be associated with abortion in *C. fetus* subsp. *fetus* (6, 7). Previously, the *luxS* gene, which mediates autoinducer-2 (AI-2) production, and the genes encoding the capsular polysaccharide have been identified as critical in IA3902 for intestinal colonization and/or translocation across gut epithelium into the bloodstream (8, 9). Point mutations in the major outer membrane protein encoded by *porA* of IA3902 have also been shown to be sufficient to cause the abortion phenotype when compared to the syntenic non-abortive strain NCTC 11168 (10). The fact that relatively mild changes in genomic sequences have led to significantly enhanced ability to cause disease by *C. jejuni* IA3902, as described above, suggests that even slight differences in sequence variations or gene regulation may have a major impact on virulence variation among differing strains of *C. jejuni*.

*C. jejuni* has only three known sigma factors that regulate gene transcription: σ^70^ (encoded by *rpoD*), σ^54^ (encoded by *rpoN*) and σ^28^ (encoded by *fliA*) (11). Besides control at the transcriptional level, regulation of gene expression can also occur by post-transcriptional modulation of mRNA translation, stability and processing, for which small non-coding RNAs are the primary players (12–14). Non-coding RNAs (ncRNAs) can be rapidly produced and serve to regulate multiple different targets within a cell in a variety of ways to coordinate rapid responses to changing environments. Thus, regulation of cellular processes by ncRNAs can provide several advantages to the bacteria when compared to the traditional model of protein-mediated regulation (15).

Prior to completion of the transcriptional start site map via high throughput RNA sequencing (RNAseq) of *H. pylori* (16), the ε-proteobacteria were thought not to be capable of using small and antisense non-coding RNA as a regulatory mechanism, partly due to a lack of the RNA chaperone protein Hfq (17). Indeed, initial attempts using computational approaches to identify ncRNAs in *Campylobacter* failed to yield any potential candidates, with only 3 potential loci being identified in *Helicobacter* (18). Recently, however, clear evidence that *C. jejuni* also harbors these important regulators has been published, revealing the existence and expression of a wealth of ncRNAs present in strains such as NCTC 11168, 81-176, 81116, RM1221 and IA3902 (19–24). Dugar *et al.* (21), when comparing the transcriptomes of 4 different *C. jejuni* isolates, observed a large variation in transcriptional start sites as well as expression patterns of both mRNA and ncRNA between strains. This suggests that variations in the existence and expression of ncRNAs even among closely related strains may play a role in differentiating virulence. Despite these advances, identification of the functions of ncRNAs in *Campylobacter* has been slow to follow, with most lacking both functional characterization as well as mechanistic investigation.

The first report attempting to elucidate the function of two previously identified non-coding RNAs in *C. jejuni* suggests these ncRNAs may play a role in flagellar biosynthesis; however, the authors were unable to demonstrate phenotypic changes following inactivation of either non-coding RNA (25). A later paper, however, did establish the role of an RNA antitoxin (*cjrA*) as the first non-coding antisense small RNA functionally characterized in *Campylobacter* to date (26). Just recently, the small RNA pair, CjNC180/CjNC190, has been identified to alter colonization in a newly developed 3D *in vitro* model of disease in strain NCTC 11168 (27); the mechanism by which this occurs remains to be fully determined. Thus, beyond simply establishing the existence of non-coding RNA transcripts in *Campylobacter*, there is critical need to continue to determine the physiological functions of these potential regulators in this important zoonotic pathogen.

Our previous work has demonstrated the *in vivo* and *in vitro* expression of numerous ncRNAs in IA3902 (24), including several that are conserved in other strains of *C. jejuni* (21). One of the expressed ncRNAs identified in our study, the conserved small RNA CjNC110, is located in the intergenic region immediately downstream of the *luxS* gene. This ncRNA was of particular interest to our group as previous work in our lab has already highlighted the importance of the *luxS* gene in the virulence of *C. jejuni* IA3902 (8). In *Campylobacter*, the *luxS* gene is known to serve two important functions, production of the quorum sensing molecule autoinducer-2 (AI-2) and conversion of *S*-ribosylhomocysteine to homocysteine in the activated methyl cycle (AMC) (28). Of particular note, following the identification of CjNC110, it has also been suggested that different methods of generation of the *luxS* mutation may have led to polar effects on this downstream ncRNA that may help explain observed differences in phenotypes and gene expression between various studies of *luxS* mutants in *Campylobacter* (29).

Based on these observations, in this study we chose to focus on beginning to elucidate the functional role of CjNC110 in the pathobiology of *C. jejuni* IA3902, as well as investigate the transcriptomic and phenotypic differences between single mutation of CjNC110 and *luxS* versus co-mutation of both genomic regions. Using high-throughput RNA sequencing (RNAseq), we successfully demonstrated differences in the transcriptional landscape following mutagenesis of CjNC110 and identified a number of potential mRNA targets for CjNC110 regulation. From these potential targets, we then demonstrated that inactivation of CjNC110 affects several important phenotypes in IA3902, including motility, autoagglutination activity, AI-2 localization, hydrogen peroxide sensitivity, and chicken colonization; our results also demonstrate that these phenotypes consistently differ from those of seen with inactivation of *luxS*. The work presented here provides compelling evidence that expression of the ncRNA CjNC110 is important for the pathobiology of *C. jejuni* IA3902, and lays the foundation for future work investigating the mechanism of regulation of post-transcriptional gene expression in IA3902 by CjNC110.

## RESULTS

### Northern blot analysis validates expression of small ncRNA CjNC110 in IA3902 wild-type and mutant constructs ΔluxS and ΔCjNC110c, and absence in ΔCjNC110

Expression of CjNC110 had been previously validated via northern blot analysis in several strains of *C. jejuni*, including NCTC 11168, RM1221, 81-176, and 81116, and further genetic analysis suggested CjNC110 to be present in IA3902 as well (21). Therefore, to begin to study the role of CjNC110 in IA3902, insertional deletion of the entire predicted coding sequence was utilized to construct ΔCjNC110 in IA3902, and complementation was achieved via re-insertion of CjNC110 into the 16S and 23S rRNA operon (*rrs-rrl*) of ΔCjNC110 via homologous recombination using plasmid pRRK, creating ΔCjNC110c (**Table 1** lists all strains used in this study). To validate expression of CjNC110 in IA3902, an initial northern blot using the same probe sequence as Dugar *et al*. (21) and 15 μg of total RNA extracted at early stationary phase of growth from cultures collected at the same A_600_ was performed to compare IA3902 wild-type, ΔCjNC110 and ΔCjNC110c. The results of this northern blot analysis clearly demonstrate that small ncRNA CjNC110 is expressed by IA3902 wild-type and ΔCjNC110c but not ΔCjNC110 **(Fig. 1)**, validating expression of CjNC110 in IA3902 and confirming successful mutant and complement constructs. Visual inspection of the northern blot data from both Dugar *et al.* and the work presented here also demonstrates the presence of multiple bands of differing sizes in both studies (21). All bands were eliminated in the ΔCjNC110 mutant and restored in the ΔCjNC110c complement, which suggests processing of the CjNC110 transcript may be occurring; further work is needed to investigate this observation.

**FIG 1.**
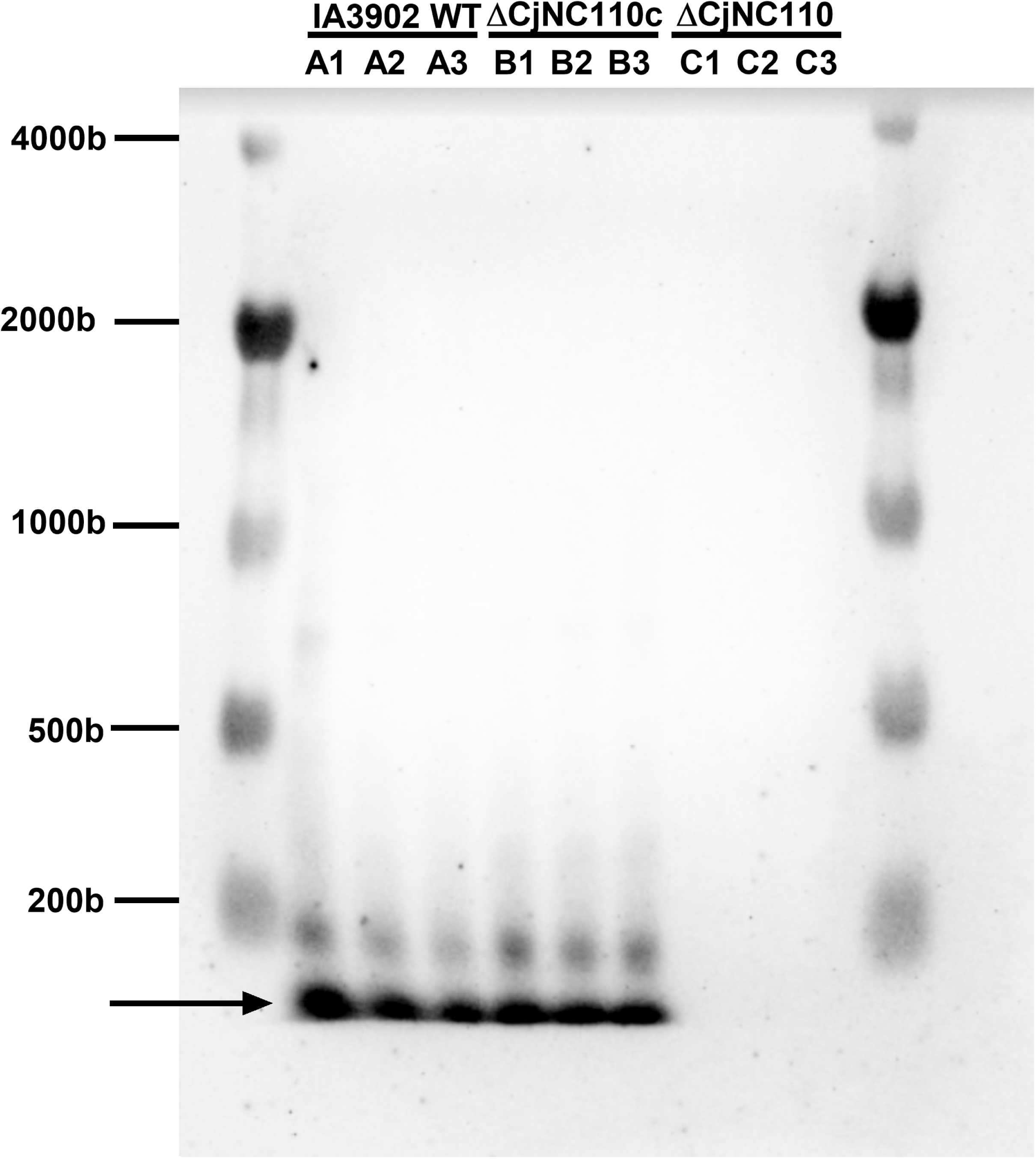
Northern blotting demonstrates expression of CjNC110 in wild-type IA3902 and ΔCjNC110c and absence of expression in ΔCjNC110. Cultures for RNA extraction were collected at early stationary phase of growth and northern blot analysis was conducted using 15 μg of total RNA in each of three separate replicates per strain tested. The arrow indicates the most prominent band which corresponds to the previously predicted size of CjNC110 in *C. jejuni* (21).

**TABLE 1.** Bacterial strains utilized in this study.

| Strain | Description | Source or Reference |
| --- | --- | --- |
| ***Campylobacter jejuni*** |  |  |
| W7 | Wild-type motile variant of NCTC 11168 | (8) |
| W7 ΔCjNC110 | W7 ΔCjNC110::Cm<sup>R</sup> | This study |
| Sheep Abortion (SA) IA3902 | Wild-type *C. jejuni* | (3) |
| IA3902 ΔCjNC110 | IA3902 ΔCjNC110::Cm<sup>R</sup> | This study |
| IA3902 ΔluxS | IA3902 ΔluxS::Kan<sup>R</sup> | (33) |
| IA3902 ΔCjNC110ΔluxS | IA3902 ΔCjNC110::Cm<sup>R</sup> ΔluxS::Kan<sup>R</sup> | This study |
| IA3902 ΔCjNC110c | IA3902 ΔCjNC110::Cm<sup>R</sup> CjNC110::Kan<sup>R</sup> | This study |
| ***Escherichia coli*** |  |  |
| DH5α | *fhuA2* Δ(*argF-lacZ*)U169 *phoA glnV44* Φ80<br>Δ(*lacZ*)M15 *gyrA96 recA1 relA1 endA1 thi-1 hsdR17* | New England Biolabs,<br>Ipswich, MA |
| ***Vibrio harveyi*** |  |  |
| BB152 | AI-1-/AI-2+; luxM::Tn5 | (74) |
| BB170 | AI-2 reporter strain; luxN::Tn5 | (93) |

Kan<sup>R</sup> = kanamycin resistance cassette
Cm<sup>R</sup> = chloramphenicol resistance cassette

As previous studies have questioned the method of mutant generation of ΔluxS on CjNC110 expression (29), a second northern blot comparing wild-type IA3902 to a previously constructed ΔluxS mutant utilized by our lab (8) was performed and confirmed that CjNC110 expression in the IA3902ΔluxS mutant is also equivalent to wild-type levels (**Fig. S1**).

Homologous recombination was utilized to transfer the CjNC110 mutation into this ΔluxS mutant to create the double-knockout mutant, ΔCjNC110ΔluxS, with the goal of comparing the transcriptomic and phenotypic differences between mutation of ΔluxS or ΔCjNC110 alone versus in combination (ΔCjNC110ΔluxS).

### Differential gene expression analysis of RNAseq data suggests several potential regulatory targets for CjNC110

To begin to identify potential regulatory targets for CjNC110, RNAseq was utilized to compare gene expression changes between ΔCjNC110, ΔluxS and ΔCjNC110ΔluxS mutants. Total RNA was isolated from an *in vitro* growth curve performed in triplicate and colony counts (CFU/mL) over time were used to select samples representative of exponential (3 hours) and early stationary (12 hours) phases of growth (**Fig. S2A**). ANOVA did identify a statistically significant difference in growth between strains (*P <*0.05), however, multiple comparison analysis of individual time points and strains when compared to wild-type growth revealed that the only significant difference was a decrease in the A_600_ of ΔCjNC110 at 30 hours (**Fig S2B)** which represents the point where a decline in the A_600_ was noted to occur in all strains tested. Further examination of the actual CFU/mL over the course of the growth curve revealed that by 30 hours cell death was beginning to occur and colony counts were decreasing. As further growth is assumed to be ceased at this time, there is likely minimal effect of this difference on samples collected during growth prior to 24 hours, however, there is possibly biological significance to the difference noted during the late stationary phase which may warrant further investigation. It should be also be noted that the use of CFU/mL more clearly allowed for delineation of actual growth stage of each strain as the A_600_ results were noted to significantly lag the results of actual colony counts; therefore, the CFU/mL data was utilized to determine selection of timepoints for RNAseq.

Sequencing of the total RNA libraries prepared from the growth curves was performed on an Illumina HiSeq 2500 in high-output single read mode with 100 cycles, and Rockhopper was utilized to analyze the resulting data for differential gene expression between the mutants (30). Overall, 24 barcoded libraries were sequenced yielding over 109 million reads, with close to 100 million high quality reads aligning to either the genome or pVir plasmid of *C. jejuni* IA3902 and averaging 4,553,847 reads per library (**Tables S1 and S2**). The vast majority of reads (average of 83% of total reads) mapped to protein coding genes of the chromosome, with only 7% of reads mapping to ribosomal RNA on average following rRNA depletion with Ribo-Zero (median of 3%). Over half of the libraries contained less than or equal to 3% ribosomal RNA reads, which is consistent with the manufacturer’s predicted rRNA removal efficiency. One third of the libraries did not exhibit efficient rRNA removal (>10% rRNA reads); the reason for this is unclear. Despite this difference in efficiency, all samples provided a more than adequate number of non-rRNA reads and thus were successfully utilized in the analysis. Less than 1% of reads mapped to antisense regions of the annotated genome. Rockhopper also identified a number of putative ncRNAs in the dataset; **Table S3** includes the curated list of identified ncRNA candidates.

Following computational analysis via Rockhopper, a change in gene expression was deemed significant when the *Q*-value (false discovery rate) was <0.05 and a >1.5 fold change in expression was observed. A summary of the differences in numbers of genes with significantly increased and decreased expression in the various strains and timepoints is given in **Table 2**, with a listing of specific genes given in **Table 3**. In the ΔCjNC110 mutant strain, six genes were found to be downregulated and four genes were found to be upregulated when compared to the IA3902 wild-type strain during exponential growth (complete information **Table S4)**. In addition, a previously described ncRNA, CjNC140 (21) was found to be upregulated in the mutant condition. During early stationary phase, 16 genes were found to be downregulated and seven genes were found to be upregulated in the mutant strain when compared to the wild-type strain. Of the differentially expressed genes, three genes [*neuB2, hisF, ptmA*] were downregulated in the ΔCjNC110 mutant during both the exponential and stationary phases, and one, CJSA_1261 was found to be upregulated in both conditions. In addition, five separate operons predicted by Rockhopper demonstrated multiple genes within the operon affected by the mutant condition. Analysis of functionality via the COG database revealed that multiple upregulated genes (*luxS, cetA*, *cetB*) were present in the “Signal transduction mechanisms” category; multiple downregulated genes were included in the “Cell wall/membrane biogenesis” (*neuB2, ptmB,* CJSA_1352) and “Post-translational modification, protein turnover, chaperones” (*tpx,* CJSA_0687) functional categories. These differentially expressed genes represent potential targets of CjNC110 regulation in IA3902, and thus were used to inform further phenotypic study of the function of this ncRNA. Genes highlighted in bold in **Table 3** indicate potential mRNA targets of CjNC110 that were further investigated using phenotypic assays.

**TABLE 2.** Summary of differential gene expression results between mutant strains.

|  | Condition |  |  |  |  |  |
| --- | --- | --- | --- | --- | --- | --- |
|  | ΔCjNC110 |  | ΔluxS |  | ΔCjNC110ΔluxS |  |
|  | 3 h | 12 h | 3 h | 12 h | 3 h | 12 h |
| **Protein-coding genes** |  |  |  |  |  |  |
| Number downregulated | 6 | 16 | 1 | 6 | 61 | 32 |
| Number upregulated | 4 | 7 | 14 | 6 | 47 | 29 |
| **Non-coding RNA genes** |  |  |  |  |  |  |
| Number downregulated | 0 | 0 | 0 | 0 | 25<sup>b</sup> | 2 |
| Number upregulated | 1<sup>a</sup> | 0 | 1<sup>a</sup> | 0 | 1<sup>a</sup> | 0 |

<sup>a</sup> = previously described ncRNA, CjNC140 (21)
<sup>b</sup> = 17 tRNA genes, 3 known RNA genes (tmRNA, SRP, 6S) and 4 newly predicted ncRNA

**TABLE 3.**
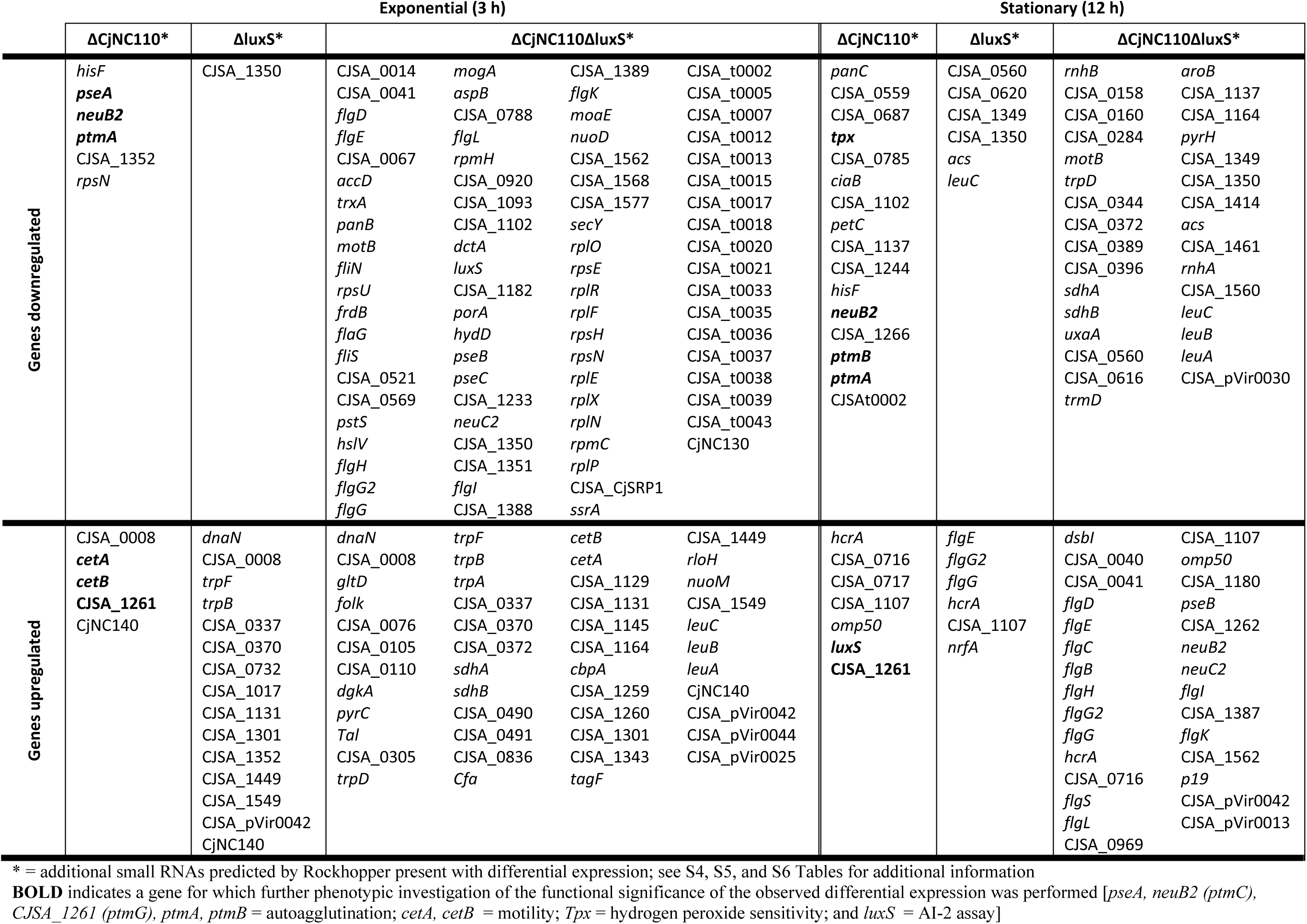
Summary of differential gene expression (>1.5 fold change, *Q* <0.05) in the IA3902 ΔCjNC110, ΔluxS and ΔCjNC110ΔluxS mutants as determined by RNAseq and analyzed via Rockhopper.

### ΔluxS displays minimal alterations in gene expression via RNAseq, with more significant gene expression changes observed in ΔCjNC110ΔluxS

Differential gene expression analysis via RNAseq and Rockhopper was also performed for ΔluxS and ΔCjNC110ΔluxS and compared to wild-type IA3902 to assess the difference between single mutation of *luxS* and concurrent mutation of both CjNC110 and *luxS*. In the ΔluxS mutant, a surprisingly small number of genes were noted to be differentially expressed. One gene was found to be downregulated and 14 genes were found to be upregulated when compared to IA3902 wild-type during exponential growth (summary **Table 3**, complete information **Table S5**). Similar to the ΔCjNC110 mutant, the previously described ncRNA CjNC140 was also found to be upregulated in the mutant condition during exponential growth only. During early stationary phase, six genes were found to be downregulated and six upregulated in the ΔluxS mutant strain when compared to wild-type. Of the differentially expressed genes, only one gene, CJSA_1350, a putative methyltransferase, was found to be downregulated in the ΔluxS mutant during both exponential and stationary phases. Three separate operons predicted by Rockhopper demonstrated multiple genes within the operon affected by the mutant condition (two operons upregulated, one operon downregulated). Analysis of functionality via the COG database revealed that multiple up- (*trpF, trpB*) and down- (CJSA_0620*, leuC*) regulated genes were present in the “Amino acid transport and metabolism” functional category; multiple upregulated genes were also included in the “Cell motility” (*flgE, flgG2, flgG*) functional category. Only two genes were differentially expressed in both the ΔCjNC110 and ΔluxS mutants, CJSA_0008 (upregulated during exponential phase) and CJSA_1107 (upregulated during stationary phase); both genes are annotated as hypothetical proteins at this time. Despite the critical role that LuxS plays in the AMC pathway, significant effects on the expression of other genes within the AMC pathway were not observed.

In the ΔCjNC110ΔluxS double knockout mutant, however, a large increase in both downregulated and upregulated genes when compared to wild-type was observed at both timepoints (summary **Table 3**, complete information **Table S6**). Sixty-one protein coding genes, seventeen tRNA genes, three known ncRNA genes (tmRNA, SRP, 6S) and seven newly predicted ncRNAs were downregulated during exponential phase compared to IA3902 wild-type. In addition, 47 protein coding genes were upregulated in exponential phase, along with the previously described ncRNA CjNC140, which was also observed to be upregulated in the ΔCjNC110 and ΔluxS single knockout mutants. During early stationary phase, 32 genes and two newly predicted ncRNAs were found to be downregulated, while 29 genes were upregulated in the mutant strain when compared to wild-type. Of the observed genes with differential expression, many were observed to show altered expression during both exponential and stationary phase when compared to wild-type; however, most were differentially expressed in the opposite direction between the two timepoints. Only two genes (*motB,* CJSA_1350) were downregulated in both conditions, while no genes were observed to be upregulated at both timepoints. Many of the differentially upregulated genes demonstrated in both background mutants, ΔCjNC110 and ΔluxS, were observed in the double knockout (**Figs. 2A** and **2B**), while fewer of the downregulated genes were shared (**Figs. 2C** and **2D**).

**FIG 2.**
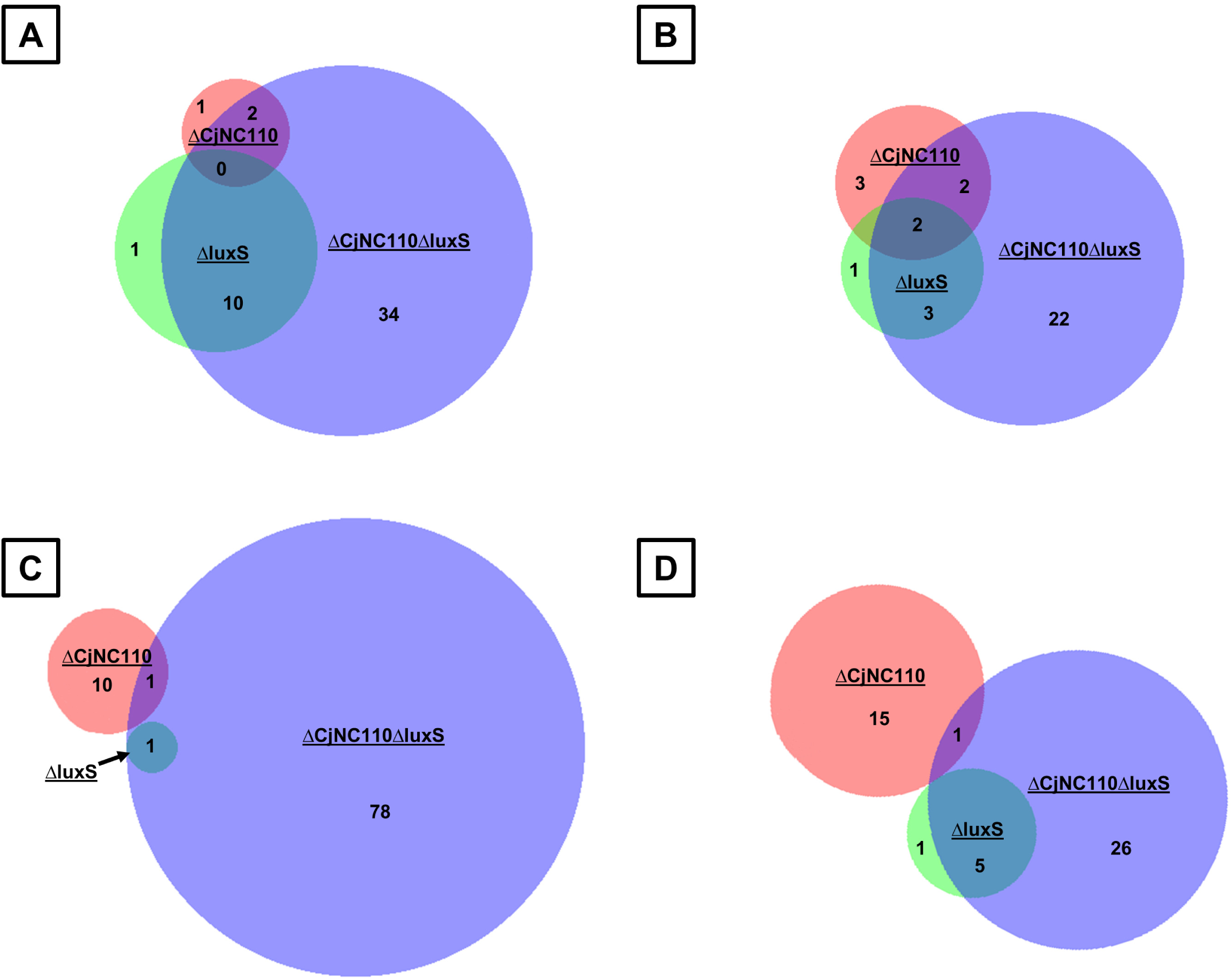
Venn diagram depicts overlap of shared up- and downregulated genes during the exponential and stationary growth phases of all 3 mutant strains. BioVenn was used to compare the lists of known genes identified by Rockhopper as upregulated in the RNAseq data in all 3 mutant strains during both exponential [A] and stationary [B] growth; downregulated genes were also compared for all 3 mutant strains during both exponential [C] and stationary [D] growth.

Of the differentially expressed genes affected by the ΔCjNC110ΔluxS double knockout mutation, 20 separate operons predicted by Rockhopper demonstrated multiple genes within the operon affected by the mutant condition (eight operons upregulated, four operons downregulated, and eight operons that showed opposing regulation at the different timepoints). **Figure S3** shows the number of genes in each COG category that were differentially expressed in the double mutant condition during either exponential growth. The categories most affected by these mutations were “Energy production and conversion,” “Amino acid transport and metabolism,” “Translation,” “Cell wall/membrane biogenesis,” “Cell motility,” and “Signal transduction mechanisms.” Using KEGG Pathways (31), a significant effect was observed in the flagellar assembly pathway in the double knockout mutant as compared to either mutant alone (**Figs. S4 and S5)**. Of particular interest, this affect was also observed to be altered based on growth phase, with decreased expression of σ^54^-associated genes noted during exponential phase, and increased expression noted during early stationary phase.

### Phenotypic evaluation of increased *luxS* expression unexpectedly reveals alteration of AI-2 transport in ΔCjNC110

Utilizing the potential targets identified in the RNAseq data for ΔCjNC110, phenotypic testing was next performed to begin to determine the physiologic role of CjNC110 in IA3902. Further investigation of a potential interaction with *luxS,* which was identified as demonstrating significantly increased expression in ΔCjNC110 during early stationary phase, was initially pursued due to the proximity of the two genes. LuxS activity is traditionally measured using the *Vibrio harveyi* bioluminescence activity assay as an approximation of AI-2 production (32). Using this assay as previously described, AI-2 levels were initially evaluated within the extracellular environment by collecting cell-free supernatant (E-CFS) at various time points during the growth of IA3902 wild-type and mutant strains. To collect these samples, a second set of growth curves independent of those utilized for RNAseq was performed in triplicate to include the ΔCjNC110c (**Fig. 3**). Similar to the first growth curves utilized for RNAseq, ANOVA again identified a statistically significant difference between the growth of the strains (*P <*0.05), however, the only significant difference was a decrease in the A_600_ of ΔCjNC110 after 27 hours, similar to the difference seen in the first set of growth curves. This defect was corrected in the ΔCjNC110c complement in the second set of growth curves, indicating that while the biological significance is unknown, this observation is repeatable and related to the loss of CjNC110.

**FIG 3.**
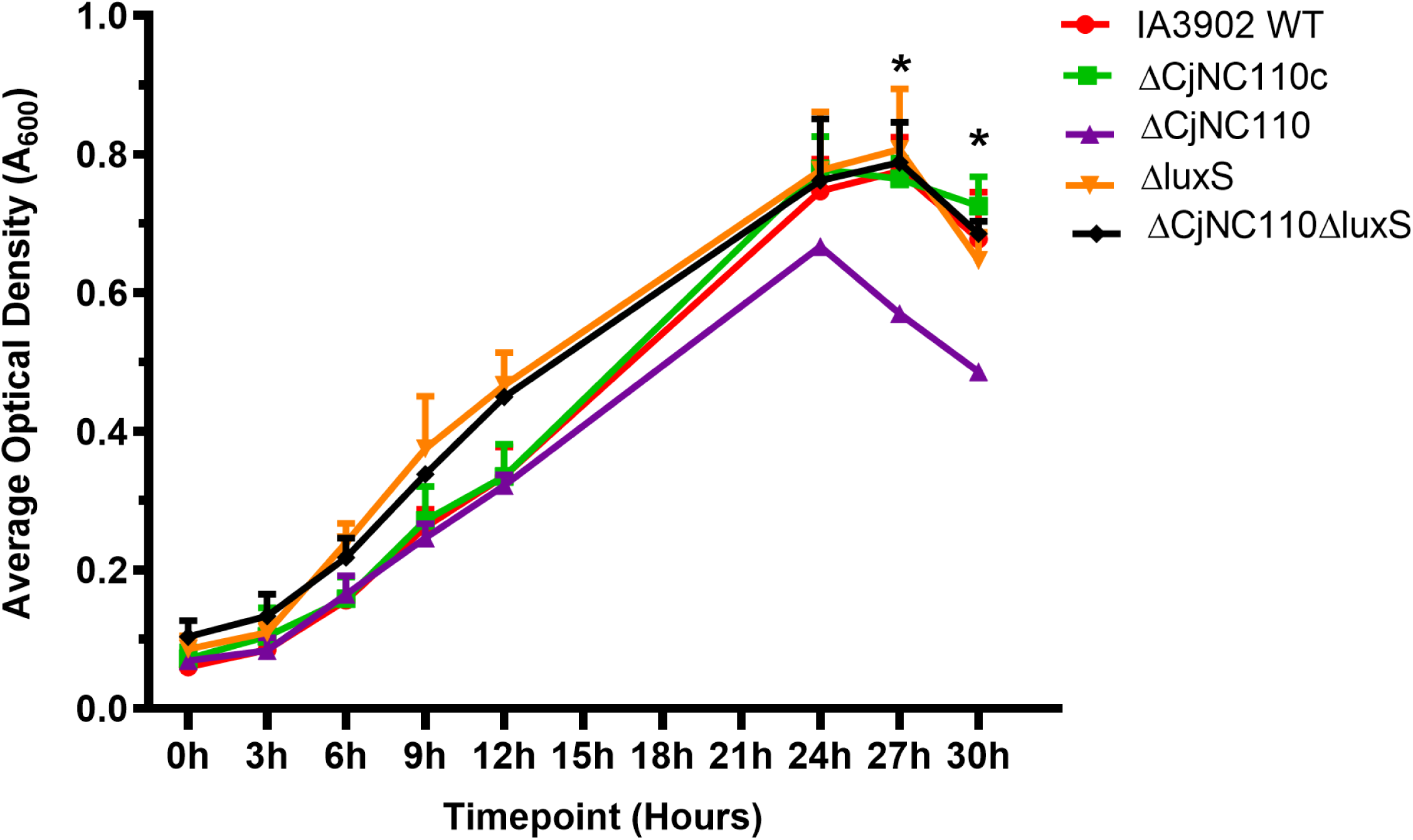
Shaking growth curve of wild-type IA3902 (WT) and isogenic mutants (mean ± SEM) confirms normal exponential growth. Results of four replicates (A_600_) of a shaking growth curve performed in 250 mL Erlenmeyer flasks under microaerophilic conditions in MH broth. Samples were pulled from this growth curve and used for qRT-PCR and AI-2 bioluminescence assays; for the results of the growth curve from which the RNAseq samples were taken, please see **Fig. S2A and S2B.** Significance is denoted by “*” above each timepoint.

As expected based on previous studies (8, 33), the results of the *V. harveyi* bioluminescence assay demonstrated that both the ΔluxS and ΔCjNC110ΔluxS mutant strains displayed no bioluminescence activity at any point during growth, indicating a complete lack of AI-2 production which is consistent with the absence of a functional LuxS protein (**Fig. 4A**). For the ΔCjNC110 mutant, however, bioluminescence was determined to be statistically significantly decreased when compared to wild-type at time points 6, 9 and 12 hours (*P <*0.05) of the growth curves, occurring during mid to late exponential phase and early stationary phase. This is in distinct contrast to the expected result based on RNAseq analysis which demonstrated increased *luxS* mRNA expression. Complementation of ΔCjNC110 completely restored extracellular AI-2 to wild-type levels, strongly suggesting that the observed defect in extracellular AI-2 was a true phenotype warranting further exploration.

**FIG 4.**
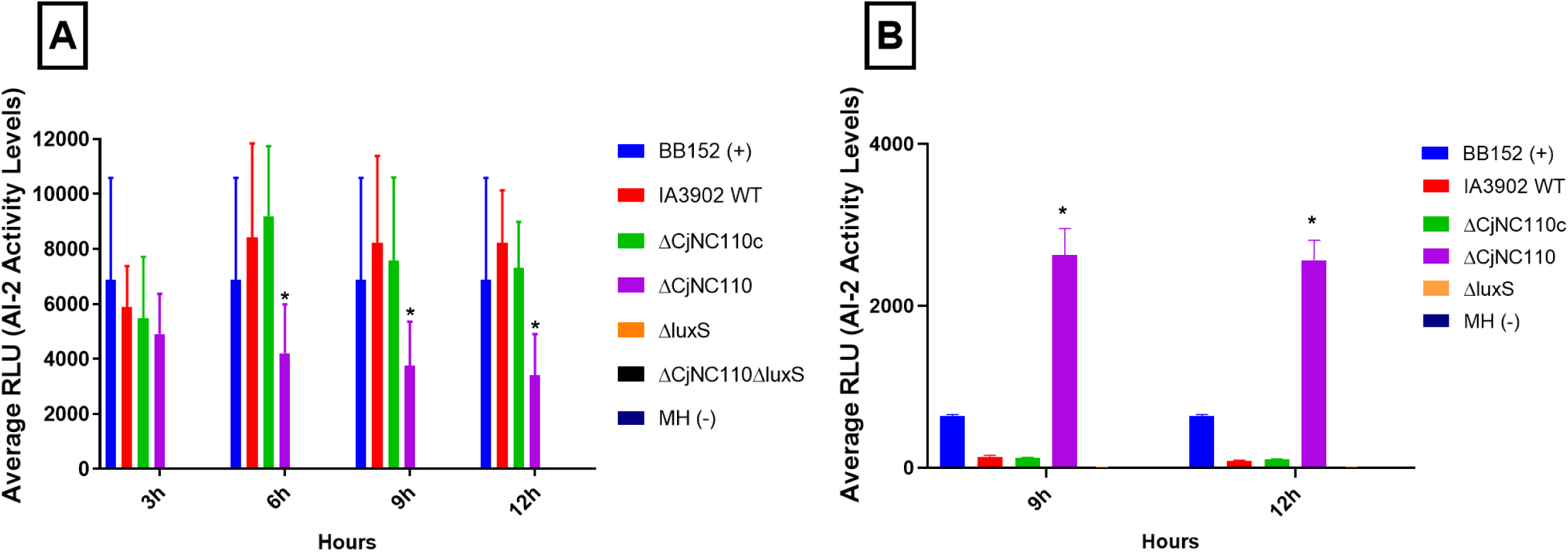
Bioluminescence activity as measured via the *Vibrio harveyi* bioassay over the course of growth demonstrates. (A) decreased extracellular AI-2 levels with (B) a concomitant increase in intracellular AI-2 in ΔCjNC110 when compared to wild-type IA3902 (WT) (mean ± SEM). The *V. harveyi* bioluminescence assay was used initially to detect AI-2 levels in the extracellular CFS to approximate LuxS activity (A). A second experiment was performed to investigate AI-2 levels in the intracellular CFS (B). Each bar represents the average RLU (AI-2 activity level) from three biological replicates consisting of three technical replicates each. Significance is denoted by “*” above each timepoint.

As the RNAseq results (i.e. – increased *luxS* gene expression) did not match the results of the AI-2 assay (i.e.- decreased extracellular AI-2), qRT-PCR was performed on samples collected at 12 hours from the same growth curve as the AI-2 assay using targets specific for the 3’ region of the *luxS* gene. **Fig. 5** demonstrates the expression levels of *luxS* in the ΔCjNC110 mutant compared to wild-type and confirms that increased *luxS* mRNA levels are indeed present during early stationary phase in the mutant, resulting in a fold change increase of 3.20 when compared to IA3902 wild-type (*P* <0.05). Additionally, ΔCjNC110c was tested to determine if complementation of the mutation returned *luxS* to wild-type levels; interestingly, ΔCjNC110c also demonstrated a fold change increase of 3.77 when compared to IA3902 wild-type (*P* <0.05).

**FIG 5.**
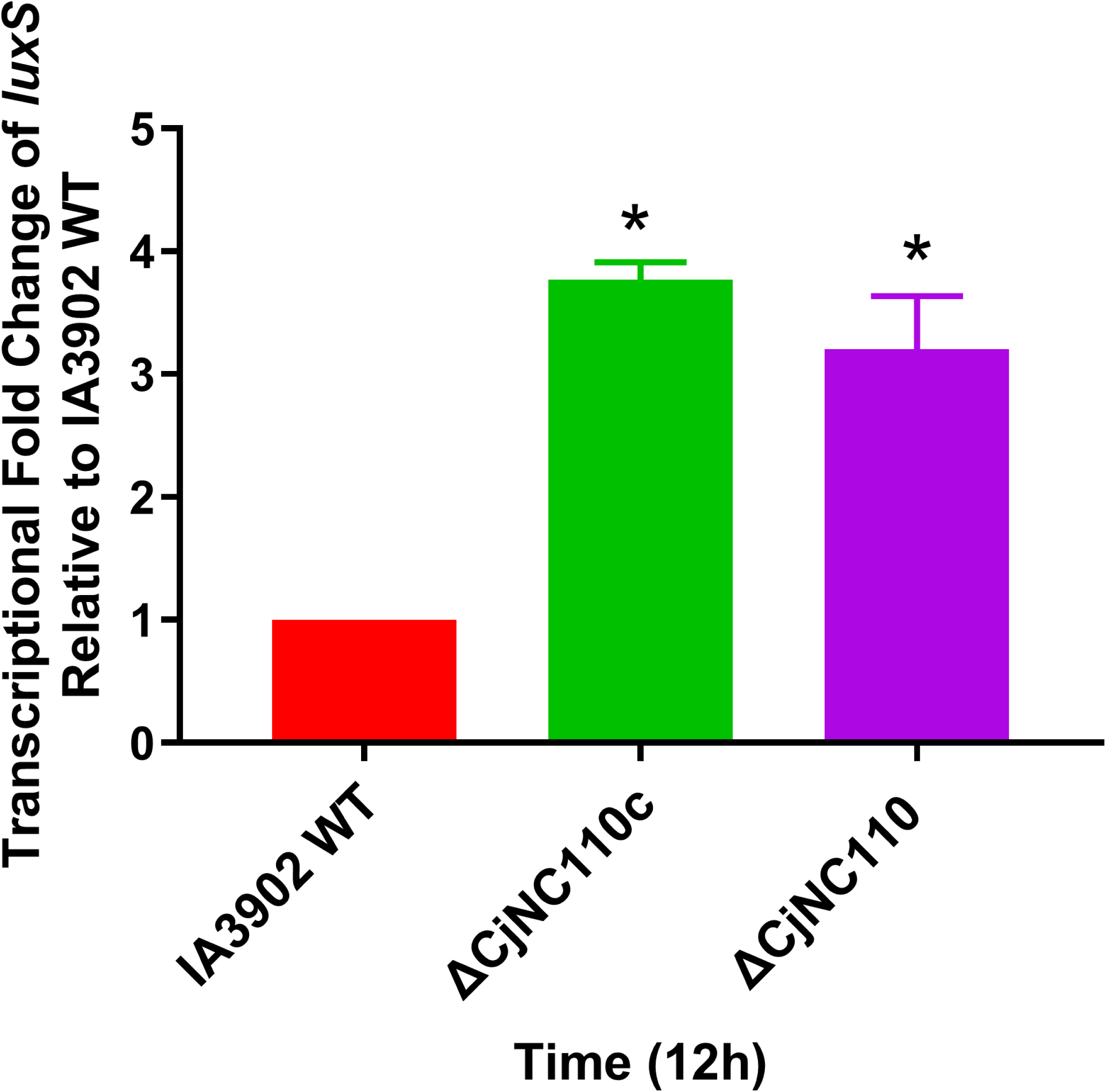
qRT-PCR confirms *luxS* is transcribed at higher levels in the ΔCjNC110 and ΔCjNC110c backgrounds when compared to IA3902 wild-type (WT) (mean ± SEM). *C. jejuni* cultures were collected at 12 hours for RNA conversion to cDNA for qRT-PCR to assess expression of mRNA *luxS*. Each bar represents the average fold change expression from three biological replicates. Significance is denoted by “*” above each timepoint.

Proteomic analysis using liquid chromatography with tandem mass spectrometry (LC-MS/MS) was then conducted to determine if the relative abundance of the LuxS protein was also increased in ΔCjNC110. Whole protein extract LC-MS/MS analysis resulted in a total of 262 identified proteins that were discovered in each biological group. Two-way ANOVA with FDR correction was used to identify statistically significant differences in protein abundance; these statistically significant proteins are listed in **Table 4**. In both ΔCjNC110 and ΔCjNC110c, the abundance of LuxS was noted to be statistically significantly increased (*P* <0.0001). Hierarchical heat maps were also constructed to further investigate protein expression patterns when comparing IA3902 wild-type to ΔCjNC110 and ΔCjNC110c. The top 20 hierarchical clustered proteins are illustrated for both ΔCjNC110 and ΔCjNC110c compared to wild-type IA3902 in **Fig S6**; LuxS was also identified via this approach as being differentially expressed in both ΔCjNC110 and ΔCjNC110c. Therefore, the proteomics data confirmed the RNAseq and qRT-PCR observations of significantly increased expression of LuxS when comparing wild-type to both ΔCjNC110 and ΔCjNC110c.

**TABLE 4:** Significant differentially expressed proteins identified from proteomic analysis ΔCjNC110 and ΔCjNC110c versus IA3902 wild-type

| Protein ID | Geometric Mean<br>(Abundance) |  | Fold<br>Change<br>vs WT | Q-value* |
| --- | --- | --- | --- | --- |
|  | WT | Mutant |  |  |
| <u>**ΔCjNC110**</u> |  |  |  |  |
| CJSA_0067 - iron-sulfur protein | 23726566.41 | 851708.3688 | -27.9 | <0.0001 |
| CJSA_0489 - rhodanese-like domain-containing | 35962749.77 | 2965820.801 | -12.1 | 0.0001 |
| ThiJ - thiazole monophosphate synthesis protein | 1589344.014 | 31307392.07 | 19.7 | <0.0001 |
| FrdA - fumarate reductase flavoprotein subunit | 3178688.029 | 280958.9826 | -11.3 | 0.0002 |
| **LuxS - S-ribosylhomocysteine lyase** | **3913424.009** | **50859008.46** | **13.0** | **<0.0001** |
| <u>**ΔCjNC110c**</u> |  |  |  |  |
| CJSA_0067 - iron-sulfur protein | 23726566.41 | 1123835.93 | -21.1 | <0.0001 |
| CJSA_0489 - rhodanese-like domain-containing | 35962749.77 | 2767208.654 | -13.0 | 0.0001 |
| CJSA_0166 – lipoprotein | 25429504.23 | 2581896.969 | -9.8 | 0.0006 |
| RplN - 50S ribosomal protein L14 | 10327587.87 | 1048576 | -9.8 | 0.0007 |
| RplU - 50S ribosomal protein L21 | 15653696.03 | 3178688.029 | -4.9 | 0.0007 |
| RecJ - single-stranded-DNA-specific exonuclease | 2408995.053 | 19271960.42 | 8.0 | 0.002 |
| AtpF - ATP synthase subunit B | 2965820.801 | 27254667.8 | 9.2 | 0.001 |
| **LuxS - S-ribosylhomocysteine lyase** | **3913424.009** | **54509335.6** | **13.9** | **<0.0001** |
| CJSA_0926 - lipoprotein | 1703416.738 | 23726566.41 | 13.9 | 0.0001 |

Geometric mean calculated by back-transforming log2 transformation value
\*Two-way ANOVA FDR set at 0.05
WT = wild-type

As decreased LuxS abundance was not the cause of the observed decrease in extracellular AI-2, several additional hypotheses were proposed, including: 1) decreased activity of the LuxS protein leading to decreased AI-2 production, 2) increased degradation of AI-2, or 3) altered AI-2 transport into or out of the extracellular environment. To assess for altered AI-2 transport, the traditional *V. harveyi* bioluminescence activity assay was modified as described in the Materials and Methods to assess for intracellular AI-2 levels using intracellular cell free supernatant (I-CFS) obtained from the cell pellet of the same growth curve sample as the E-CFS. Results of this assay are shown in **Fig. 4B** and clearly demonstrate that intracellular AI-2 levels increased to statistically significant (*P <*0.05) levels in the ΔCjNC110 mutant when compared to IA3902 wild-type at both timepoints tested. Again, complementation of ΔCjNC110 corrected the phenotypic alteration of increased intracellular AI-2 presence to wild-type. When taken together, these results indicate that the observed decrease in extracellular AI-2 activity is not due to impaired activity of LuxS, but instead strongly suggests that transport of AI-2 across the cell membrane is altered in the ΔCjNC110 mutant leading to decreased extracellular and increased intracellular AI-2 accumulation.

### Mutation of CjNC110 significantly increases motility and hydrogen peroxide sensitivity while decreasing autoagglutination in IA3902, opposite of the effects seen with mutation of *luxS*

To further investigate the role of CjNC110 in IA3902, several additional phenotypes suggested by the RNAseq results were assessed, including motility, hydrogen peroxide sensitivity, and autoagglutination. Loss of c*etA* and *cetB* has been previously shown to affect energy taxis and motility (34), therefore, the motility of the mutant strains in semi-solid agar was initially compared to wild-type IA3902 at 30 hours post-inoculation. The results of this assay confirmed that all isolates were highly motile, and statistical analysis via one-way ANOVA indicated that there was a significant difference between strains (*P* <0.0001) (**Fig. 6**). Further post-hoc analysis via Tukey’s multiple comparisons test did not reach statistical significance (*P* >0.05) when comparing ΔCjNC110 to wild-type despite motility for the ΔCjNC110 strain observed to be consistently increased above the wild-type phenotype in all biological replicates performed. Results did reach statistical significance (*P* <0.05), however, when comparing wild-type to both the ΔCjNC110ΔluxS and ΔluxS mutants which displayed statistically significant decreased motility; this result is consistent with previous reports of decreased motility of *luxS* mutants in the IA3902 background (8). As many of the strains had migrated to the edge of the plate by 30 hours, a second set of experiments was performed with measurements taken at 24 hours post-inoculation. When measured at 24 hours, motility for the ΔCjNC110 strain did demonstrate a statistically significant increase (*P* <0.05) when compared to all other strains tested, and motility of both the ΔCjNC110ΔluxS and ΔluxS mutants was again noted to be significantly decreased **(Fig. 6)**. Following complementation of CjNC110, no significant difference was observed when compared to IA3902 wild-type in either experiment, confirming that complementation of CjNC110 quantitatively restored the phenotypic change back to wild-type levels. These results indicate an increased motility of ΔCjNC110, which differs from ΔluxS which displays decreased motility in IA3902.

**FIG 6.**
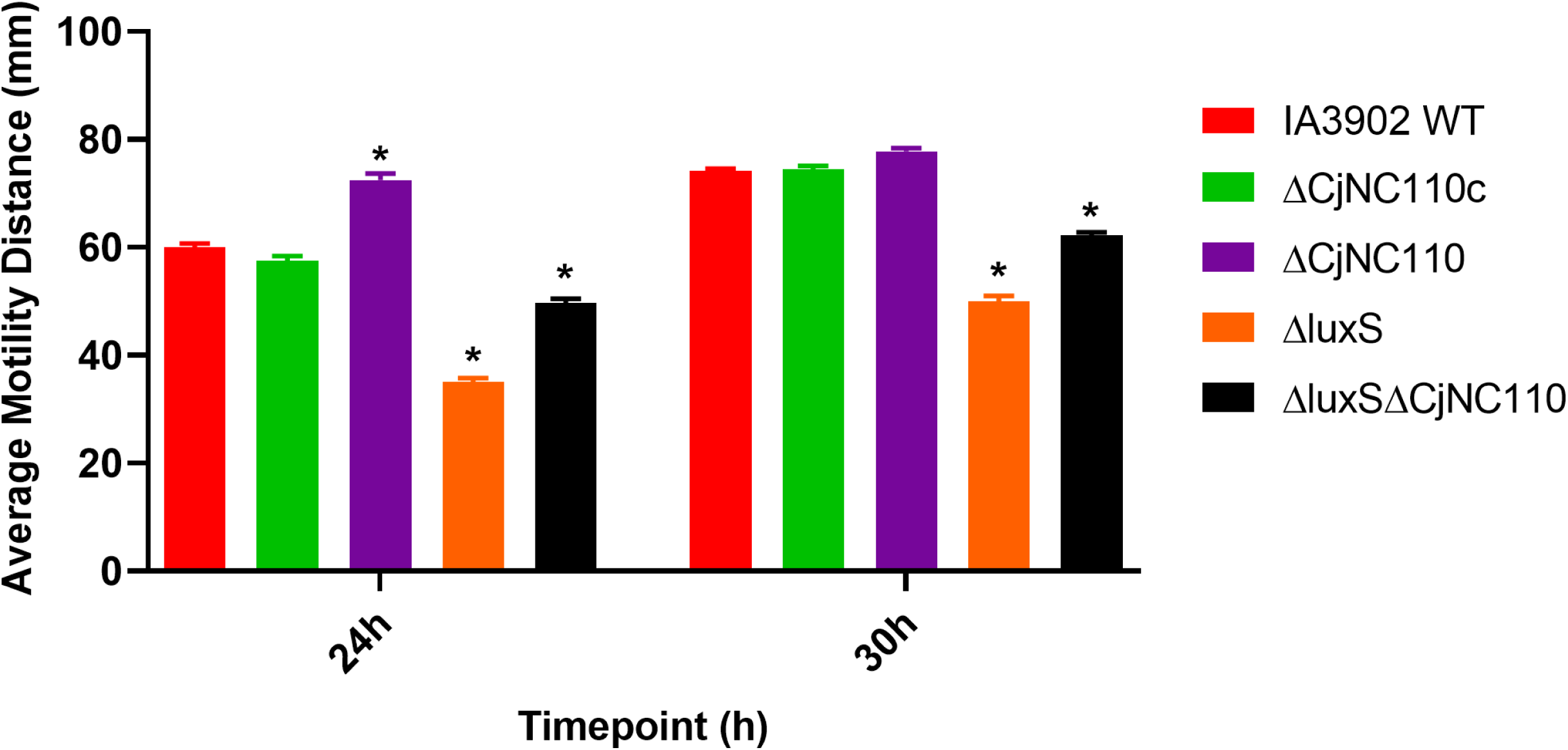
Motility assays confirm isogenic mutants remain motile, with ΔCjNC110 motility increased and ΔluxS motility decreased over IA3902 wild-type (WT) (mean± SEM). Swarming motility in semi-solid agar was assessed via measurement in mm of the outermost reach of growth at both 30 hours and 24 hrs. Each bar represents the average motility of six biological replicates from six independent studies. Significance is denoted by “*” above each bar when comparing wild-type to other respective strains at each independent timepoint.

Based on the potential regulatory interaction identified in our RNAseq data of CjNC110 with *tpx,* which has been previously demonstrated to encode a dedicated hydrogen peroxide reductase in *Campylobacter* sp., a hydrogen peroxide (H_2_0_2_) disk inhibition assay was performed as previously described (35). **Fig. 7** demonstrates the results of this assay which indicates that the lack of CjNC110 significantly impacts H_2_0_2_ sensitivity when compared to wild-type as increased sensitivity was observed in both CjNC110 mutant backgrounds (ΔCjNC110 and ΔCjNC110ΔluxS) (*P* <0.05). Complementation of CjNC110 led to a significantly decreased sensitivity to H_2_0_2_ compared to wild-type, indicating that the potential overexpression of CjNC110 in ΔCjNC110c may lead to overcorrection of the phenotypic change (*P* <0.05). There was no significant difference (*P* >0.05) noted between wild-type and ΔluxS, which is consistent with previous studies of *luxS* mutation in other strains of *C. jejuni* such as NCTC 11168 (36). These results indicate that CjNC110 may play a key role in sensing and responding to oxidative stress in the environment for IA3902, and again represent a differing phenotype between ΔCjNC110 and ΔluxS.

**FIG 7.**
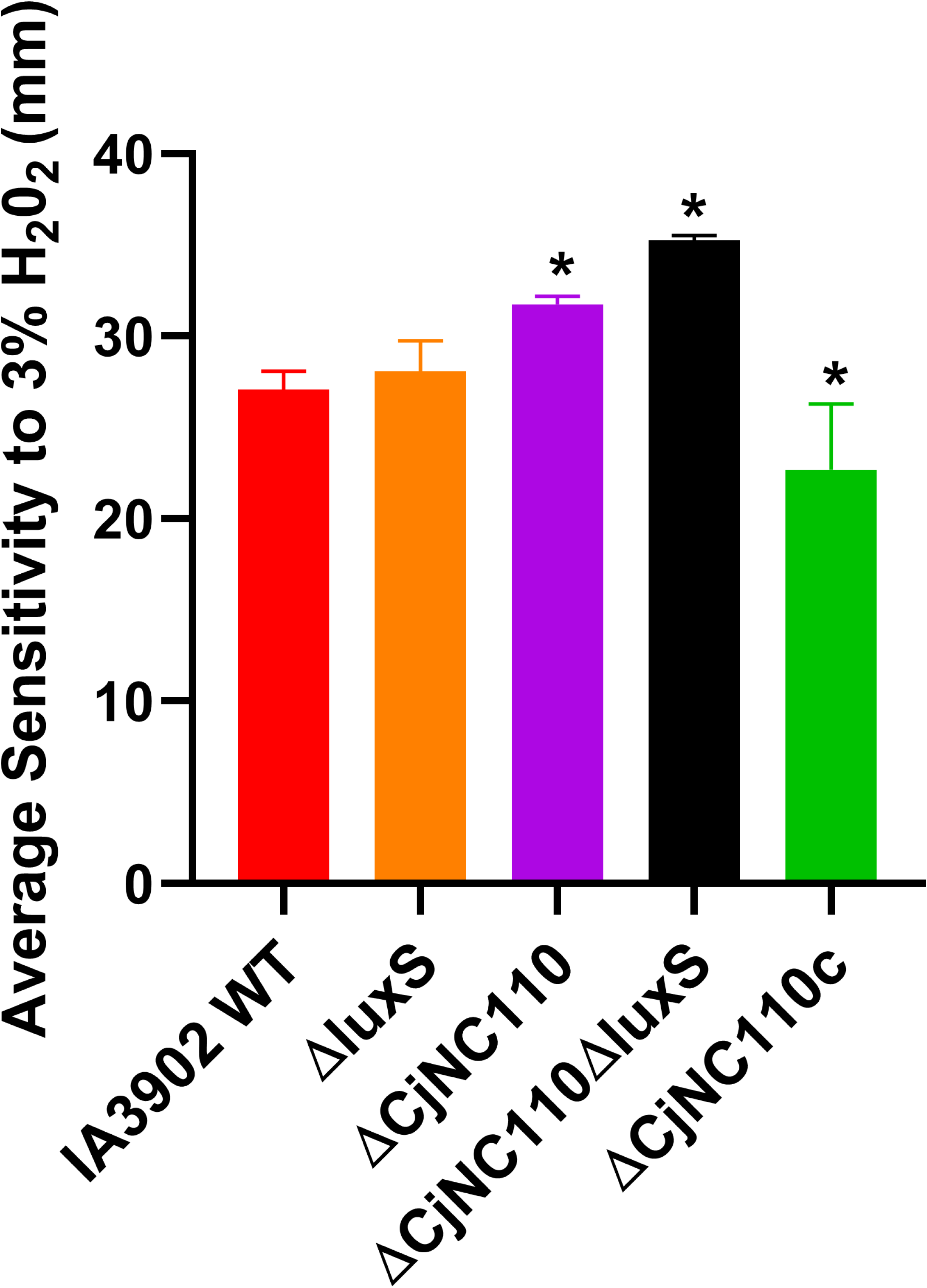
ΔCjNC110 displays increased hydrogen peroxide (H_2_0_2_) sensitivity *in vitro* when compared to wild-type IA3902 (WT) (mean ±SEM). Cell cultures normalized to an A_600_ of 1.0 were inoculated into melted MH agar then incubated with a 6 mm disk soaked in 3% H_2_0_2_ placed at the center of the solidified MH agar plate; after 24 hours the zone of sensitivity to H_2_0_2_ was measured in mm. Each bar represents the average of three biological replicates from four independent studies. Significance is denoted by “*” above each bar when comparing wild-type to other respective strains at each independent timepoint.

Differential expression of several genes associated with flagellar modification [*pseA, ptmA, ptmB, neuB2,* and CJSA_1261] was also observed in ΔCjNC110; these genes have been previously shown to affect the autoagglutination ability of *C. jejuni* (37–39). Therefore, autoagglutination was also assessed as previously described (40) to determine if the observed gene expression changes in ΔCjNC110 led to a phenotypically observable alteration. Autoagglutination activity was measured at both 25°C and 37°C following 24 hours of incubation with an increase in optical density correlating to decreased autoagglutination ability, and a decrease in optical density correlating to increased autoagglutination ability. A statistically significant difference (*p <*0.0001) between strains at both 25°C and 37°C was noted based on initial analysis via one-way ANOVA (**Fig. 8**). When compared to IA3902 wild-type, ΔCjNC110 exhibited statistically significant decreased autoagglutination activity (*P <*0.05) at both temperatures, while autoagglutination activity for ΔluxS and ΔCjNC110ΔluxS was noted to be increased at a statistically significant level (*P <*0.05), again at both temperatures. Complementation of CjNC110 restored the autoagglutination ability of ΔCjNC110 to wild-type levels. These results indicate that *luxS* and CjNC110 function to influence autoagglutination activity in opposing directions; interestingly, when mutation of both genes was combined, the results favored the ΔluxS mutation but did return closer to wild-type levels.

**FIG 8.**
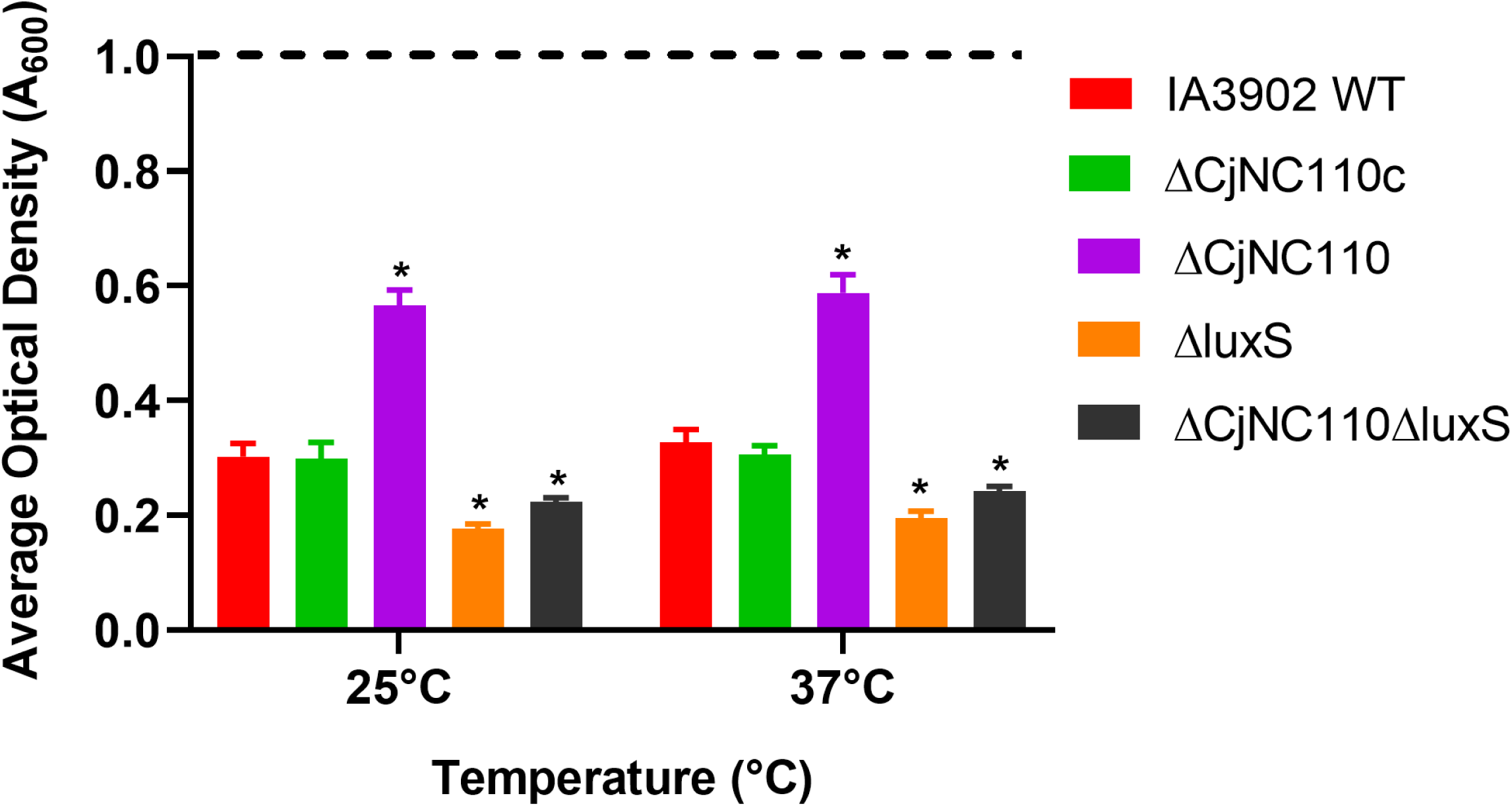
Autoagglutination ability of IA3902 is decreased in ΔCjNC110 and increased in ΔluxS when compared to wild-type IA3902 (WT) (mean± SEM). Autoagglutination as determined by optical density (A_600_) was measured following a 24 hour incubation at either 25°C or 37°C. Each bar represents the average autoagglutination of 3 biological replicates consisting of 4 technical replicates during each independent study. An increase in optical density correlates to decreased autoagglutination ability, and a decrease in optical density correlates to increased autoagglutination ability. Significance is denoted by “*” above each bar when comparing wild-type to other respective strains at each independent timepoint.

### Sustained colonization of chickens is impaired in ΔCjNC110 and absent in mutants ΔCjNC110ΔluxS and ΔluxS, but is restored to wild-type colonization levels in ΔCjNC110c

The ability to colonize the host is the critical first step in establishment of infection by *Campylobacter* sp. Previous studies have demonstrated a loss of colonization ability of ΔluxS in IA3902 (8), therefore, chicken colonization was utilized to determine if a similar phenotype was present in either ΔCjNC110 or ΔCjNC110ΔluxS. Three-day old chicks were orally inoculated with approximately 200 µl of 10^7^ CFU per strain (wild-type, ΔCjNC110, ΔluxS, ΔCjNC110ΔluxS and ΔCjNC110c). Once weekly for 3 weeks following inoculation, 6 chicks were humanely euthanized and cecel contents collected for analysis of colonization levels via plating of serial dilutions and colony counts. For ΔCjNC110, while colonization was still present at all timepoints, a statistically significant decrease in colonization was noted compared to wild-type on days post-inoculation (DPI) 5, 12, and 19 (*P <*0.05); complementation of the ΔCjNC110 mutant restored colonization to wild-type levels at all timepoints analyzed (**Fig. 9**). Similar to previous studies, while some colonization, albeit significantly decreased, was noted for ΔluxS at DPI 5, no colonization was observed by DPI 12 and 19 (*P <*0.05). For the ΔCjNC110ΔluxS mutant, normal colonization levels compared to wild-type were observed on DPI 5, however, by DPI 12 and 19, no colonization of ΔCjNC110ΔluxS was detectable (*P <*0.05). These results indicate that the presence of CjNC110 is necessary for optimal colonization, while *luxS* is required for establishing sustained colonization of IA3902 in chickens.

**FIG 9.**
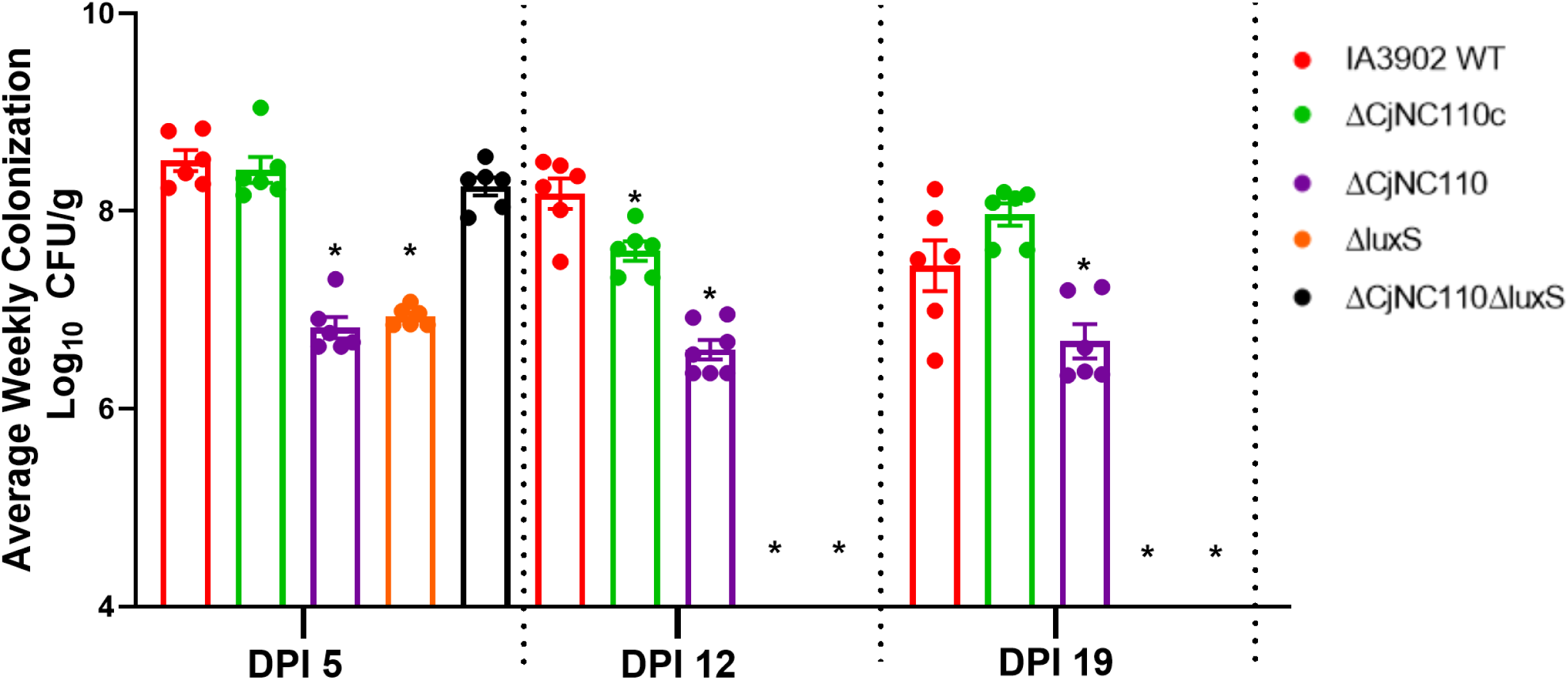
Sustained colonization in chickens is impaired in ΔCjNC110 and absent in ΔluxS and ΔCjNC110ΔluxS mutants compared to wild-type (mean ± SEM). Results of a single study investigating the colonization ability of mutants strains compared to wild-type (WT) IA3902 are reported as log_10_ CFU/mL of ceca content for each sampling day [DPI 5 (Week 1), DPI 12 (Week 2), and DPI 19 (Week 3)]. Each bar represents the average colonization levels for each biological group with weekly technical replicates using a minimum of 6 birds. Significance is denoted by “*” above each timepoint.

## DISCUSSION

In other species of bacteria that harbor known RNA chaperones such as Hfq, methods of identification of cognate mRNA binding partners of ncRNAs are available via several straightforward approaches [recently reviewed in (41)]. As *Campylobacter* sp. lack all known RNA chaperones, identification of targets via these methods is not feasible in these pathogens; previous attempts at computational predictions have also failed (18). However, recent work in *Campylobacter* has demonstrated that RNAseq can be useful for analyzing transcriptomic differences between wild-type and mutant strains following inactivation of protein-coding genes (42, 43). Indeed, over the past several years, RNAseq has become the method of choice to analyze the transcriptomes of bacteria under various conditions as it allows evaluation of the entire transcriptome rather than only previously annotated regions as with older technology such as microarrays (44–46). In theory, RNAseq should also be able to be utilized for the same purpose of discovering the global effects of inactivation of ncRNAs. Small ncRNAs have been shown to be able to stabilize mRNA transcripts by multiple mechanisms (47), which should lead to increased levels of transcript availability and identification of significant differences using RNAseq. Some interactions with ncRNAs have also been shown to increase transcript turnover by targeting transcripts for RNase degradation or exposing RNase cleavage sites (47), which should lead to decreased levels of transcripts available for identification via RNAseq. Thus, assessment of differential gene expression via RNAseq should be useful for uncovering potential interactions of ncRNAs with mRNA targets when that interaction directly leads to altered levels of mRNA transcripts in the cell; target interactions that do not lead to direct changes in mRNA transcript levels but instead affect the translation efficiency of the target mRNA or are the result of direct protein-ncRNA interactions cannot be determined via this approach. While our differential gene expression results do not guarantee a direct mRNA-ncRNA interaction or provide a mechanism for that interaction, the results of the work presented here clearly indicate that RNAseq can be a useful tool for investigating the role of ncRNAs in post-transcriptional regulation of gene expression in bacteria, and can successfully be used to inform the design of phenotypic studies to begin to elucidate the function of ncRNAs in pathogenic bacteria who lack RNA chaperones.

One particular area of interest from the onset of our work was the potential for an interaction between CjNC110 and *luxS.* Our results related to a potential interaction between *luxS* and CjNC110 were surprising in that despite increased LuxS expression, extracellular AI-2 levels were decreased while intracellular AI-2 increased; this ultimately suggested that CjNC110 plays a key role in the transport of AI-2 between the intra- and extracellular environment of IA3902. In *E. coli,* the YdgG (TqsA) protein, part of the AI-2 exporter superfamily, has been described to export AI-2 molecules (48); additional *in vitro* studies assessing the membrane permeability of AI-2 also demonstrate that it is hydrophilic with low affinity towards lipids, which strongly suggests the need for a transport system (49). Previous BlastP analysis has shown potential homologues of the YdgG protein exist within the *Campylobacterales* family, but none directly within the *C. jejuni* species (50). Within our proteomics data, several periplasmic and lipoproteins which could be located in the cytoplasmic membrane were identified to exhibit decreased expression in ΔCjNC110 that was corrected to wild-type levels in ΔCjNC110c. The function of many of these proteins has yet to be elucidated in *Campylobacter,* therefore, future work assessing whether these proteins may be regulated by CjNC110 and could serve as an AI-2 exporter in *C. jejuni* is warranted. In other species of bacteria such as *E. coli*, uptake of AI-2 from the extracellular environment has also been shown to occur and is mediated by an active LSR system (51). Comparative genomics has again failed to identify a homologous AI-2 uptake system in *C. jejuni*, however, a number of ABC-type transporters whose function remains to be elucidated do exist in *Campylobacter* sp. (28). Several studies evaluating the concentration of AI-2 over time following addition of exogenous AI-2 to cell culture have attempted to determine whether or not AI-2 is able to be internalized in *C. jejuni* with mixed results (52, 53). Thus, further research into an AI-2 uptake system in *C. jejuni* and the role that CjNC110 may play in regulation of that uptake may also be warranted.

Based on the AI-2 results, the increased expression of LuxS in ΔCjNC110 and failure of complementation of *luxS* gene expression and LuxS protein abundance in ΔCjNC110c was unexpected, particularly as all other phenotypes tested, including AI-2 detection intra- and extracellularly, fully complemented to wild-type. The reason for the increased *luxS* expression and LuxS protein abundance in ΔCjNC110 and ΔCjNC110c is currently unknown but one reasonable hypothesis is that this increased expression may be related to additional unknown effects of CjNC110 on the AMC leading to feedback regulation of *luxS* expression. While regulation of the AMC in other Proteobacteria has been studied in detail, the regulatory pathways of the AMC are currently unknown in *Campylobacter* as this genus lacks all of the known transcription factors and riboswitches typically involved regulation (54). As CjNC110 is located immediately downstream of *luxS,* en bloc removal of the CjNC110 coding sequence could also have led to disruption of normal termination of the *luxS* transcript, thus altering the natural mRNA stability; this effect would have been maintained in the complement as well. Further investigation to determine the location of a terminator via the WebGesTer database failed to identify a predicted terminator sequence for *luxS* (http://pallab.serc.iisc.ernet.in/gester/) (55). Therefore, additional work is needed to determine which hypothesis might explain the increase of LuxS in both ΔCjNC110 and ΔCjNC110c. Regardless of the cause, our data clearly demonstrates that the increased *luxS* expression and LuxS protein abundance in ΔCjNC110 is unrelated to the observed changes in extracellular and intracellular AI-2 concentrations which fully complemented back to wild-type levels in ΔCjNC110c.

Several other important phenotypes were also affected by mutation of CjNC110 based on the RNAseq data, including motility, autoagglutination, hydrogen peroxide sensitivity, and chicken colonization. Motility is considered critical for the *in vivo* virulence of *C. jejuni* and requires a functional flagellar apparatus (56, 57); however, this alone is not sufficient for normal motility to be present (58). In our RNAseq data, the *cetAB* operon (*<u>C</u>ampylobacter* <u>e</u>nergy taxis proteins A and B), which is known to mediate the energy taxis response in *Campylobacter* (34), was statistically significantly upregulated in the ΔCjNC110 mutant when compared to wild-type during the exponential phase of growth; expression of *cetAB* was increased during stationary phase as well, however, did not reach the level of statistical significance. Defects in both *cetA* and *cetB* have been shown to lead to altered motility phenotypes, particularly in response to migration towards critical factors in *Campylobacter* metabolism such as sodium pyruvate and fumarate (58). Thus, the increase in *cetAB* correlates well to the statistically significant increase in motility found between the ΔCjNC110 mutant and wild-type when motility was assessed at 24 hours. Investigation of expression of *cetA* and *cetB* has shown that levels of the gene products are unaffected by mutation of sigma factors σ^54^ or σ^28^, indicating that transcription of the *cetAB* operon is likely controlled by σ^70^ or another yet unknown transcription factor (34). Based on this information, it seems plausible that CjNC110 might normally act as a repressor of the CetAB energy taxis system. Our proteomics data failed to detect the CetA and CetB proteins, therefore, we could not validate that increased transcriptional expression of *cetA* and *cetB* resulted in increased protein expression levels. Many of the proteins identified in our study exhibited extremely high expression levels, which, without fractioning, prevented proteins of lesser abundance from being detected. Whole cell proteomic analysis with fractioning and time-course analysis would be required to reveal the entire proteomic network regulated by CjNC110 and validate expression patterns of proteins of lesser abundance than the 262 proteins that were identified in our study. Further work to determine if a direct interaction between CjNC110 and the *cetAB* operon exists and leads to increased expression of the CetA and CetB proteins is warranted.

No statistically significant differences in the expression of genes associated with the flagellar apparatus were noted in the transcriptome of ΔCjNC110; this may help to explain why an increase in energy taxis would allow for an increase in observed motility with the assumption that normally functioning flagella are present. The presence of normal flagella has also been associated with autoagglutination ability and is considered necessary but not sufficient for this important trait (58). Previous studies in *C. jejuni* have demonstrated that in addition to the presence of normal flagella, interactions between modifications on adjacent flagellar filaments, particularly those provided by O-linked protein glycosylation, are required for normal autoagglutination ability (59). In the present study, the ΔCjNC110 mutant exhibited decreased autoagglutination when compared to wild-type at both 25°C and 37°C. Several genes that have been associated with flagellar glycosylation in other strains of *C. jejuni* were downregulated in our RNAseq data in the ΔCjNC110 mutant when compared to wild-type during both exponential and stationary growth phases. The O-linked protein glycosylation system which is responsible for flagellar glycosylation has been extensively studied in only one strain of *C. jejuni* (81-176) and has not been studied to date in IA3902 (38); this system is one of the most diverse genomic loci in *Campylobacter* spp. and also represents one of the key areas of variation between IA3902 and NCTC 11168 (5). Further genetic analysis of the flagellar modification region of the IA3902 genome reveals the presence of genes associated with both of the previously identified pseudaminic acid and legionaminic acid flagellar glycosylation pathways known to exist in *Campylobacter* sp.; this analysis and the subsequent gene expression data comparison of ΔCjNC110 to wild-type is presented in **Table S7**. The pseudaminic acid pathways (encoded by the *pseA-I* genes which are fully annotated in IA3902) lead to flagellar glycosylation with either pseudaminic acid (PseAc) (Pse5Ac7Ac), which is the major glycan found on *Campylobacter* flagella, or derivatives such as PseAm (Pse5Am7Ac) which has an acetamidino substitution (60). In our RNAseq dataset, *pseA* was significantly decreased in expression during exponential growth, and also decreased without reaching significance during stationary phase. Mutation of *pseA* has previously been shown to lead to flagellar modification with only PseAc, and not PseAm (38, 60); *pseA* has also been shown to be required for optimal adherence and invasion of human epithelial cells and virulence in a ferret model (59).

The legionaminic acid (LegAm) flagellar glycosylation pathway, encoded by the *ptmA-H* genes, has been less extensively studied in *C. jejuni*, in part because it is not present in 81-176; in fact, much of the work that has been done in *Campylobacter* spp. related to this pathway has been in *C. coli* (37, 39). NCTC 11168 does contain homologs to all of the *ptm* genes (Cj1324-Cj1332) albeit with different gene nomenclature (*leg* or *neu*) often used to describe the genes, and mutation of these genes in NCTC 11168 has been shown to decrease autoagglutination ability as well as reduce but not eliminate the ability to colonize chickens (61). While not currently annotated as such, our further analysis has identified that in IA3902, gene locus CJSA_1261-1268 represent homologs to all of the *ptm* genes present in NCTC 11168 (**Table S7)**. Based on this data, it is reasonable to suggest that IA3902 also harbors an active LegAm flagellar glycosylation pathway. Several genes related to legionaminic acid flagellar glycosylation were identified in our RNAseq dataset as being differentially expressed in the CjNC110 mutant during both stages of growth, which may help explain altered autoagglutination levels of ΔCjNC110. In particular, *ptmA* and *neuB2* (homologous to *ptmC)* were both observed to be statistically significantly decreased at both exponential and stationary phase, while *ptmB* was statistically significantly decreased at stationary phase and decreased but not reaching statistical significance during exponential phase. Mutation of the *ptmA-G* genes in *C. coli* results in the loss of all LegAm derivatives and a conversion to glycosylation with PseAc only; these mutants still retain a normal flagellar apparatus and motility (37, 39, 62). In our RNAseq data, CJSA_1261 (homologous to *ptmG*) was the sole flagellar modification gene that demonstrated a statistically significant increase in expression during both growth phases; *ptmG* has been shown to be involved in conversion of LegAm to various derivatives (Leg5AM7Ac and Leg5AmNMe7Ac) (37). Alterations in *ptmG* levels in the CjNC110 mutant are of particular interest as it has been recently shown to be also regulated by another small non-coding RNA pair in *C. jejuni*, CjNC180/CjNC190 in NCTC 11168 (27). While CjNC180/CjNC190 were not detected in this dataset by Rockhopper, these ncRNAs are present in IA3902 and CjNC180 has previously been demonstrated to be significantly upregulated in the *in vivo* host gallbladder environment in IA3902 (24). When taken together, this data suggests that CjNC110 may play a key role in modulation of flagellar glycosylation in IA3902, thus directly affecting autoagglutination ability. Other potential causes for decreased autoagglutination ability include alteration of the flagellar filament. While not identified via any other approach, the FlaA protein was demonstrated to be decreased via hierarchal heat mapping in our proteomics data in the CjNC110 mutant with a return to wild-type levels in the complement (**Fig. S6**); if this change corresponds to an alteration of the flagellar structure, it could in theory also explain the observed decrease in autoagglutination ability. Thus, further work including determination of the normal flagellar glycosylation of IA3902 and assessment for changes in both the flagellar filament structure and glycosylation of the ΔCjNC110 mutant when compared to the IA3902 wild-type is warranted.

The RNAseq data also demonstrated that in the ΔCjNC110 background *tpx* was significantly downregulated during the exponential phase of growth, prompting phenotypic assessment of hydrogen peroxide sensitivity. Thiol peroxidase (Tpx) has been previously shown to function as a hydrogen peroxide reductase in NCTC 11168, however, in that strain, Tpx-defective mutants were not shown to be sufficient to cause alteration of hydrogen peroxide sensitivity via disc diffusion and required a second mutation of the Bcp gene to observe increased sensitivity (35). IA3902 also harbors a Bcp homolog that is 96% homologous to NCTC 11168; this gene did not demonstrate altered expression via RNAseq analysis, and functional evaluation of this gene in IA3902 has not been performed to date. While our results clearly indicate that the loss of CjNC110 increases hydrogen peroxide sensitivity, and that the expression of *tpx* in the mutant decreased, it is not possible to definitively state that the observed phenotype is due solely to an interaction of CjNC110 with *tpx* alone. Further analysis of potential alterations of CjNC110 with additional known oxidative stress genes in the RNAseq data did indicate that *sodB,* which has been previously identified to contribute to hydrogen peroxide sensitivity in *C. jejuni* (63), demonstrated a *Q*-value that reached the threshold for significance (*Q* = 0.004), however, the fold change just missed the cutoff for significance at −1.4. Thus, further investigation for the potential interaction of CjNC110 with both *tpx* and additional genes that control the oxidative stress response in *C. jejuni* such as *sodB* is also warranted.

When taken together, the results of the *in vitro* phenotypic studies, which identified several key phenotypes affected by the loss of CjNC110, strongly suggested that mutation of CjNC110 may interfere with normal colonization of IA3902. Indeed, our results show that mutation of ΔCjNC110 led to a significantly decreased although still present colonization ability in IA3902 compared to wild-type, indicating that this ncRNA has the potential to play a key role in modulating colonization by *C. jejuni*. The exact mechanism for the decreased colonization ability remains unknown, but could be due to any combination of factors identified in the phenotypic screening, including increased sensitivity to ROS, decreased autoagglutination ability, or altered quorum sensing ability; further work to elucidate how these factors may interact to affect *in vivo* colonization is thus warranted. The loss of the ability of ΔluxS to sustain colonization in chickens, as demonstrated in this study, has previously been described in IA3902 but the same phenotype was not observed in the closely related NCTC 11168 (8); the reason for this difference is currently unknown, but has been speculated to be related to strain-specific differences in the metabolism of methionine or *S-*adenosylmethionine (SAM) recycling which are also affected by the *luxS* mutation (28). Additional work in our lab has shown that the *luxS* mutation in IA3902 does disrupt the AMC, which functions to recycle SAM and produce methionine (64), however, DNA methylation in IA3902 was not affected by the *luxS* mutation (65). Comparative genomics reveals that NCTC 11168 does harbor an additional genomic system for methionine biosynthesis in the form of *metAB* which is not present in IA3902 and may represent an alternative pathway for synthesis of this required amino acid and an explanation for the observed difference in colonization ability. Further work is ongoing in our lab to determine if differing methionine metabolism may explain the difference in colonization ability of *luxS* mutants between strains, as well as to assess if mutation of CjNC110 in NCTC 11168 also leads to a defect in colonization, or, if similar to *luxS,* the phenotype differs from IA3902.

After attempting to compare various strains and methods of mutation of the *luxS* gene and finding wide variation in phenotypical and transcriptional changes, Adler *et al.* (29) suggested that some of the differences reported between *luxS* mutants of *C. jejuni* in various studies may be due to unknown polar affects caused by alteration of expression of CjNC110 based on the mutation strategy used for the *luxS* mutation. Our northern blot analysis confirmed that CjNC110 was still expressed at the same level as the wild-type strain in our *luxS* mutant construct. In addition, our phenotypic data clearly demonstrates very different phenotypes between ΔCjNC110 and ΔluxS in IA3902 for all phenotypes tested, with variation in the phenotype of ΔCjNC110ΔluxS based on the dominant phenotype of the two mutants. Thus, based on the work presented here, for our *luxS* mutant constructs, we can definitively state that polar effects on CjNC110 are not present.

Initial analysis of transcriptome changes in the ΔluxS mutant in our study revealed very few genes identified as differentially expressed, and the list of genes identified did not overlap with the list of differentially expressed genes generated primarily via microarray in various other strains (66–68). One explanation for this discrepancy may be differences between strains of *C. jejuni* or culture conditions when compared to these previous studies. However, in distinct contrast to the lack of similar findings to previous gene expression studies exhibited by the ΔluxS mutant, when the ΔCjNC110 and ΔluxS mutations were combined into ΔCjNC110ΔluxS, many of the previously noted transcriptional changes attributed to mutation of the *luxS* gene alone in a different strain of *C. jejuni*, 81-176, which also encodes and expresses CjNC110, became apparent (67). Of the 57 genes listed as differentially expressed by the *luxS* mutant under various conditions in He *et al.*(67), 23 were also found to be differentially expressed in our RNAseq data (any mutant construct). Of the 23 overlapping genes, no genes were found only in the ΔluxS mutant, and only the flagellar genes *flgE, flgG2,* and *flgG* and a putative helix-turn-helix protein CJSA_1449 were differentially expressed in both the ΔluxS and ΔCjNC110ΔluxS mutants. In contrast, three genes (*tpx, ptmB, ptmA*) were differentially expressed only in the ΔCjNC110 mutant. The He *et al.* study (67) also reported increased sensitivity to H_2_O_2_ in their *luxS* mutant construct, which more closely aligns with our ΔCjNC110 and ΔCjNC110ΔluxS mutant data and not ΔluxS. These findings strongly suggest that as hypothesized by Adler *et al.* (29), in some previous studies of ΔluxS, the main driver for the transcriptional and phenotypic changes seen was likely an inadvertent inactivation of both CjNC110 and *luxS,* and not *luxS* alone. Thus, further work is needed to re-evaluate the expression of CjNC110 in various *luxS* mutant constructs to determine if alteration of expression of CjNC110 is present and may account for some of the differences seen in phenotypes associated with *luxS* mutation in *C. jejuni*.

Of the genes identified in both our study and He *et al.* (67), a large number of the flagellar assembly hook-basal body associated proteins (FlgD, FlgE, FlgG, FlgG2, FlgH, FlgI, and FlgK) that are under the control of σ^54^ (RpoN) promotors were identified as differentially expressed (69, 70). Expression from σ^54^ promotors has been shown to require activation of the FlgRS two-component regulatory system (69, 71). Many of these genes showed opposite expression based on growth phase (decreased exponential, increased stationary), and it has been previously demonstrated that regulation by these sigma-factors is growth cycle dependent, with σ^54^-regulated genes typically expressed between σ^70^- and σ^28^-associated genes (69). It has been suggested that an additional unknown factor may control the temporal regulation of σ^54^- dependent flagellar genes (72), and the intergenic region between *luxS* and CJSA_1137 where CjNC110 is located has previously been identified to demonstrate differential expression in an *rpoN* (σ^54^) mutant in NCTC 11168 (19). Therefore, it is reasonable to consider based on previous studies and our results that the genomic region encompassing *luxS* and CjNC110 may play a role in the growth-phase dependent regulation of flagellar assembly.

While not explored further in our phenotypic studies, Rockhopper did identify two additional ncRNAs, CjNC140 and CjNC130/6S, as being differential expressed in our mutants; these ncRNAs have been previously identified and confirmed to be transcribed in multiple other *C. jejuni* strains (21). Predicted to be transcribed in the intergenic region upstream of *porA*, the CjNC140 ncRNA was upregulated during exponential growth in all three of the mutant strains when compared to wild-type (**Fig. S7**). As increased expression was not observed during stationary growth, this suggests that CjNC140 may serve as a regulator involved in mediating changes during different stages of bacterial growth. The fact that its expression is similarly altered under all 3 mutant conditions also suggests that the three genes, *luxS*, CjNC110 and CjNC140, may normally interact within the cell in some way to facilitate these changes. Point mutations in the nearby *porA* have also been recently determined to be sufficient to cause the abortion phenotype (10), making this ncRNA particularly interesting for further study.

Finally, while the specific mechanisms by which ncRNA CjNC110 interacts with each individual target were not elucidated in this study, the research presented herein clearly identified several specific roles of CjNC110 in the pathophysiology of *C. jejuni* IA3902. Indeed, even for well-studied ncRNAs such as MicA in *Enterobacteriacae,* new publications are still being produced which describe new roles and mechanisms of action even after a decade of research [recently reviewed in (73)]. Future work is necessary to investigate the hypotheses generated herein for genes and proteins that may be regulated by CjNC110 and to determine the mechanism by which CjNC110 interacts with each of the potential mRNA targets identified in this study.

In summary, the results of our research clearly demonstrate a significant role for CjNC110 in the pathobiology of *C. jejuni* IA3902. The collective results of the phenotypic and transcriptomic changes observed in our data complement each other and suggest that CjNC110 may be involved in regulation of energy taxis, flagellar glycosylation, cellular communication via quorum sensing, oxidative stress and chicken colonization in *C. jejuni* IA3902. This work provides for the first time valuable insights into the potential regulatory targets of the CjNC110 small ncRNA in the zoonotic pathogen *C. jejuni* and suggests a wide range of research avenues into the role of ncRNAs in the pathobiology of this important zoonotic pathogen.

## MATERIALS AND METHODS

### Bacterial strains, plasmids, primers and culture conditions

*C. jejuni* IA3902 was initially isolated from an outbreak of sheep abortion in Iowa during 2006 and has been utilized by our laboratory as the prototypical isolate of clone SA (3). W7 is a highly motile variant of the commonly utilized laboratory strain *C. jejuni* NCTC 11168 (8). *C. jejuni* strains and their isogenic mutants were routinely grown in Mueller-Hinton (MH) broth or agar plates (Becton-Dickinson, Franklin Lakes, NJ) at 42°C under microaerophilic conditions (5% O_2_, 10% CO_2_, 85% N_2_). For the mutant strains containing a chloramphenicol resistance cassette, 5 μg/mL chloramphenicol was added to either the broth or agar plates when appropriate. For strains containing a kanamycin resistance cassette, 30 μg/mL kanamycin was added to either the broth or agar plates when appropriate.

For genetic manipulations, *Escherichia coli* competent cells were grown at 37°C on Luria-Bertani (LB) agar plates or broth (Becton-Dickinson, Franklin Lakes, NJ) with shaking at 125 rpm. When appropriate, 50 μg/mL kanamycin, 20 μg/mL chloramphenicol or 100 μg/mL ampicillin was added to the broth or agar plates for selection of colonies. *Vibrio harveyi* strains were grown in autoinducer broth (AB) at 30°C with shaking at 175 rpm as described previously (74).

All strains used in this study are described in **Table 1**, with all relevant plasmids listed in **Table S8** and primer sequences and LNA DIG-labeled probe sequence listed in **Table S9**. **Fig. S8** illustrates the genetic modification strategy described below to create the mutants utilized in this study. All strains were maintained in 20% glycerol stocks at −80°C and passaged from those stocks as needed for experimental procedures.

### Creation of *C. jejuni* ΔCjNC110 and ΔCjNC110ΔluxS mutants in IA3902

An isogenic CjNC110 mutant of *C. jejuni* IA3902 was constructed via deletional mutagenesis utilizing a combination of synthetic double-stranded DNA (dsDNA) fragments and traditional cloning methods. Based on previously published data depicting the proposed transcriptional start site for CjNC110 (21), the coding region of CjNC110 in IA3902 and the prototypical *C. jejuni* strain NCTC 11168 were first confirmed to be identical. Then, a 200 bp section of the IA3902 genome starting 20 bp upstream and including the entire 137 nt transcript of the CjNC110 sequence as predicted in Dugar *et al.* (21) was replaced with 820 bp of the promoter and coding sequence of the chloramphenicol acetyltransferase (*cat*) gene of *Campylobacter coli* plasmid C-589 (75). Synthetic dsDNA including approximately 500 bp upstream and 500 bp downstream of the region replaced with the *cat* cassette was then synthesized in 4 fragments of 500 bp each with overlapping homologous ends (Integrated DNA Technologies, Coralville, IA). The Gibson Assembly method was then utilized to assemble the synthetic dsDNA fragments using the Gibson Assembly Master Mix (New England Biolabs, Ipswich, MA) (76). Following assembly, primers (CjNC110F2 and CjNC110R2) were designed to amplify a 1785 bp product of the assembled dsDNA; PCR amplification was achieved using TaKaRa Ex Taq DNA Polymerase (ClonTech, Mountain View, CA). This amplified PCR product was then cloned into the pGEM-T Easy Vector using T4 ligase (Promega, Madison, WI), resulting in the construction of pCjNC110::cat which was then transformed into chemically competent *E. coli* DH5α (New England Biolabs, Ipswich, MA). Transformants were then selected on LB agar plates containing chloramphenicol (20 μg/ml), ampicillin (100 μg/ml), and ChromoMax IPTG/X-Gal Solution (Fisher Scientific, Pittsburgh, PA). pCjNC110::cat was purified from the transformed *E. coli* using the QIAprep Miniprep kit (QIAGEN, Germantown, MD) and confirmed by PCR to contain the construct again using the CjNC110F2 and CjNC110R2 primers.

The pCjNC110::cat plasmid DNA was then introduced to *C. jejuni* W7, a highly motile variant of NCTC 11168, as a suicide vector and the deletion transferred into the genome of *C. jejuni* W7 via homologous recombination. Transformants were selected on MH agar plates containing chloramphenicol (5 μg/ml) and deletional mutagenesis was again confirmed via PCR analysis and Sanger sequencing to create *C. jejuni* W7ΔCjNC110. Following confirmation, natural transformation was used to move the gene deletion into *C. jejuni* IA3902 as previously described to create *C. jejuni* IA3902ΔCjNC110 (77). Natural transformation was again used to move the CjNC110 gene deletion into the previously created *luxS* insertional mutant *C. jejuni* IA3902ΔluxS (8) to create the double knockout mutant *C. jejuni* IA3902ΔCjNC110ΔluxS. Transformants were selected on MH agar plates containing chloramphenicol (5 μg/mL) and kanamycin (30 μg/mL) and confirmed via PCR analysis and Sanger sequencing of the CjNC110 region along with the entire upstream (*luxS*) and downstream (CJSA_1137) genes using primers Cj1198F1 and Cj1199R3. All colonies were screened for presence/absence of motility as described below, and only colonies with verified motility were used for future studies. Expression of CjNC110 in wild-type and elimination of expression in the ΔCjNC110 mutant was confirmed via northern blot analysis using an LNA custom designed probe and 15 μg of total RNA as described below.

### Creation of *C. jejuni* ΔCjNC110 complement in IA3902

Complementation of ΔCjNC110 was achieved via insertion of the coding region into the intergenic region of the 16S and 23S rRNA operon (*rrs-rrl*) of IA3902ΔCjNC110 via homologous recombination using plasmid pRRK. A copy of the CjNC110 gene including 190bp upstream of the transcription initiation location (total of 570bp) was cloned into an *XbaI* site of pRRK located upstream of the kanamycin resistance determinant to create pRRK-CjNC110 as previously described in our lab (78) with some modifications of the original protocol (79). Briefly, the CjNC110 gene and flanking regions of IA3902 were amplified using forward and reverse primers designed with an *XBaI* restriction enzyme site (CjNC110cF1 and CjNC110cR1). Following PCR amplification with TaKaRa Ex Taq DNA Polymerase (ClonTech, Mountain View, CA), the PCR product and pRRK plasmid were digested with the *XBaI* restriction endonuclease (New England Biolabs, Ipswich, MA). The digested plasmid was then incubated twice with Antarctic Phosphatase (New England Biolabs, Ipswich, MA) to prevent self-ligation of cut plasmid. Following ligation and transformation, transformants were selected on LB agar plates containing chloramphenicol (20 μg/ml) and kanamycin (50 μg/ml kanamycin) yielding a plasmid containing the construct pRRK::CjNC110 in *E. coli* DH5α. Following PCR confirmation and Sanger sequencing to confirm preservation of correct gene sequence, the plasmid was then utilized as a suicide vector and the gene was inserted into the intergenic region of the 23S rRNA operon (*rrs-rrl*) of IA3902ΔCjNC110 via homologous recombination to generate IA3902ΔCjNC110c transformants. The transformants were selected on MH agar plates containing chloramphenicol (5 μg/ml) and kanamycin (30 μg/ml) and confirmed via PCR analysis and Sanger sequencing to contain both the expected ΔCjNC110 deletional mutation and ΔCjNC110c insertion, validating successful genetic complementation. All positive colonies were screened for presence/absence of motility as described below and a motile complement was selected to create IA3902ΔCjNC110c. Expression of CjNC110 in ΔCjNC110c was confirmed via northern blot analysis using an LNA custom designed probe and 15 μg of total RNA as described below.

### Growth curves

Two separate growth curves were completed in triplicate utilizing the same method as described below with the only variation being the addition of the complement strain ΔCjNC110c to the second growth curve. To begin the growth curve, the A_600_ of overnight cultures were adjusted to 0.5 using sterile MH broth on a Genesys 10S VIS spectrophotometer (Thermo Scientific™, Waltham, MA). Cultures were then diluted 1:10 for a final targeted starting A_600_ of 0.05 in 90 mL MH broth and placed in a sterile 250 mL Erlenmeyer glass flask. Cultures were incubated at 42°C under microaerophilic conditions with shaking at 125 rpm for 30 hours with removal of samples from the flasks at designated time points [3, 6, 9, 12, 24, 27 (2^nd^ growth curve only) and 30 hours]. For the first growth curve (which included the following strains: IA3902 wild-type, ΔCjNC110, ΔluxS and ΔCjNC110ΔluxS), samples were processed as described below for RNA isolation as well as assessed for A_600_ and actual colony counts using the drop-plate method as previously described (80). For the second growth curve (which included ΔCjNC110c in addition to the four previous strains), samples were collected and processed for A_600_ and as described below for qRT-PCR and assessment of AI-2 levels via the bioluminescence assay. The A_600_ over time were statistically analyzed for the two independent growth curves, using a two-way ANOVA with repeated measures and Dunnett’s multiple comparison test (GraphPad Prism).

### Total RNA extraction

Broth culture samples processed for total RNA extraction were either centrifuged at 8000 x *g* for 2 minutes (for sequencing and qRT-PCR) or 10,000 x *g* for 4 minutes (northern blot) at 4°C immediately following collection. Following pelleting of the cells, the supernatant was decanted and 1 mL QIAzol Lysis Reagent (QIAGEN) was added to resuspend the cell pellet and protect the RNA. QIAzol-protected cultures were then stored at −80°C for up to two months prior to proceeding with total RNA isolation. Total RNA isolation was performed using the miRNeasy Mini Kit (QIAGEN) followed by purification using the RNeasy MinElute Cleanup kit (QIAGEN) as previously described (24). RNA quality was measured using the Agilent 2100 Bioanalyzer RNA 6000 Nano kit (Agilent Technologies, Santa Clara, CA) and all RNA samples utilized for downstream RNAseq library preparation had an RNA integrity number (RIN) of >9.0, indicating high quality RNA. Verification of complete removal of any contaminating DNA was performed via PCR amplification of a portion of the CJSA_1356 gene, which is unique to *C. jejuni* IA3902, using primers SA1356F and SA1356R (81).

### RNAseq library preparation and sequencing

The 3 hour (exponential phase) and 12 hour (stationary phase) timepoints were selected for RNAseq analysis based on assessment of log_10_ CFU/mL (**Fig. S2A**). 2.5 µg of confirmed DNA-free total RNA was treated with Ribo-Zero rRNA Removal Kit for Bacteria according to the manufacturer’s instructions (Illumina, San Diego, CA). Following rRNA removal, the ribosomal depleted total RNA was purified using the RNeasy MinElute Cleanup kit (QIAGEN) using the same modifications as described previously (24). Following clean-up, the RNA quality, quantity, and rRNA removal efficiency was then analyzed via the Agilent 2100 Bioanalyzer RNA 6000 Pico kit (Agilent Technologies). Library preparation for sequencing on the Illumina HiSeq platform was completed using the TruSeq stranded mRNA HT library preparation kit (Illumina) with some modifications as described previously (24). The pooled library was then submitted to the Iowa State University DNA Facility and sequenced on an Illumina HiSeq 2500 machine in high-output single read mode with 100 cycles.

### Differential gene expression analysis of RNAseq data

Rockhopper (http://cs.wellesley.edu/~btjaden/Rockhopper/) was used to analyze the differences in gene expression between strains and timepoints; the standard settings of the program were utilized for analysis (30). Following computational analysis via Rockhopper, a change in gene expression was deemed significant when the *Q*-value (false discovery rate) was <0.05 and a >1.5 fold change in expression was observed. Visual assessment of read count data was performed using the Integrated Genome Viewer (IGV) (https://www.broadinstitute.org/igv/) (82, 83). Assessment of function of the differentially expressed genes was performed using the Clusters of Orthologous Groups (COG) (84) as previously described in IA3902 (5). Venn diagrams were used to depict overlap of genes differentially regulated in multiple mutant strains and were generated using BioVenn (http://www.cmbi.ru.nl/cdd/biovenn/index.php) (85). KEGG Pathways was used to perform metabolic pathway analysis (http://www.genome.jp/kegg/pathway.html) (31).

In addition to determining differential gene expression between the mutant and wild-type strains of IA3902, Rockhopper has the capability to predict non-coding RNAs present within the data. Prior to further manual analysis, Rockhopper predicted a total of 59 ncRNAs present in IA3902 (57 chromosomal, 2 pVir). Manual curation was performed to remove candidate non-coding RNAs if they were related to the 16S or 23S genes as these were thought to be spuriously identified due to Ribo-Zero depletion differences between replicates. In addition, a single antisense RNA was discarded due to incorrect annotation of the *rnpB* gene to the wrong strand in IA3902.

### Northern blot detection of CjNC110

For northern blot detection of CjNC110, two separate experiments utilizing identical conditions were performed as previously described with some modifications (86). Experiment 1 included wild-type IA3902, ΔCjNC110, and ΔCjNC110c, while the second included wild-type and ΔluxS. For both, bacterial cells were grown to early stationary phase of growth in MH broth and total RNA isolation performed as described above. The total RNA input was normalized to 15 μg using Qubit™ BR RNA Assay Kit (Invitrogen™, USA) for each respective strain. For northern blot detection, total RNA from bacterial cells (15 μg) and a pre-stained DynaMarker® RNA High Ladder (Diagnocine LLC, Hackensack, NJ) were loaded onto 1.3% denaturing agarose gel (Bio-rad, Hercules, CA), consisting of 1x MOPS buffer (Fisher, USA) and 5% solution of 37% formaldehyde (Fisher, USA), electrophoresed at 100V for 1h and transferred to positively-charged nylon membrane (Roche, Indianapolis, IN) using a Vacuum Blot System (Bio-Rad) transferring at 60mbar for 2 hours. To make RNA cross-linking solution, 245 μl of 12.5 M 1-methylimidazole (Sigma, St. Louis, MO) was suspended in 9 mL RNase free water, maintaining a pH of 8.0. Then, a total of 0.753 g 1-ethyl-3-(3-dimethylaminopropyl) carbodiimide (EDC) was added and the total volume adjusted to 24 mL with RNase-free water. RNA was then cross-linked to the membrane by placing the nylon membrane, side devoid of RNA, on saturated 3mm Whatman chromatography paper and incubating it in the freshly prepared EDC solution at 60°C for 2 hours. After cross-linking the membrane was washed with RNase-free water. Next, CjNC110-LNA DIG-labeled probe (/5’DigN/ GCACATCAGTTTCAT/3’Dig_N/) (QIAGEN) was added to 15 mL DIG EasyHyb™ Buffer (Roche) to reach a final concentration of 25 ng/mL. Hybridization was performed using a hybridization oven and bottle, rotating at 54°C for 12 hours. After overnight incubation, subsequent washes and incubations were performed again using the hybridization chamber. The blot was first washed twice with low stringency buffer (2x SSC, 0.1% SDS) for 15 minutes at 60°C, and three times with high stringency buffer (0.1% SSC and 0.1% SDS) for 10 minutes at 60°C. DIG Washing and DIG Blocking Buffers were prepared using the DIG Wash and Block Set (Roche, USA). The blot was washed twice with Washing Buffer for 10 minutes at 37°C. Next, the blot was incubated in 1x Blocking Buffer for 3 hours at room temperature. DIG antibody solution was added by mixing Blocking Buffer with DIG antibody at a ratio of 1:5,000. The blot was then incubated for 30 minutes at room temperature. Next, excess DIG antibody was removed by rinsing with the DIG Washing Buffer at 25°C a total of four times for 15 minutes each. For development and detection, the membrane was placed in Development/Detection Buffer for 10 minutes, and 2 mL ready-to-use CDP-*Star*® (Roche, USA) was added to cover the blot completely. To image the blot, Chemidoc Imager (Bio-Rad) was used, exposing the blot for 5 minutes on the chemiluminescence setting. Image layout and annotations were performed using GraphPad Prism. ImageLab software (Bio-Rad) was used to generate a standard curve using RF and nt size of the ladder to predict the average band size of the target RNA.

### Enumeration of *luxS* transcriptional levels utilizing qRT-PCR

To validate the RNAseq results and compare *luxS* expression between mutants, quantitative reverse transcriptase PCR (qRT-PCR) was performed using total RNA isolated from IA3902 wild-type, ΔCjNC110, and ΔCjNC110c purified in three separate biological replicates of broth cultures grown to 12 hours from the second growth curve described above. Total RNA was extracted and qRT-PCR was performed using the iScript cDNA Synthesis Kit (Bio-Rad, Hercules, CA) and 1000 ng of total RNA according to the manufacturer’s instructions as previously described (64). Each sample was normalized to the same starting amount of cDNA using Qubit™ BR DNA Assay Kit (Invitrogen). cDNA purity was measured using the NanoDrop ND-1000 spectrophotometer. qPCR assays were run using the SsoAdvanced™ Universal SYBR® Green Supermix (Bio-Rad, Hercules, CA) and the CFX Maestro™ Real-Time PCR detection system (Bio-Rad). Dilutions of cDNA template for both standards and all unknowns were run in triplicate with reaction volumes of 10 μl. Amplification of converted cDNA occurred with 35 cycles of denaturation at 95°C for 10 seconds and then annealing for each primer pair (**Table S9**) at 58°C for 30 seconds. Prior to analysis, both standards curves (16S and *luxS*) were experimentally validated to have high efficiency 90%> of amplification and precision R^2^=0.98 or greater. Relative fold change in *luxS* mRNA expression between IA3902 wild-type, ΔCjNC110, and ΔCjNC110c was calculated using the ISU Gallup Method Equation (87). Statistical analyses was performed using one-way ANOVA (GraphPad Prism) to determine significance in gene expression levels. A *P*-value of <0.05 was considered significant.

### Liquid chromatography with tandem mass spectrometry-based proteomic analysis

For extraction of whole cell protein, bacterial cells from IA3902 wild-type, ΔCjNC110 and ΔCjNC110c were grown to stationary phase of growth at 16 hours on MH agar. Lawn cultures were then harvested from the plates using 1 mL of MH broth. Collected lawns were normalized to an A_600_ of 0.2 and aliquoted into 1 mL technical replicates. This was repeated for three biological replicates. To wash the cells, each aliquot was re-suspended in 1 mL of PBS, cells were spun down at 8000 x *g* for 3 minutes, and the supernatant was decanted. Next, 50 µl of lysis buffer (filtered sterilized water + 1% Triton X-100) was added to the protein pellets for resuspension [adapted from (88)]. The cells were then mechanically sheared by boiling at 96°C for 5 minutes followed by rapid pipetting. After 3 cycles of boiling-shearing, total cell extracts were centrifuged at 10,000 × *g* for 5 min to remove cell debris. The supernatants were collected, and protein concentration was determined using the Qubit™ Protein Assay and Fluorometer (Invitrogen™). Crude protein was then submitted to the Iowa State Protein Facility. Crude protein extracts were digested overnight in solution with trypsin/Lys-C (25-50 µg). After digestion, Pierce™ Peptide Retention Time Calibration Mixture (PRTC) (ThermoFisher Scientific) was spiked into the samples to serve as an internal control (25 fmol/µL). The PRTC areas were used to normalize collision energies, retention times, and peak intensities to allow for quantitative analysis between samples.

Proteomic analysis was performed via liquid chromatography with tandem mass spectrometry (LC-MS/MS) using a Q Exactive™ Hybrid Quadrupole-Orbitrap Mass Spectrometer (Thermo Scientific™) system. The system was coupled with EASY-nLC 1200 nanopump with integrated auto-sampler (Thermo Scientific™). Liquid chromatography was used to separate the peptides followed by fragmentation and MS/MS analysis. The resulting intact fragmentation pattern was compared to a theoretical fragmentation pattern (MASCOT) to find peptides that can be used to identify high-confidence proteins. Minora Feature Detector was used for label-free quantification to detect and quantify discovered isotopic clusters.

Abundances of proteins were compared by grouping by strain and averaging the abundances of each protein identified. For data filtering and data imputation, PANDA-view, a freely available proteomic software (https://sourceforge.net/projects/panda-view/), was used (89). Briefly, only proteins identified in two out of three biological replicates were included in analysis. Nearest neighbor (kNN) imputation was used to fill any missing protein values (90). All data was log transformed using log_2_. For statistical analysis, two-way ANOVA with false discovery rate (*Q-*value) correction set at 0.05 was performed (Graphpad Prism). Graphical hierarchical clustering heat maps were generated using freely available MetaboAnalyst software (91). The top 20 proteins were group and analyzed using the following heat map settings: distance measurement of Euclidean and the average clustering algorithm (91, 92).

### *Vibrio harveyi* bioluminescence assay

Culture samples collected from time points 3, 6, 9 and 12 hours (2 mL each timepoint) from the second growth curve described above were centrifuged at 20,000 x *g* for 5 minutes at 4°C. The supernatant was then removed from the cell pellet and filter-sterilized using a 0.2 μm syringe filter to create cell-free supernatant representative of the extracellular environment (E-CFS). The cell pellet was then washed twice with PBS; following each wash, the cells were spun again at 8000 x *g* and the resulting supernatant was completely removed from the cell pellet. Both the E-CFS and the cell pellet were then frozen at −80°C. For processing of the cell pellet to create intracellular cell-free supernatant (I-CFS), samples were thawed on ice and then resuspended in 600 μl of ice-cold MH broth. The resuspended cell pellets were then lysed using a Bullet Blender® (NextAdvance, Troy, NY). To lyse the cells, 1.5 mL microcentrifuge tubes (Corning, Corning, NY) were loaded with sterilized 0.9-2.0 mm diameter stainless steel beads (NextAdvance). The Bullet Blender was used for 5 minutes at max speed and stored at 4°C. Lysed cellular debris were spun at 8000 x *g* for 2 minutes using a microcentrifuge, and the resulting supernatant filtered using a 0.20 μm sterile syringe filter, resulting in intracellular cell-free supernatant (I-CFS). Both E-CFS and I-CFS were frozen at −80 °C until proceeding with the bioluminescence assay.

Autoinducer-2 levels within the collected supernatant for both intracellular (I-CFS) and extracellular (E-CFS) samples were then enumerated using the *Vibrio harveyi* bioluminescence assay as previously described (32). Briefly, 10 µL of each CFS sample was added to 90 µL AB media containing a 1:5000 dilution of the reporter strain, *V. harveyi* strain BB170 (93), in triplicate. Relative light units (RLU) were measured every 15 minutes over 8 hours using the FLUOstar Omega (BMG Labtech, Ortenburg, Germany). MH broth and AB media were used as negative controls, while E-CFS collected from *V. harveyi* strain BB152 (74) was used as a positive control. The timepoints utilized for analysis were those occurring during the nadir of values for the negative control wells and at a standardized duration of time (approximately 3 hours) following initiation of increasing values for the positive wells. Results reported are the average of three independent growth curves with three technical replicates used at each timepoint. Statistical analysis was conducted using a two-way ANOVA with repeated measures and Sidak’s multiple comparisons test (GraphPad Prism). A *P*-value of <0.05 was considered significant.

### Motility assay

Motility was determined via inoculation of plates consisting of MH broth with 0.4% agar as previously described in our laboratory (8). Briefly, the A_600_ of overnight cultures was adjusted to 0.3 using sterile MH broth on a Genesys 10S VIS spectrophotometer (Thermo Scientific™). A 1 µL volume inoculation stick was then dipped into a set volume of the standardized culture contained in the bottom of a 15 mL conical tube which was then used to make a stab inoculation into the center of the freshly made motility agar (MH broth with 0.4% Bacto agar) with a new inoculation stick for each plate. Plates were incubated at 42°C under microaerophilic conditions as described above with the exception that the plates were incubated right-side up and in a single layer. Measurement of the outermost reach of the halo was performed at 30 hours following inoculation for the first set of experiments and at 24 hours for the second set of experiments. All strains were assessed for average motility using six biological replicates consisting of six technical replicates during each independent study. The six experiments were statistically analyzed using a one-way ANOVA and differences between each strain assessed via Tukey’s multiple comparisons test (GraphPad Prism). A *P*-value of <0.05 was considered significant.

### Autoagglutination assay

Autoagglutination was assessed according to the method described previously (40) with some modifications. Briefly, the A_600_ of overnight cultures was adjusted to 1.0 in sterile Dulbecco’s phosphate buffered saline (PBS) (Corning cellgro, Manassas, VA) using a Genesys 10S VIS spectrophotometer (Thermo Scientific™). The suspension was then aliquoted (2 mL each) into standard glass culture tubes. One subset of cultures were kept at controlled room temperature (25°C) under microaerophilic conditions; the others were incubated at 37°C microaerophilic. At 24 hours, 1 mL of the upper aqueous phase was carefully removed and A_600_ measured to determine autoagglutination activity. All strains were assessed in quadruplicate at each temperature in three independent experiments. The three experiments were statistically analyzed using one-way ANOVA and differences between each strain assessed via Tukey’s multiple comparisons test (GraphPad Prism). A *P*-value of <0.05 was considered significant.

### Hydrogen peroxide disk inhibition assay

The hydrogen peroxide (H_2_0_2_) disk inhibition assay was performed as previously described with some modifications (63). Briefly, *C. jejuni* IA3902 wild-type, and mutant strains were grown to log phase in MH broth. The bacteria were harvested by centrifugation at 12,000 x *g* for 2 minutes and subsequently normalized by resuspension into MH broth to A_600_ of 1.0. Next, 24 mL of melted MH agar was combined with 1 mL of the normalized bacterial suspension on petri plates, mixed well, and allowed to solidify. 20 μL of aqueous 3% H_2_O_2_ solution (Sigma-Aldrich, St. Louis, MO) was then pipetted onto 6-mm diameter disk, which was placed at the center of the plate containing the bacterial agar suspension. Plates were incubated for 24 hours at 42°C under microaerophilic conditions. The zones of H_2_O_2_ sensitivity were measured in millimeters (mm), by measuring the diameter of the clear zones. The H_2_0_2_ disk inhibition assay was repeated three times using freshly prepared 3% H_2_0_2_ with 4 technical replicates per strain. The data was statistically evaluated using one-way ANOVA and Dunnett’s multiple comparison test. A *P-*value of <0.05 was deemed significant.

### Chicken colonization

All studies involving animals were approved by the Iowa State University Institutional Animal Care and Use Committee (IACUC) prior to initiation (1-18-8675-G) and followed all appropriate animal care guidelines. Chicken colonization studies were performed as previously described in our laboratory utilizing 1 day-old broiler chicks obtained from a commercial hatchery (8). A subset of chicks from each group were screened via cloacal swabs plated on MH agar containing both selective supplement and growth supplement (MH+sel+sup) (Oxoid, ThermoFisher Scientific) and found to be negative for *Campylobacter* carriage prior to inoculation. At 3 days of age, chicks were inoculated by oral gavage with 200 µl of bacterial suspension containing approximately 1 x 10^7^ of one of the tested strains. Groups of chicks inoculated with specific strains were housed in separate brooders with no contact between the groups. At pre-determined timepoints post-inoculation (5 days, 12 days, and 19 days), 6 chicks from each group were randomly selected for humane euthanasia which was immediately followed by necropsy. Cecal contents were harvested aseptically and stored on ice until further processing could be completed. Cecal samples were then weighed and subjected to a 10-fold serial dilution series, plated on MH+sel+sup agar, and grown under routine culture conditions as described above. To confirm that the isolates recovered were in fact the mutant strain and no cross contamination occurred, representative colonies were selected from each strain and timepoint, grown in MH+sel+sup broth, and plated on the respective antimicrobial selective agar of the mutant strain as described above. In addition, a minimum of two CFU from each group was collected for PCR analysis and confirmation of no cross-contamination between biological groups. The detection limit of the assay was determined to be 100 CFU/g; for samples that failed to reach the optimal target range of 30-300 CFU on the initial dilution plate, a count of 3 x 10^3^ was used to enable statistical analysis. For statistical analysis, one-way ANOVA and Tukey’s multiple comparison test was used to determine significant differences in colonization between biological groups (GraphPad Prism), using the null hypothesis that colonization rates between groups are the same. A *P*-value of <0.05 was considered significant.

### Data availability

The RNAseq dataset generated in this publication has been deposited in the NCBI Database under BioProject ID Number PRJNA590513 (http://www.ncbi.nlm.nih.gov/bioproject/590513).

## ACKNOWLEDGEMENTS

The authors would like to thank Dr. Lei Dai and Dr. Zuowei Wu for technical assistance in mutant construction and northern blot technique.

## Notes

### Competing Interest Statement

The authors have declared no competing interest.

