## Supplemental figures and tables for "Small non-coding RNA CjNC110 influences motility, autoagglutination, AI-2 localization, hydrogen peroxide sensitivity and chicken colonization in *Campylobacter jejuni*"

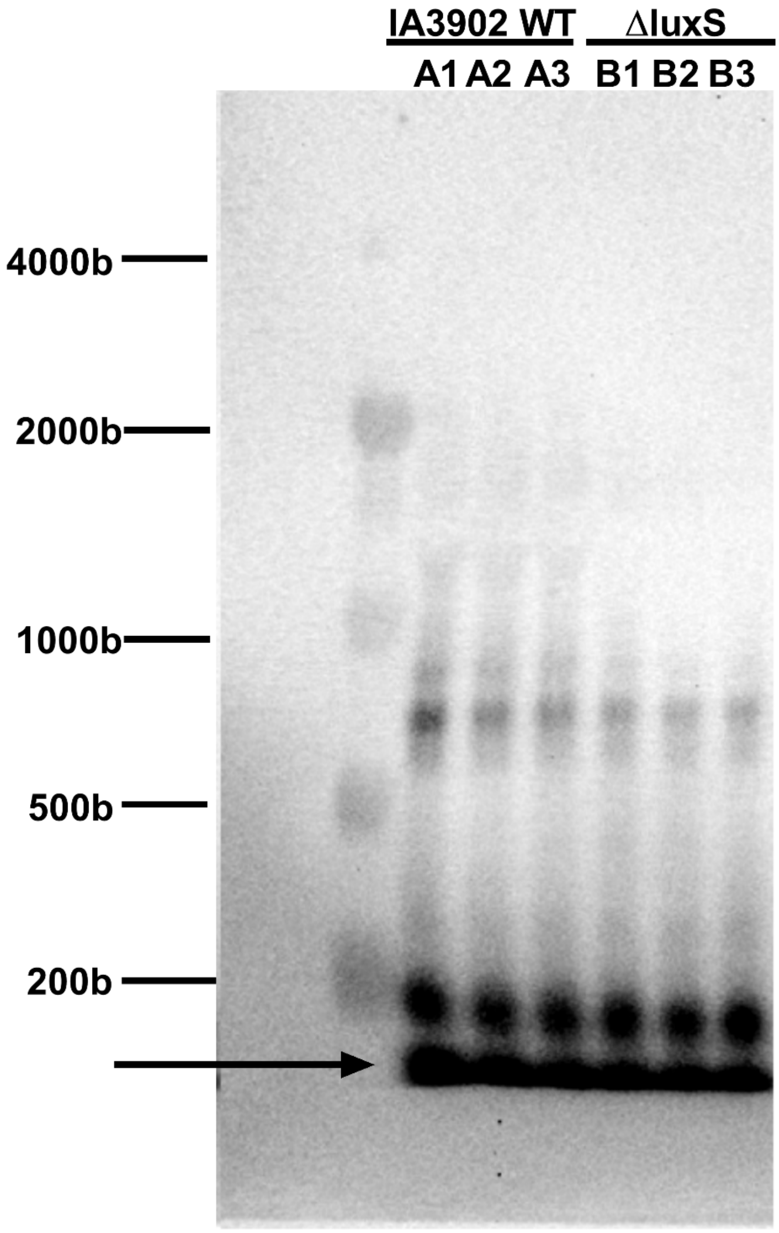


**FIG S1**. Northern blotting demonstrates similar expression of CjNC110 in wild-type IA3902 and ΔluxS. Cultures for RNA extraction were collected at early stationary phase of growth from three independent growth curves and northern blot analysis was conducted using 15 μg of total RNA in each of three separate lanes per strain tested. The arrow indicates the most prominent band which corresponds to the previously predicted size of CjNC110 in *C. jejuni*.

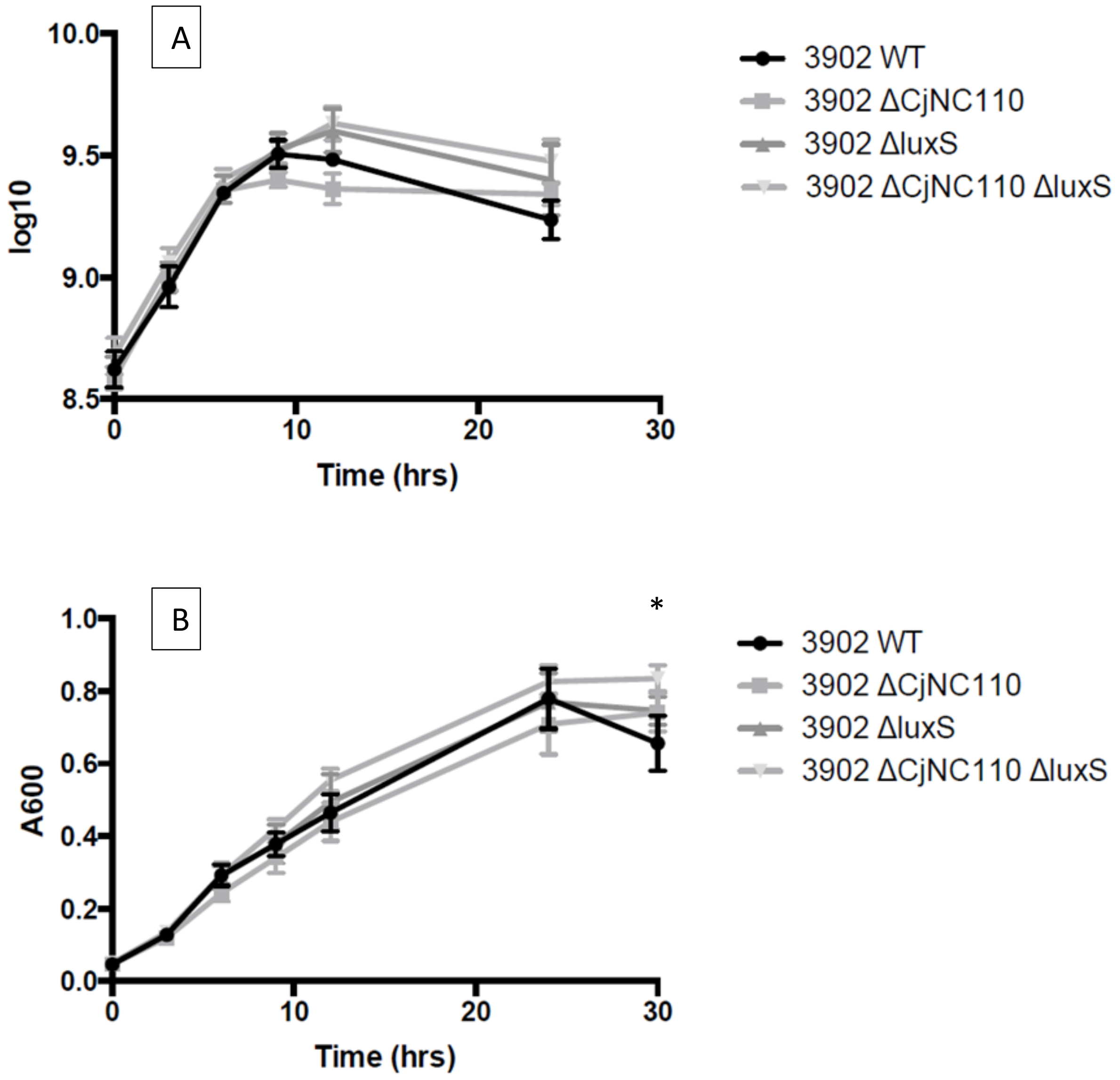


**FIG S2. Shaking growth curve utilized for RNA extraction for RNAseq of wild-type IA3902 and isogenic mutants (mean ± SEM).** $A_{600}$ (A) and log10 CFU/mL (B) results of four replicates of a shaking growth curve performed in 250 mL Erlenmeyer flasks under microaerophilic conditions in MH broth. Analysis via two-way ANOVA revealed a statistically significant difference between strains ($p < 0.05$), however, multiple comparison analysis of individual time points and strains when compared to wild type growth revealed that the only time point where the differences was considered to be statistically significant was at 30 hours for all strains (denoted by *).

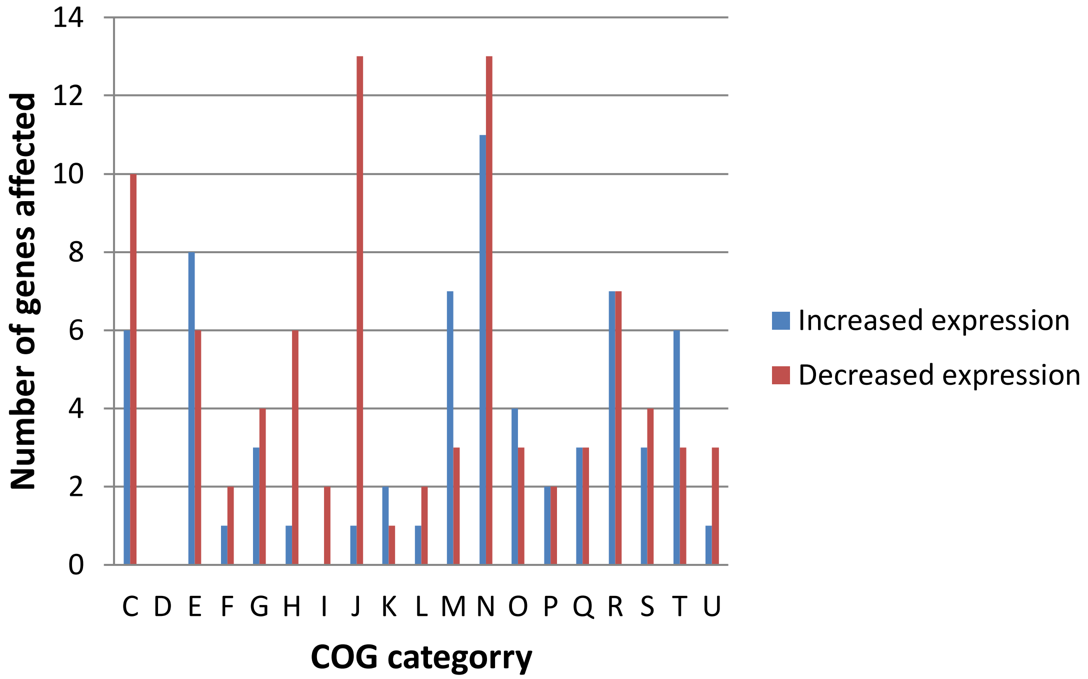


**FIG S3**. **COG functional categories of differentially expressed genes in the ΔCjNC110ΔluxS mutant during exponential growth.** Clusters of Orthologous Groups (COG) categories are indicated on the x-axis, with the number of genes enriched shown on the y-axis; blue bars indicate increased expression, red bars indicate decreased expression. C - Energy production and conversion; D - Cell cycle control, mitosis and meiosis; E - Amino acid transport and metabolism; F - Nucleotide transport and metabolism; G - Carbohydrate transport and metabolism; H - Coenzyme transport and metabolism; I - Lipid transport and metabolism; J – Translation; K – Transcription; L - Replication, recombination and repair; M - Cell wall/membrane biogenesis; N - Cell motility; O - Posttranslational modification, protein turnover, chaperones; P - Inorganic ion transport and metabolism; Q - Secondary metabolites biosynthesis, transport and catabolism; R - General function prediction only; S - Function unknown; T - Signal transduction mechanisms; U - Intracellular trafficking and secretion; V - Defense mechanisms; W - Extracellular structures

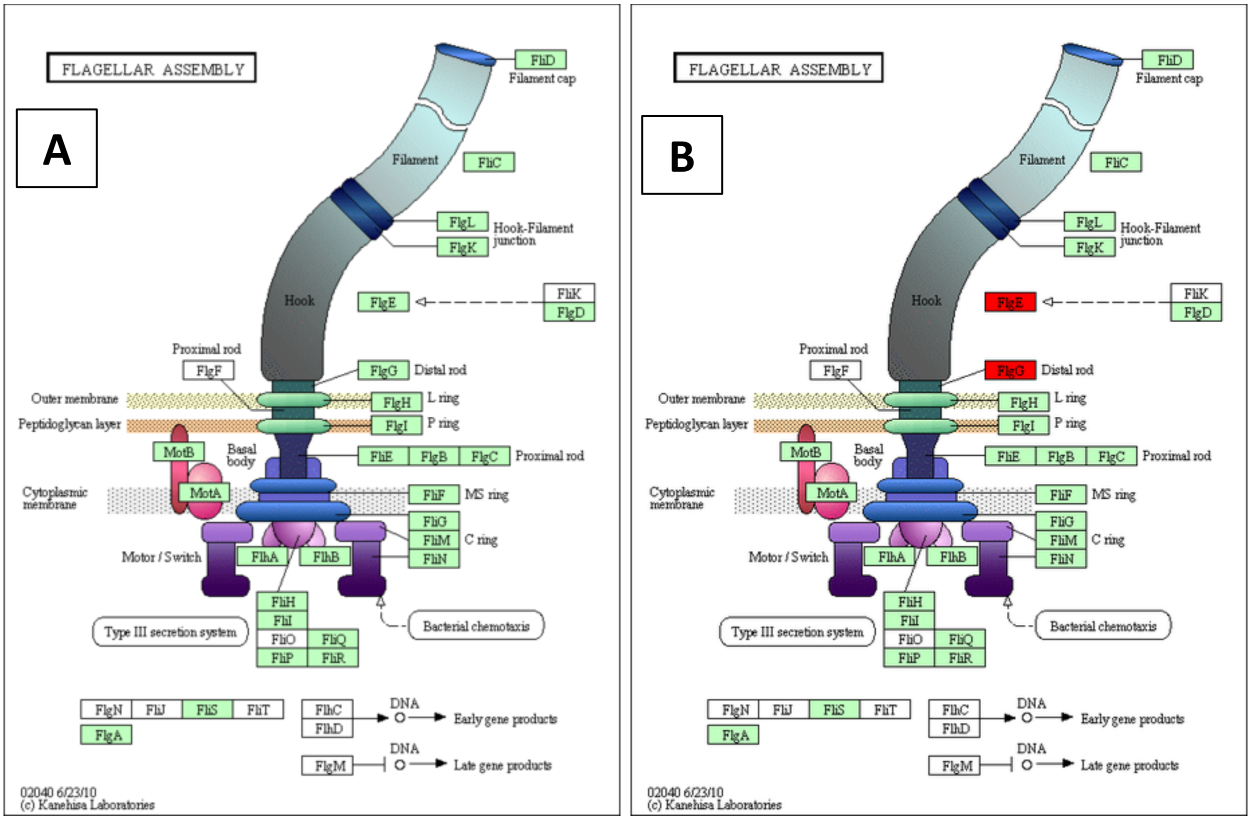


**Fig S4. KEGG Pathway for flagellar assembly in *C. jejuni:* genes affected by the single mutation of either ΔCjNC110 (A) or ΔluxS (B) mutation during stationary growth phase.** Blue color indicates downregulation of gene expression, red color indicates up regulation of gene expression, green indicates that the gene is present in *C. jejuni* IA3902, and white indicates that the gene is not present in IA3902.

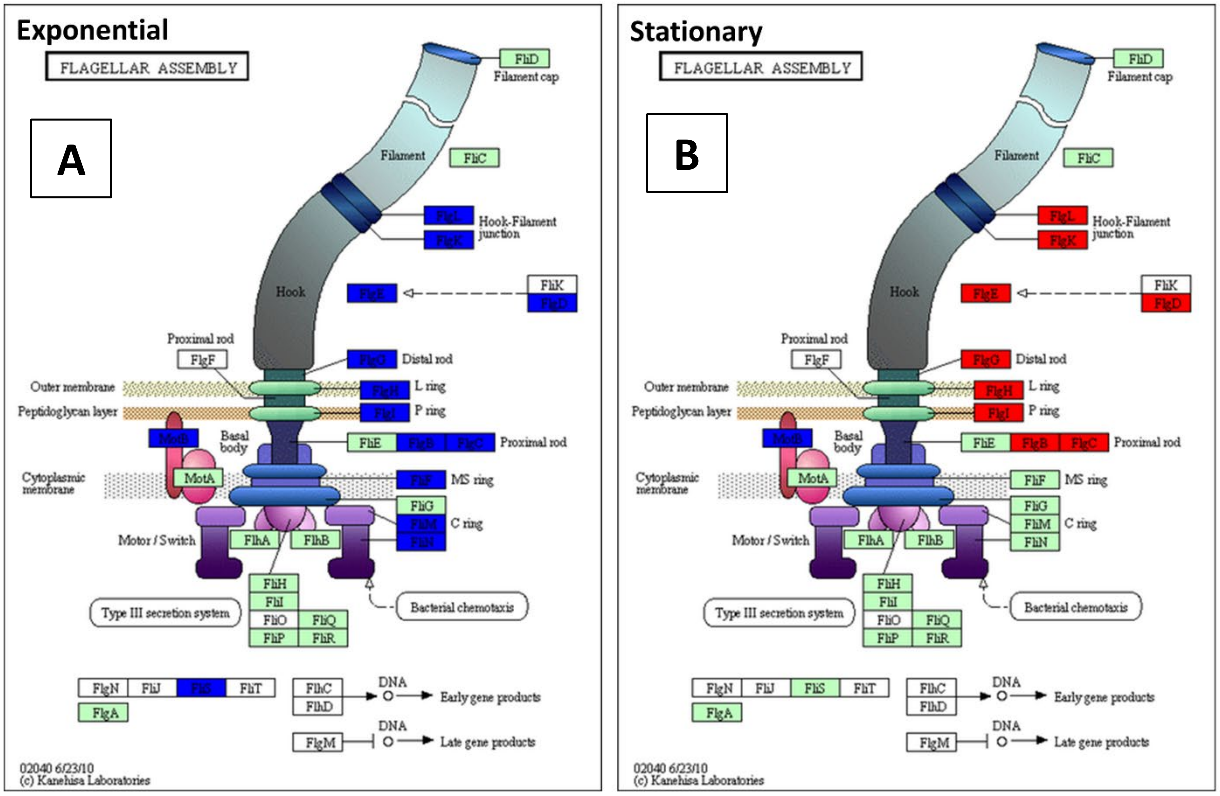


**FIG S5. KEGG Pathway analysis demonstrates alteration of expression of multiple genes involved in flagellar assembly in *C. jejuni* in the ΔCjNC110ΔluxS mutant during both exponential (A) and stationary (B) growth phases.** Blue color indicates downregulation of gene expression, red color indicates up regulation of gene expression, green indicates that the gene is present in C. jejuni IA3902, and white indicates that the gene is not present in IA3902.

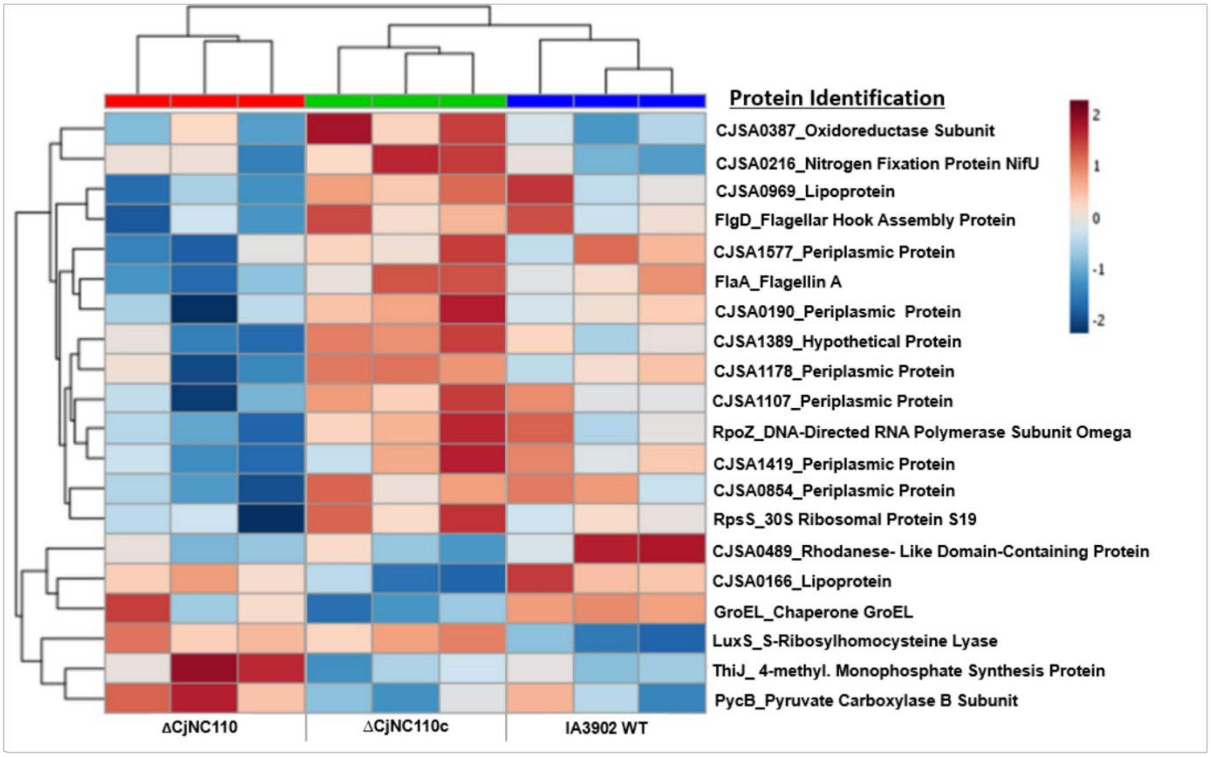


**FIG S6. Heatmaps reveal differential protein clustering patterns induced in both ΔCjNC110 and ΔCjNC110c when compared to IA3902 wild-type (WT).** All raw abundances were normalized, imputated, transformed to log2 scale values, and clustered based on resulting hierarchical patterns. The top 20 proteins were grouped and analyzed using heat map settings: distance measurement of Elucidean and average clustering algorithm.

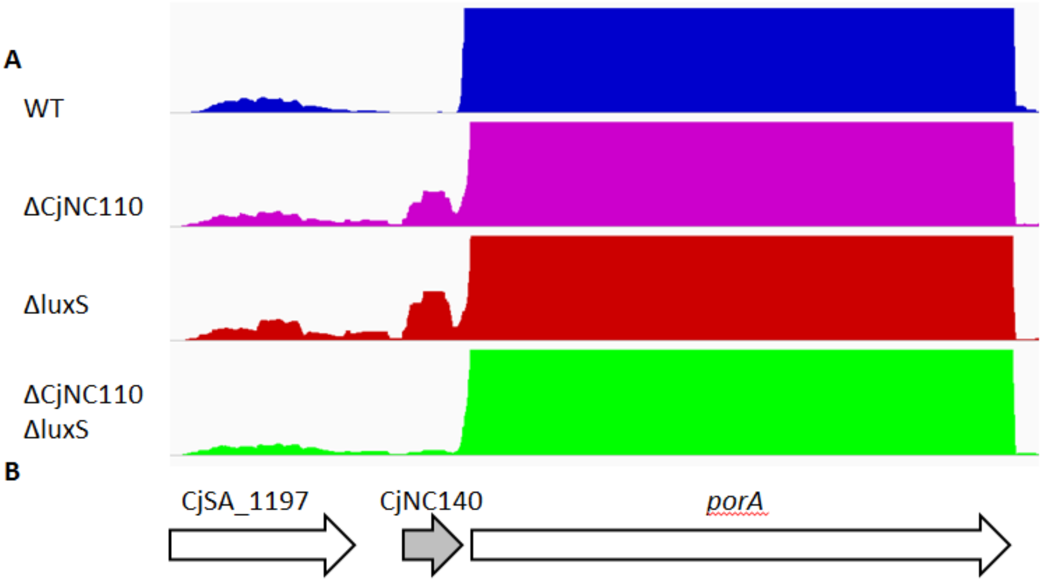


**FIG S7. Graphic view of CjNC140 expression from RNAseq**. (A) A screen capture from IGV of the *porA* and CjNC140 regions of the genome, corresponding to the genome structure as depicted in (B), of all of the strains sequenced using RNAseq.

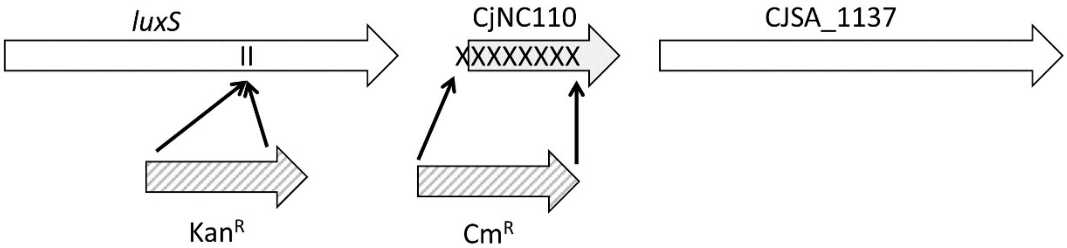


**FIG S8. Graphic view of mutation strategy for ΔluxS and ΔCjNC110 in IA3902**. Insertional mutagenesis was utilized to create ΔluxS, while insertional deletion was utilized to create ΔCjNC110.

### Supplemental tables

**Table S1.** Summary of RNAseq results following rRNA depletion and strand specific library preparation – successfully aligned reads.

| Library | Total reads | Number of successfully aligned reads | | | Percent mapped reads |
|---|---|---|---|---|---|
| | | Chromosome | pVir | Total | |
| IA3902 WT-3hr-1 | 4167800 | 4060985 | 27341 | 4088326 | 97.4% |
| IA3902 WT-3hr-2 | 3785630 | 3686851 | 39278 | 3726129 | 98.4% |
| IA3902 WT-3hr-3 | 3574775 | 3460965 | 43366 | 3504331 | 98.0% |
| IA3902 WT-12hr-1 | 4179614 | 4042874 | 43963 | 4086837 | 97.8% |
| IA3902 WT-12hr-2 | 4286716 | 4159744 | 45225 | 4204969 | 98.1% |
| IA3902 WT-12hr-3 | 2968022 | 2103578 | 21429 | 2125007 | 71.6% |
| IA3902 ΔCjNC110-3hr-1 | 3995995 | 3853431 | 61455 | 3914886 | 98.0% |
| IA3902 ΔCjNC110-3hr-2 | 3640043 | 3512435 | 36437 | 3548872 | 97.5% |
| IA3902 ΔCjNC110-3hr-3 | 4114232 | 3943639 | 63649 | 4007288 | 97.4% |
| IA3902 ΔCjNC110-12hr-1 | 4646416 | 4488809 | 57391 | 4546200 | 97.8% |
| IA3902 ΔCjNC110-12hr-2 | 4158490 | 4028072 | 60311 | 4088383 | 98.3% |
| IA3902 ΔCjNC110-12hr-3 | 5109264 | 3721853 | 27654 | 3749507 | 73.4% |
| IA3902 ΔluxS-3hr-1 | 3251135 | 3143754 | 51047 | 3194801 | 98.3% |
| IA3902 ΔluxS-3hr-2 | 3026876 | 2939706 | 26144 | 2965850 | 98.0% |
| IA3902 ΔluxS-3hr-3 | 2315385 | 1202081 | 17248 | 1219329 | 52.7% |
| IA3902 ΔluxS-12hr-1 | 4497066 | 4378061 | 42525 | 4420586 | 98.3% |
| IA3902 ΔluxS-12hr-2 | 3294082 | 3190105 | 36272 | 3226377 | 97.9% |
| IA3902 ΔluxS-12hr-3 | 19473343 | 15405040 | 174495 | 15579535 | 80.0% |
| IA3902 ΔCjNC110ΔluxS-3hr-1 | 4480648 | 4339847 | 59374 | 4399221 | 98.2% |
| IA3902 ΔCjNC110ΔluxS-3hr-2 | 3117870 | 3013513 | 41098 | 3054611 | 98.0% |
| IA3902 ΔCjNC110ΔluxS-3hr-3 | 3680686 | 2866303 | 43328 | 2909631 | 79.1% |
| IA3902 ΔCjNC110ΔluxS-12hr-1 | 5329709 | 5181117 | 58908 | 5240025 | 98.3% |
| IA3902 ΔCjNC110ΔluxS-12hr-2 | 4179965 | 3853179 | 42962 | 3896141 | 93.2% |
| IA3902 ΔCjNC110ΔluxS-12hr-3 | 4018571 | 3875936 | 42462 | 3918398 | 97.5% |
| **AVERAGE** | 4553847 | 4102162 | 48473 | 4150635 | 92.2% |
| **MINIMUM** | 2315385 | 1202081 | 17248 | 1219329 | 52.7% |
| **MAXIMUM** | 19473343 | 15405040 | 174495 | 15579535 | 98.4% |
| **MEDIAN** | 4066402 | 3853305 | 43145 | 3905514 | 97.9% |
| **TOTAL** | 109292333 | | | 99615240 | |

**Table S2.** Summary of RNAseq results following rRNA depletion and strand specific library preparation – read mapping results.

| | Percent mapped reads | | | | | | | | |
|---|---|---|---|---|---|---|---|---|---|
| | Chromosome | | | | | | | pVir | |
| | Protein coding | | Ribosomal RNA | | Other known RNA | | Unannotated | Protein coding | Unannotated |
| Library | Sense | Antisense | Sense | Antisense | Sense | Antisense | regions | Sense | regions |
| IA3902 WT-3hr-1 | 72 | 1 | 15 | 0 | 5 | 3 | 3 | 93 | 6 |
| IA3902 WT-3hr-2 | 85 | 1 | 4 | 0 | 5 | 3 | 2 | 92 | 8 |
| IA3902 WT-3hr-3 | 87 | 1 | 3 | 0 | 4 | 2 | 3 | 91 | 9 |
| IA3902 WT-12hr-1 | 88 | 1 | 2 | 0 | 5 | 2 | 3 | 91 | 8 |
| IA3902 WT-12hr-2 | 86 | 1 | 3 | 0 | 5 | 1 | 3 | 91 | 8 |
| IA3902 WT-12hr-3 | 74 | 0 | 17 | 1 | 4 | 1 | 2 | 91 | 9 |
| IA3902 ΔCjNC110-3hr-1 | 88 | 1 | 1 | 0 | 4 | 2 | 3 | 90 | 9 |
| IA3902 ΔCjNC110-3hr-2 | 74 | 0 | 16 | 0 | 4 | 2 | 3 | 94 | 6 |
| IA3902 ΔCjNC110-3hr-3 | 90 | 1 | 2 | 0 | 3 | 2 | 3 | 90 | 9 |
| IA3902 ΔCjNC110-12hr-1 | 87 | 0 | 2 | 0 | 6 | 2 | 3 | 91 | 8 |
| IA3902 ΔCjNC110-12hr-2 | 89 | 1 | 1 | 0 | 6 | 1 | 2 | 91 | 9 |
| IA3902 ΔCjNC110-12hr-3 | 69 | 0 | 21 | 1 | 4 | 1 | 3 | 92 | 7 |
| IA3902 ΔluxS-3hr-1 | 89 | 1 | 1 | 0 | 4 | 2 | 3 | 90 | 10 |
| IA3902 ΔluxS-3hr-2 | 78 | 0 | 11 | 0 | 4 | 3 | 3 | 93 | 7 |
| IA3902 ΔluxS-3hr-3 | 65 | 1 | 27 | 2 | 3 | 1 | 2 | 89 | 11 |
| IA3902 ΔluxS-12hr-1 | 86 | 1 | 3 | 0 | 6 | 2 | 3 | 91 | 8 |
| IA3902 ΔluxS-12hr-2 | 88 | 1 | 2 | 0 | 5 | 2 | 3 | 91 | 8 |
| IA3902 ΔluxS-12hr-3 | 79 | 0 | 11 | 0 | 5 | 1 | 2 | 91 | 9 |
| IA3902 ΔCjNC110ΔluxS-3hr-1 | 88 | 1 | 3 | 0 | 4 | 2 | 3 | 91 | 8 |
| IA3902 ΔCjNC110ΔluxS-3hr-2 | 88 | 1 | 3 | 0 | 3 | 2 | 3 | 91 | 9 |
| IA3902 ΔCjNC110ΔluxS-3hr-3 | 80 | 1 | 11 | 0 | 3 | 1 | 3 | 90 | 10 |
| IA3902 ΔCjNC110ΔluxS-12hr-1 | 88 | 1 | 2 | 0 | 5 | 2 | 3 | 92 | 8 |
| IA3902 ΔCjNC110ΔluxS-12hr-2 | 84 | 1 | 5 | 0 | 5 | 2 | 3 | 90 | 9 |
| IA3902 ΔCjNC110ΔluxS-12hr-3 | 82 | 0 | 5 | 0 | 6 | 2 | 3 | 92 | 8 |
| **AVERAGE** | 83 | 1 | 7 | 0 | 5 | 2 | 3 | 91 | 8 |

**Table S3.** List of non-coding RNAs predicted by Rockhopper to exist using RNAseq transcriptomics.

| Transcription | | | | | | | |
|---|---|---|---|---|---|---|---|
| **Start** | **Stop** | **Strand** | **Length** | **LFG** | **RFG** | **Comments** | **Previously identified?** |
| **Pseudogenes with transcription** | | | | | | | |
| 1047422 | 1046663 | - | 759 | | CjSA_1052 | CjSA_1052 - annotated as a pseudogene | no |
| 1331028 | 1331442 | + | 414 | | CjSA_1323 | CjSA_1323 - annotated as a degenerate pseudogene | no |
| 1630614 | 1630418 | - | 196 | | CjSA_1630 | 3' region of CjSA_1630 - annotated as a pseudogene | no |
| 1630874 | 1630848 | - | 26 | | CjSA_1630 | 5' region of CjSA_1630 - annotated as a pseudogene | no |
| 1457497 | 1457695 | + | 198 | | CjSA_1444 | CjSA_1444 - annotated as a pseudogene | no |
| 1555611 | 1555599 | - | 12 | | CjSA_1543 | CjSA_1543 - annotated as a pseudogene | no |
| **Predicted *cis* RNA** | | | | | | | |
| 67249 | 67227 | - | 22 | SRP | CjSA_0046 | Overlaps 5' end of CjSA_0046 (pseudogene) | no |
| 1183929 | 1183946 | + | 17 | CjSA_1188 | CjSA_1189 | antisense to 5' UTR *purD* | Yes - CjNC130/6S |
| **Predicted *trans* RNA** | | | | | | | |
| 71394 | 71379 | - | 15 | CjSA_0049 | CjSA_0050 | intergenic CjSA_0049 and CjSA_0050 | no |
| 197215 | 197309 | + | 94 | CjSA_0191 | CjSA_0192 | intergenic CjSA_0191 and CjSA_0192 | no |
| 199473 | 199366 | - | 107 | CjSA_0192 | CjSA_1093 | intergenic *tetO* and CjSA_0192 | no |
| 250049 | 249967 | - | 82 | CjSA_0242 | CjSA_0243 | intergenic CjSA_0242 and CjSA_0243 | Yes - CjNC20 |
| 271319 | 271271 | - | 48 | CjSA_0265 | CjSA_0265 | intergenic *peb3* and *lpxB* | no |
| 441750 | 441796 | + | 46 | CjSA_0444 | CjSA_0445 | intergenic *rplK* and *rplA*, opposite strand | no |
| 462280 | 462334 | + | 54 | CjSA_0462 | CjSA_0463 | intergenic *rpsL* and *rpsG* | no |
| 601205 | 601168 | - | 37 | CjSA_0604 | CjSA_0605 | intergenic *ppA and msrA* | no |
| 675392 | 675240 | - | 152 | CjSA_0682 | CjSA_0682 | intergenic *dnaE* and CjSA_0681 | Yes - CjNC60 |
| 726401 | 726387 | - | 14 | CjSA_0728 | CjSA_0729 | Intergenic, possible 5' UTR of CjSA_0728 | no |
| 738637 | 738648 | + | 11 | CjSA_0737 | CjSA_0738 | intergenic *napA and napG* | no |
| 1112058 | 1112049 | - | 9 | CjSA_1127 | CjSA_1128 | very small intergenic | no |
| 1132258 | 1132313 | + | 55 | CjSA_1136 | CjSA_1137 | intergenic *luxS* and CjSA_1137, 3' end of luxS | no |
| 1153480 | 1153499 | + | 19 | CjSA_1157 | CjSA_1158 | intergenic CjSA_1157 and *groEL* | no |
| 1193307 | 1193399 | + | 92 | CjSA_1197 | CjSA_1198 | intergenic CjSA_1197 and *porA* | Yes - CjNC140 |

**Table S3 continued**

| | | | | | | | |
|---|---|---|---|---|---|---|---|
| 1243977 | 1243958 | - | 19 | CjSA_1247 | CjSA_1248 | intergenic CjSA_1247 and CjSA_1248 | no |
| 1358594 | 1358557 | - | 37 | CjSA_1350 | CjSA_1351 | intergenic CjSA_1350 and CjSA_1351 | no |
| 1397360 | 1397371 | + | 11 | CjSA_1387 | CjSA_1388 | intergenic CjSA_1387 and CjSA_1388 | no |
| 1397574 | 1397607 | + | 33 | CjSA_1388 | CjSA_1389 | intergenic CjSA_1388 and CjSA_1389 | no |
| 1577169 | 1577153 | - | 16 | CjSA_1567 | CjSA_1568 | Intergenic CjSA_1568 and *nhaA1* | no |
| 1589879 | 1589849 | - | 30 | CjSA_1582 | CjSA_1583 | intergenic CjSA_1582 and *eno* | no |
| 1602665 | 1602650 | - | 15 | CjSA_1592 | CjSA_1593 | intergenic CjSA_1593 and *gltA* | no |
| 1619089 | 1619055 | - | 34 | CjSA_1918 | CjSA_1619 | intergenic CjSA_1618 and CjSA_1619 | no |
| **Predicted antisense RNA** | | | | | | | |
| 180678 | 180760 | + | 82 | CjSA_0173 | CjSA_0174 | antisense: CJSA_0174 - junction of 2 genes | no |
| 198877 | 199050 | + | 173 | CjSA_0191 | CjSA_1092 | antisense: CJSA_0192 - antisense to 3'end | no |
| 907368 | 907271 | - | 97 | CjSA_0910 | CjSA_0911 | antisense: CJSA_0911 - antisense to 5'end | no |
| 1489555 | 1489543 | - | 12 | | CjSA_1476 | antisense: CJSA_1476 - antisense to 5'end | no |
| 1549038 | 1549028 | - | 10 | | CjSA_1535 | antisense: CJSA_1535 - antisense to 3'end | no |
| 1586230 | 1586134 | - | 96 | | CjSA_1576 | antisense: CJSA_1576 | no |
| 1198424 | 1198330 | - | 94 | | CjSA_1202 | antisense: *recR* - antisense to 3'end | no |
| **Predicted pVir** | | | | | | | |
| 7643 | 7658 | + | 15 | pVir0009 | pVir0010 | intergenic | no |
| 25261 | 25403 | + | 142 | pVir0033 | pVir0032 | intergenic | yes - CJpv2 |

**Table S4.** Differential gene expression in the IA3902 ΔCjNC110 mutant as determined by RNAseq.

| Name | Synonym | COG Code | Product | Expression (RPKM) WT | Expression (RPKM) ΔCjNC110 | Significance Q Value | Significance Fold change |
|---|---|---|---|---|---|---|---|
| | | | **Exponential phase** | | | | |
| **Genes downregulated** | | | | | | | |
| *hisF* | CJSA_1252[a] | E | imidazole glycerol phosphate synthase subunit HisF | 84 | 30 | 1.2E-04 | -2.8 |
| *pseA* | CJSA_1254[a] | D | pseudaminic acid biosynthesis PseA protein | 129 | 50 | 1.1E-05 | -2.6 |
| *neuB2* | CJSA_1263[b] | M | N-acetylneuraminate synthase | 171 | 57 | 7.0E-09 | -3.0 |
| *ptmA* | CJSA_1268[b] | QR | flagellin modification protein A | 107 | 41 | 4.8E-05 | -2.6 |
| - | CJSA_1352 | M | putative sugar transferase | 38 | 17 | 4.3E-03 | -2.2 |
| *rpsN* | CJSA_1603 | J | 30S ribosomal protein S14 | 541 | 289 | 3.7E-02 | -1.9 |
| **Genes upregulated** | | | | | | | |
| - | CJSA_0008 | S | hypothetical protein | 20 | 62 | 9.0E-11 | 3.1 |
| *cetB* | CJSA_1127[c] | T | bipartate energy taxis response protein cetB | 28 | 78 | 3.1E-03 | 2.8 |
| *cetA* | CJSA_1128[c] | NT | bipartate energy taxis response protein cetA | 45 | 100 | 2.8E-04 | 2.2 |
| - | CJSA_1261 | D | hypothetical protein | 34 | 66 | 1.0E-02 | 1.9 |
| CjNC140 | predicted RNA | - | 92nt length, positive strand from 1193307 to 1193399 | 10 | 59 | 9.0E-17 | 5.9 |
| - | predicted RNA | - | 55nt length, positive strand from 1132258 to 1132313 (3' end of *luxS*) | 36 | 157 | 1.9E-07 | 4.4 |
| | | | **Stationary phase** | | | | |
| **Genes downregulated** | | | | | | | |
| *panC* | CJSA_0271 | H | pantoate-beta-alanine ligase | 173 | 109 | 4.3E-03 | -1.6 |
| - | CJSA_0559 | - | putative lipoprotein | 135 | 91 | 3.0E-02 | -1.5 |
| - | CJSA_0687 | O | M48 family peptidase | 33 | 22 | 3.2E-04 | -1.5 |
| *tpx* | CJSA_0735 | O | thiol peroxidase | 1966 | 1326 | 5.4E-03 | -1.5 |
| - | CJSA_0785 | S | hypothetical protein | 77 | 51 | 7.8E-03 | -1.5 |
| *ciaB* | CJSA_0859 | - | invasion antigen B | 68 | 43 | 2.9E-02 | -1.6 |

**Table S4 continued**

| | | | | | | | |
|---|---|---|---|---|---|---|---|
| - | CJSA_1102 | S | hypothetical protein | 173 | 105 | 1.2E-06 | -1.6 |
| *petC* | CJSA_1122 | C | putative ubiquinol-cytochrome C reductase cytochrome C subunit | 474 | 310 | 1.3E-03 | -1.5 |
| - | CJSA_1137 | R | 2OG-Fe(II) oxygenase | 39 | 15 | 1.1E-05 | -2.6 |
| - | CJSA_1244 | - | hypothetical protein | 30 | 18 | 7.9E-04 | -1.7 |
| *hisF* | CJSA_1252 | E | imidazole glycerol phosphate synthase subunit HisF | 56 | 31 | 6.3E-06 | -1.8 |
| *neuB2* | CJSA_1263[b] | M | N-acetylneuraminate synthase | 58 | 31 | 5.4E-03 | -1.9 |
| - | CJSA_1266[b] | R | hypothetical protein | 43 | 16 | 5.4E-06 | -2.7 |
| *ptmB* | CJSA_1267[b] | M | cylneuraminate cytidylyltransferase (flagellin modification) | 158 | 42 | 2.8E-34 | -3.8 |
| *ptmA* | CJSA_1268[b] | QR | flagellin modification protein A | 102 | 24 | 2.5E-35 | -4.3 |
| - | CJSA_t0002 | - | Ile tRNA | 45 | 28 | 1.1E-03 | -1.6 |
| **Genes upregulated** | | | | | | | |
| *hcrA* | CJSA_0713 | K | heat-inducible transcription repressor | 306 | 659 | 3.5E-04 | 2.2 |
| - | CJSA_0716[d] | R | hypothetical protein | 25 | 68 | 1.8E-06 | 2.7 |
| - | CJSA_0717[d] | S | hypothetical protein | 20 | 45 | 1.5E-03 | 2.3 |
| - | CJSA_1107[e] | - | hypothetical protein | 50 | 100 | 2.9E-02 | 2.0 |
| *omp50* | CJSA_1108[e] | - | 50 kda outer membrane protein precursor | 57 | 118 | 2.9E-03 | 2.1 |
| *luxS* | CJSA_1136 | T | S-ribosylhomocysteinase | 505 | 1250 | 5.6E-07 | 2.5 |
| - | CJSA_1261 | D | hypothetical protein | 10 | 29 | 7.8E-11 | 2.9 |
| - | predicted RNA | - | 55nt length, positive strand from 1132258 to 1132313 (3' end of *luxS*) | 27 | 699 | 0.0E+00 | 25.9 |

[a, b, c, d, e] = denotes genes within the same operon as predicted by Rockhopper

underlined = signifcantly different expression at both timepoints

**Table S5.** Differential gene expression in the IA3902 ΔluxS mutant as determined by RNAseq.

| Name | Synonym | COG Code | Product | Expression (RPKM) | | Significance | |
|---|---|---|---|---|---|---|---|
| | | | | **WT** | **ΔluxS** | **Q Value** | **Fold change** |
| | | | **Exponential phase** | | | | |
| **Genes downregulated** | | | | | | | |
| - | CJSA _1350 | H | putative methyltransferase | 2047 | 543 | 2.2E-03 | -3.8 |
| - | predicted RNA | - | 55nt length, positive strand from 1132258 to 1132313 (3' end of *luxS*) | 36 | 1 | 7.2E-271 | -36.0 |
| **Genes upregulated** | | | | | | | |
| *dnaN* | CJSA_0002 | L | DNA polymerase III subunit beta | 55 | 98 | 4.0E-03 | 1.8 |
| - | CJSA_0008 | S | hypothetical protein | 20 | 53 | 1.0E-08 | 2.7 |
| *trpF* | CJSA_0321[a] | E | N-(5phosphoribosyl)anthranilate isomerase | 21 | 59 | 1.2E-03 | 2.8 |
| *trpB* | CJSA_0322[a] | E | tryptophan synthase subunit beta | 19 | 37 | 5.3E-02 | 1.9 |
| - | CJSA_0337 | - | hypothetical protein | 9 | 24 | 3.1E-02 | 2.7 |
| - | CJSA_0370 | - | hypothetical protein | 22 | 53 | 2.0E-02 | 2.4 |
| - | CJSA_0732 | - | hypothetical protein | 88 | 149 | 3.5E-02 | 1.7 |
| - | CJSA_1017 | S | flagellar assembly factor FliW | 163 | 247 | 4.2E-02 | 1.5 |
| - | CJSA_1131 | - | hypothetical protein | 10 | 27 | 5.3E-02 | 2.7 |
| - | CJSA_1301 | O | putative nucleotidyltransferase | 19 | 33 | 2.7E-02 | 1.7 |
| - | CJSA_1352 | M | putative sugar transferase | 38 | 61 | 2.6E-02 | 1.6 |
| - | CJSA_1449 | R | putative helix-turn-helix containing protein | 43 | 105 | 1.1E-06 | 2.4 |
| - | CJSA_1549 | G | hypothetical protein | 31 | 79 | 4.8E-07 | 2.5 |
| - | CJSA_pVir0042 | - | hypothetical protein | 10 | 29 | 2.2E-02 | 2.9 |
| CjNC140 | predicted RNA | - | 92nt length, positive strand from 1193307 to 1193399 | 10 | 72 | 4.2E-28 | 7.2 |

**Table S5 continued**

| | | | | | | | |
|---|---|---|---|---|---|---|---|
| | | | **Stationary phase** | | | | |
| **Genes downregulated** | | | | | | | |
| - | CJSA_0560 | - | hypothetical protein | 193 | 121 | 2.5E-02 | -1.6 |
| - | CJSA_0620 | E | M24 family peptidase | 262 | 147 | 7.4E-05 | -1.8 |
| - | CJSA_1349[b] | G | hypothetical protein | 60 | 24 | 1.1E-03 | -2.5 |
| - | CJSA_1350[b] | H | putative methyltransferase | 387 | 208 | 3.5E-09 | -1.9 |
| *acs* | CJSA_1453 | I | acetyl-coenzyme A synthetase | 110 | 65 | 1.7E-03 | -1.7 |
| *leuC* | CJSA_1626 | E | 3-isopropylmalate dehydratase large subunit | 58 | 36 | 1.0E-02 | -1.6 |
| - | predicted RNA | - | 55nt length, positive strand from 1132258 to 1132313 (3' end of *luxS*) | 27 | 1 | 0.0E+00 | -27.0 |
| **Genes upregulated** | | | | | | | |
| *flgE* | CJSA_0043 | N | flagellar hook protein | 124 | 244 | 1.7E-03 | 2.0 |
| *flgG2* | CJSA_0661[c] | N | flagellar basal-body rod protein | 308 | 502 | 5.0E-03 | 1.6 |
| *flgG* | CJSA_0662[c] | N | flagellar basal-body rod protein FlgG | 292 | 538 | 1.3E-03 | 1.8 |
| *hcrA* | CJSA_0713 | K | heat-inducible transcription repressor | 306 | 530 | 2.2E-03 | 1.7 |
| - | CJSA_1107 | - | hypothetical protein | 50 | 102 | 3.9E-03 | 2.0 |
| *nrfA* | CJSA_1292 | P | putative periplasmic cytochrome C | 189 | 324 | 4.1E-02 | 1.7 |

[a, b, c] = denotes genes within the same operon as predicted by Rockhopper

underlined = signifcantly different expression at both timepoints

**Table S6.** Differential gene expression in the IA3902 ΔCjNC110ΔluxS mutant as determined by RNAseq.

| Name | Synonym | COG Code | Product | Expression (RPKM) WT | Expression (RPKM) ΔCjNC110 ΔluxS | Significance Q Value | Significance Fold change |
|---|---|---|---|---|---|---|---|
| | | | **Exponential phase** | | | | |
| **Genes downregulated** | | | | | | | |
| - | CJSA_0014 | S | hypothetical protein | 274 | 166 | 6.8E-04 | -1.7 |
| - | CJSA_0041[a] | - | hypothetical protein | 83 | 51 | 4.8E-04 | -1.6 |
| *flgD* | CJSA_0042[a] | N | flagellar basal body rod modification protein | 159 | 97 | 1.0E-03 | -1.6 |
| *flgE* | CJSA_0043[a] | N | flagellar hook protein | 140 | 85 | 5.8E-07 | -1.6 |
| - | CJSA_0067 | C | iron-sulfur cluster binding protein | 737 | 419 | 1.2E-02 | -1.8 |
| *accD* | CJSA_0118 | I | acetyl-CoA carboxylase subunit beta | 86 | 48 | 3.7E-05 | -1.8 |
| *trxA* | CJSA_0138 | O | Thioredoxin | 897 | 548 | 1.2E-03 | -1.6 |
| *panB* | CJSA_0272 | H | 3-methyl-2-oxobutanoate | 64 | 44 | 4.9E-03 | -1.5 |
| *motB* | CJSA_0310 | N | flagellar motor protein MotB | 178 | 94 | 1.7E-03 | -1.9 |
| *fliN* | CJSA_0325 | NU | flagellar motor switch protein | 175 | 119 | 3.4E-03 | -1.5 |
| *rpsU* | CJSA_0343 | J | 30S ribosomal protein S21 | 4581 | 2456 | 9.1E-03 | -1.9 |
| *frdB* | CJSA_0383 | C | fumarate reductase iron-sulfur subunit | 1352 | 842 | 3.7E-02 | -1.6 |
| *flaG* | CJSA_0514[b] | N | flagellar protein FlaG | 559 | 379 | 3.0E-05 | -1.5 |
| *fliS* | CJSA_0516[b] | NUO | flagellar protein FliS | 399 | 266 | 8.6E-04 | -1.5 |
| - | CJSA_0521 | - | hypothetical protein | 57 | 32 | 8.1E-04 | -1.8 |
| - | CJSA_0569 | R | sodium-dependent transporter | 38 | 24 | 8.9E-03 | -1.6 |
| *pstS* | CJSA_0581 | P | phosphate transport system substrate-binding protein | 47 | 25 | 2.0E-02 | -1.9 |
| *hslV* | CJSA_0628 | O | ATP-dependent protease peptidase subunit | 237 | 162 | 5.7E-03 | -1.5 |
| *flgH* | CJSA_0651 | N | flagellar basal body L-ring protein | 144 | 75 | 3.6E-02 | -1.9 |
| *flgG2* | CJSA_0661[c] | N | flagellar basal-body rod protein | 220 | 138 | 1.3E-05 | -1.6 |
| *flgG* | CJSA_0662[c] | N | flagellar basal-body rod protein FlgG | 359 | 213 | 1.3E-05 | -1.7 |
| *mogA* | CJSA_0689 | H | molybdenum cofactor biosynthesis protein | 198 | 132 | 3.9E-02 | -1.5 |

**Table S6 continued**

| | | | | | | | |
|---|---|---|---|---|---|---|---|
| *aspB* | CJSA_0718 | E | aspartate transaminase | 122 | 79 | 1.5E-03 | -1.5 |
| - | CJSA_0788 | F | putative oxidoreductase | 143 | 80 | 2.3E-02 | -1.8 |
| *flgL* | CJSA_0833 | N | flagellar hook-associated protein FlgL | 142 | 79 | 1.3E-02 | -1.8 |
| *rpmH* | CJSA_0906 | - | 50S ribosomal protein L34 | 471 | 288 | 1.2E-04 | -1.6 |
| - | CJSA_0920 | Q | hypothetical protein | 392 | 223 | 1.7E-08 | -1.8 |
| - | CJSA_1093 | C | cytochrome c553 | 3846 | 1960 | 1.9E-03 | -2.0 |
| - | CJSA_1102 | S | hypothetical protein | 190 | 120 | 1.2E-02 | -1.6 |
| *dctA* | CJSA_1130 | C | C4-dicarboxylate transport protein | 147 | 85 | 9.3E-08 | -1.7 |
| *luxS* | CJSA_1136 | T | S-ribosylhomocysteinase | 546 | 330 | 1.5E-06 | -1.7 |
| - | CJSA_1182 | C | radical SAM domain-containing protein | 63 | 43 | 3.4E-03 | -1.5 |
| *porA* | CJSA_1198 | - | major outer membrane protein | 18344 | 11845 | 4.5E-02 | -1.5 |
| *hydD* | CJSA_1203 | C | putative hydrogenase maturation protease | 472 | 250 | 3.6E-10 | -1.9 |
| *pseB* | CJSA_1231[d] | MG | UDP-GlcNAc-specific C4,6 dehydratase/C5 epimerase | 205 | 124 | 3.5E-07 | -1.7 |
| *pseC* | CJSA_1232[d] | M | C4 aminotransferase specific for PseB product | 124 | 67 | 6.9E-04 | -1.9 |
| - | CJSA_1233[d] | R | hypothetical protein | 44 | 22 | 4.9E-04 | -2.0 |
| *neuC2* | CJSA_1264 | M | putative UDP-N-acetylglucosamine 2-epimerase | 38 | 21 | 4.9E-02 | -1.8 |
| - | CJSA_1350[e] | H | putative methyltransferase | 2047 | 519 | 6.2E-12 | -3.9 |
| - | CJSA_1351[e] | H | putative methyltransferase | 1163 | 629 | 1.3E-02 | -1.8 |
| *flgI* | CJSA_1386[f] | N | lagellar basal body P-ring protein | 137 | 61 | 2.2E-04 | -2.2 |
| - | CJSA_1388[f] | - | hypothetical protein | 844 | 513 | 5.4E-05 | -1.6 |
| - | CJSA_1389[f] | - | hypothetical protein | 282 | 173 | 7.2E-03 | -1.6 |
| *flgK* | CJSA_1390[f] | N | flagellar hook-associated protein FlgK | 125 | 75 | 3.5E-07 | -1.7 |
| *moaE* | CJSA_1439 | H | putative molybdopterin converting factor,subunit 2 | 130 | 80 | 7.3E-04 | -1.6 |
| *nuoD* | CJSA_1488 | C | NADH dehydrogenase I subunit D | 164 | 93 | 7.7E-08 | -1.8 |
| - | CJSA_1562 | - | hypothetical protein | 138 | 80 | 9.4E-05 | -1.7 |
| - | CJSA_1568 | - | hypothetical protein | 4698 | 2214 | 9.9E-03 | -2.1 |
| - | CJSA_1577 | R | hypothetical protein | 246 | 168 | 4.0E-02 | -1.5 |
| *secY* | CJSA_1597[g] | U | preprotein translocase subunit SecY | 213 | 94 | 1.4E-15 | -2.3 |

**Table S6 continued**

| | | | | | | | |
|---|---|---|---|---|---|---|---|
| *rplO* | CJSA_1598[g] | J | 50S ribosomal protein L15 | 335 | 193 | 2.1E-03 | -1.7 |
| *rpsE* | CJSA_1599[g] | J | 30S ribosomal protein S5 | 463 | 202 | 6.7E-11 | -2.3 |
| *rplR* | CJSA_1600[g] | J | 50S ribosomal protein L18 | 288 | 161 | 3.9E-02 | -1.8 |
| *rplF* | CJSA_1601[g] | J | 50S ribosomal protein L6 | 397 | 208 | 2.2E-09 | -1.9 |
| *rpsH* | CJSA_1602[g] | J | 30S ribosomal protein S8 | 371 | 191 | 2.3E-04 | -1.9 |
| *rpsN* | CJSA_1603[g] | J | 30S ribosomal protein S14 | 541 | 247 | 1.2E-03 | -2.2 |
| *rplE* | CJSA_1604[g] | J | 50S ribosomal protein L5 | 481 | 252 | 3.6E-10 | -1.9 |
| *rplX* | CJSA_1605[g] | J | 50S ribosomal protein L24 | 609 | 316 | 4.7E-04 | -1.9 |
| *rplN* | CJSA_1606[g] | J | 50S ribosomal protein L14 | 554 | 300 | 1.7E-08 | -1.8 |
| *rpmC* | CJSA_1608[g] | J | 50S ribosomal protein L29 | 350 | 226 | 4.0E-02 | -1.5 |
| *rplP* | CJSA_1609[g] | J | 50S ribosomal protein L16 | 591 | 407 | 2.1E-02 | -1.5 |
| - | CJSA_CjSRP1 | | - | 4910 | 802 | 7.2E-96 | -6.1 |
| *ssrA* | CJSA_CjtmRNA1 | | - | 34176 | 20049 | 1.2E-03 | -1.7 |
| - | CJSA_t0002 | | Ile tRNA | 245 | 95 | 1.8E-07 | -2.6 |
| - | CJSA_t0005 | | Ile tRNA | 245 | 96 | 3.1E-07 | -2.6 |
| - | CJSA_t0007 | | Tyr tRNA | 661 | 396 | 1.2E-03 | -1.7 |
| - | CJSA_t0012 | | Met tRNA | 24 | 15 | 1.6E-05 | -1.6 |
| - | CJSA_t0013 | | Gln tRNA | 23 | 15 | 2.7E-05 | -1.5 |
| - | CJSA_t0015 | | Ile tRNA | 251 | 103 | 4.1E-06 | -2.4 |
| - | CJSA_t0017 | | Gly tRNA | 158 | 69 | 2.0E-03 | -2.3 |
| - | CJSA_t0018 | | Leu tRNA | 50 | 19 | 1.8E-06 | -2.6 |
| - | CJSA_t0020 | | Val tRNA | 93 | 39 | 1.9E-03 | -2.4 |
| - | CJSA_t0021 | | Arg tRNA | 312 | 165 | 2.3E-02 | -1.9 |
| - | CJSA_t0033 | | Leu tRNA | 754 | 318 | 4.3E-06 | -2.4 |
| - | CJSA_t0035 | | Ser tRNA | 121 | 50 | 2.3E-03 | -2.4 |
| - | CJSA_t0036 | | Leu tRNA | 213 | 67 | 2.5E-08 | -3.2 |
| - | CJSA_t0037 | | Arg tRNA | 800 | 368 | 9.8E-05 | -2.2 |
| - | CJSA_t0038 | | Arg tRNA | 1659 | 994 | 3.8E-03 | -1.7 |

**Table S6 continued**

| | | | | | | | |
|---|---|---|---|---|---|---|---|
| - | CJSA_t0039 | | His tRNA | 850 | 431 | 1.8E-05 | -2.0 |
| - | CJSA_t0043 | | Ala tRNA | 40 | 19 | 6.2E-11 | -2.1 |
| - | predicted RNA | | 16 nt length, - strand from 1577169 to 1577153 (IG CjSA 1568 /*nhaA1*) | 863 | 186 | 9.8E-18 | -4.6 |
| CjNC130 | predicted RNA | | 17 nt length, + strand from 1183929 to 1183946 (6S RNA) | 693 | 232 | 1.4E-05 | -3.0 |
| - | predicted RNA | | 55nt length,+ strand from 1132258 to 1132313 (3' end of *luxS*) | 36 | 13 | 5.6E-23 | -2.8 |
| - | predicted RNA | | 173nt length, + strand from 198877 to 199050 (antisense: CJSA_0192) | 2179 | 803 | 1.9E-25 | -2.7 |
| - | predicted RNA | | 37 nt length, - strand from 1358594 to 1358557 (IG CjSA_1350/CjSA_1351) | 2324 | 919 | 4.6E-13 | -2.5 |
| - | predicted RNA | | 198 nt length, + strand from 1457497 to 1457695 (CjSA 1444 internal) | 217 | 102 | 1.8E-02 | -2.1 |
| - | predicted RNA | | 15nt length, + strand pVir from 7643 to 7658 | 205 | 99 | 8.8E-08 | -2.1 |
| **Genes upregulated** | | | | | | | |
| *dnaN* | CJSA_0002 | L | DNA polymerase III subunit beta | 55 | 128 | 8.5E-04 | 2.3 |
| - | CJSA_0008[h] | S | hypothetical protein | 20 | 62 | 1.2E-09 | 3.1 |
| *gltD* | CJSA_0009[h] | ER | glutamate synthase subunit beta | 92 | 171 | 1.0E-02 | 1.9 |
| *folk* | CJSA_0059 | H | 2-amino-4-hydroxy-6-hydroxymethyldihydropteridine pyrophosphokinase | 67 | 108 | 4.9E-02 | 1.6 |
| - | CJSA_0076 | M | putative aspartate racemase | 43 | 76 | 1.7E-02 | 1.8 |
| - | CJSA_0105[i] | S | hypothetical protein | 156 | 362 | 1.4E-04 | 2.3 |
| - | CJSA_0110[i] | Q | putative pyrazinamidase/nicotinamidase | 46 | 88 | 2.3E-02 | 1.9 |
| *dgkA* | CJSA_0234[j] | M | diacylglycerol kinase | 29 | 75 | 1.6E-02 | 2.6 |
| *pyrC* | CJSA_0236[j] | F | Dihydroorotase | 25 | 55 | 1.5E-03 | 2.2 |
| *Tal* | CJSA_0257 | G | Transaldolase | 34 | 59 | 1.8E-02 | 1.7 |
| - | CJSA_0305 | - | hypothetical protein | 286 | 606 | 3.0E-03 | 2.1 |
| *trpD* | CJSA_0320[k] | E | anthranilate synthase component II | 72 | 133 | 2.0E-02 | 1.8 |
| *trpF* | CJSA_0321[k] | E | N-(5phosphoribosyl)anthranilate isomerase | 21 | 83 | 3.4E-09 | 4.0 |
| *trpB* | CJSA_0322[k] | E | tryptophan synthase subunit beta | 19 | 56 | 1.8E-06 | 2.9 |
| *trpA* | CJSA_0323[k] | E | tryptophan synthase subunit alpha | 12 | 29 | 4.2E-02 | 2.4 |
| - | CJSA_0337 | - | hypothetical protein | 9 | 24 | 2.5E-02 | 2.7 |
| - | CJSA_0370 | - | hypothetical protein | 22 | 55 | 2.9E-02 | 2.5 |
| - | CJSA_0372 | R | colicin V production protein-like protein | 90 | 214 | 1.2E-03 | 2.4 |

**Table S6 continued**

| | | | | | | | |
|---|---|---|---|---|---|---|---|
| *sdhA* | CJSA_0409[l] | C | succinate dehydrogenase, flavoprotein subunit | 30 | 70 | 2.3E-03 | 2.3 |
| *sdhB* | CJSA_0410[l] | C | succinate dehydrogenase, iron-sulfur protein subunit | 31 | 61 | 4.3E-03 | 2.0 |
| *sdhB* | CJSA_0411[l] | C | succinate dehydrogenase subunit C | 33 | 59 | 2.0E-02 | 1.8 |
| - | CJSA_0490[m] | - | hypothetical protein | 65 | 122 | 9.0E-03 | 1.9 |
| - | CJSA_0491[m] | P | Na/Pi-cotransporter, putative | 19 | 34 | 1.2E-02 | 1.8 |
| - | CJSA_0836 | TK | DNA-binding response regulator | 38 | 69 | 4.0E-02 | 1.8 |
| *Cfa* | CJSA_1121 | M | cyclopropane-fatty-acyl-phospholipid synthase | 22 | 52 | 3.2E-04 | 2.4 |
| *cetB* | CJSA_1127[n] | T | bipartate energy taxis response protein cetB | 28 | 97 | 1.8E-07 | 3.5 |
| *cetA* | CJSA_1128[n] | NT | bipartate energy taxis response protein cetA | 45 | 95 | 1.0E-02 | 2.1 |
| - | CJSA_1129 | T | putative PAS domain containing signal-transduction sensor protein | 57 | 108 | 8.8E-03 | 1.9 |
| - | CJSA_1131 | - | hypothetical protein | 10 | 33 | 1.3E-03 | 3.3 |
| - | CJSA_1145 | OC | putative lipoprotein thiredoxin | 82 | 159 | 2.8E-02 | 1.9 |
| - | CJSA_1164 | T | two-component sensor (histidine kinase) | 110 | 173 | 4.5E-02 | 1.6 |
| *cbpA* | CJSA_1167 | O | co-chaperone protein DnaJ | 22 | 54 | 4.7E-04 | 2.5 |
| - | CJSA_1259[o] | QR | methyltransferase domain-containing protein | 34 | 74 | 1.4E-03 | 2.2 |
| - | CJSA_1260[o] | QR | methyltransferase domain-containing protein | 26 | 65 | 1.0E-03 | 2.5 |
| - | CJSA_1301 | O | putative nucleotidyltransferase | 19 | 41 | 4.9E-03 | 2.2 |
| - | CJSA_1343 | - | hypothetical protein | 33 | 59 | 1.9E-02 | 1.8 |
| *tagF* | CJSA_1365 | M | putative CDP glycerol glycerophosphotransferase | 45 | 74 | 2.0E-02 | 1.6 |
| - | CJSA_1449 | R | putative helix-turn-helix containing protein | 43 | 96 | 2.5E-03 | 2.2 |
| *rloH* | CJSA_1466 | R | putative ATP/GTP-binding protein | 18 | 34 | 1.2E-02 | 1.9 |
| *nuoM* | CJSA_1479 | C | NADH dehydrogenase I subunit M | 76 | 144 | 1.8E-02 | 1.9 |
| - | CJSA_1549 | G | hypothetical protein | 31 | 76 | 3.0E-05 | 2.5 |
| *leuC* | CJSA_1626[p] | E | 3-isopropylmalate dehydratase large subunit | 9 | 22 | 3.4E-02 | 2.4 |
| *leuB* | CJSA_1627[p] | CE | 3-isopropylmalate dehydrogenase | 11 | 30 | 1.8E-02 | 2.7 |
| *leuA* | CJSA_1628[p] | E | 2-isopropylmalate synthase | 25 | 47 | 1.4E-02 | 1.9 |
| CjNC140 | predicted RNA | - | 92nt length, positive strand from 1193307 to 1193399 | 10 | 40 | 2.5E-06 | 4.0 |
| - | CJSA_pVir0042 | - | hypothetical protein | 10 | 42 | 2.5E-06 | 4.2 |

**Table S6 continued**

| | | | | | | | |
|---|---|---|---|---|---|---|---|
| - | CJSA_pVir0044 | U | hypothetical protein | 12 | 33 | 1.3E-02 | 2.8 |
| - | CJSA_pVir0025 | - | hypothetical protein | 9 | 24 | 5.3E-02 | 2.7 |
| | | | **Stationary phase** | | | | |
| **Genes downregulated** | | | | | | | |
| *rnhB* | CJSA_0010 | L | ribonuclease HII | 76 | 51 | 1.6E-02 | -1.5 |
| - | CJSA_0158 | - | hypothetical protein | 361 | 240 | 1.5E-03 | -1.5 |
| - | CJSA_0160 | QR | hypothetical protein | 303 | 207 | 2.7E-03 | -1.5 |
| - | CJSA_0284 | P | SMR family multidrug efflux pump | 41 | 27 | 1.6E-02 | -1.5 |
| *motB* | CJSA_0310 | N | flagellar motor protein MotB | 181 | 112 | 1.9E-04 | -1.6 |
| *trpD* | CJSA_0320 | E | anthranilate synthase component II | 22 | 11 | 3.1E-02 | -2.0 |
| - | CJSA_0344 | - | hypothetical protein | 231 | 149 | 5.0E-04 | -1.6 |
| - | CJSA_0372 | R | colicin V production protein-like protein | 190 | 122 | 6.3E-05 | -1.6 |
| - | CJSA_0389 | - | hypothetical protein | 474 | 322 | 2.7E-03 | -1.5 |
| - | CJSA_0396 | S | putative acidic periplasmic protein | 32 | 19 | 9.6E-05 | -1.7 |
| *sdhA* | CJSA_0409[l] | C | succinate dehydrogenase, flavoprotein subunit | 39 | 22 | 6.3E-05 | -1.8 |
| *sdhB* | CJSA_0410[l] | C | succinate dehydrogenase, iron-sulfur protein subunit | 38 | 23 | 4.9E-02 | -1.7 |
| *uxaA* | CJSA_0452[q] | G | putative altronate hydrolase N-terminus | 54 | 33 | 4.9E-03 | -1.6 |
| *uxaA* | CJSA_0453[q] | G | putative altronate hydrolase C-terminus | 42 | 28 | 4.6E-03 | -1.5 |
| - | CJSA_0560 | - | hypothetical protein | 193 | 111 | 1.1E-05 | -1.7 |
| - | CJSA_0616 | S | OstA family protein | 27 | 17 | 1.6E-02 | -1.6 |
| *trmD* | CJSA_0677 | J | tRNA (guanine-N(1)-)-methyltransferase | 32 | 17 | 5.2E-06 | -1.9 |
| *aroB* | CJSA_0951 | E | 3-dehydroquinate synthase | 91 | 62 | 1.7E-04 | -1.5 |
| - | CJSA_1137 | R | 2OG-Fe(II) oxygenase | 39 | 8 | 2.8E-52 | -4.9 |
| - | CJSA_1164 | T | two-component sensor (histidine kinase) | 66 | 42 | 4.5E-04 | -1.6 |
| *pyrH* | CJSA_1213 | F | uridylate kinase | 250 | 156 | 5.8E-03 | -1.6 |
| - | CJSA_1349 | G | hypothetical protein | 60 | 22 | 5.9E-20 | -2.7 |
| - | CJSA_1350 | H | putative methyltransferase | 387 | 170 | 8.2E-04 | -2.3 |

**Table S6 continued**

| | | | | | | | |
|---|---|---|---|---|---|---|---|
| - | CJSA_1414 | TK | putative two-component regulator | 100 | 57 | 3.8E-05 | -1.8 |
| *acs* | CJSA_1453 | I | acetyl-coenzyme A synthetase | 110 | 44 | 5.5E-13 | -2.5 |
| - | CJSA_1461 | R | MdaB protein-like protein | 66 | 40 | 3.9E-02 | -1.7 |
| *rnhA* | CJSA_1548 | L | ribonuclease H | 38 | 24 | 1.9E-03 | -1.6 |
| - | CJSA_1560 | Q | putative ABC transport system periplasmic substrate-binding protein | 25 | 15 | 3.1E-02 | -1.7 |
| *leuC* | CJSA_1626[p] | E | 3-isopropylmalate dehydratase large subunit | 58 | 29 | 2.1E-07 | -2.0 |
| *leuB* | CJSA_1627[p] | CE | 3-isopropylmalate dehydrogenase | 56 | 25 | 5.1E-07 | -2.2 |
| *leuA* | CJSA_1628[p] | E | 2-isopropylmalate synthase | 82 | 46 | 4.2E-06 | -1.8 |
| - | CJSA_pVir0030 | - | hypothetical protein | 198 | 117 | 5.5E-05 | -1.7 |
| - | predicted RNA | | 97nt length, - strand from 907368 to 907271 (antisense: CJSA_0911) | 139 | 86 | 8.1E-03 | -1.6 |
| - | predicted RNA | | 11nt length, + strand from 1397360 to 1397371 (IG CjSA_1387/CjSA_1388) | 1543 | 1061 | 4.5E-03 | -1.5 |
| **Genes upregulated** | | | | | | | |
| *dsbI* | CJSA_0017 | O | DsbB family disulfide bond formation protein | 41 | 87 | 2.4E-02 | 2.1 |
| - | CJSA_0040[a] | - | hypothetical protein | 111 | 330 | 1.6E-05 | 3.0 |
| - | CJSA_0041[a] | - | hypothetical protein | 74 | 281 | 1.8E-09 | 3.8 |
| *flgD* | CJSA_0042[a] | N | flagellar basal body rod modification protein | 157 | 481 | 3.8E-06 | 3.1 |
| *flgE* | CJSA_0043[a] | N | flagellar hook protein | 124 | 390 | 2.4E-13 | 3.1 |
| *flgC* | CJSA_0494[r] | N | flagellar basal-body rod protein FlgC | 559 | 911 | 1.2E-03 | 1.6 |
| *flgB* | CJSA_0495[r] | N | flagellar basal-body rod protein FlgB | 245 | 644 | 3.3E-02 | 2.6 |
| *flgH* | CJSA_0651 | N | flagellar basal body L-ring protein | 241 | 687 | 1.9E-07 | 2.9 |
| *flgG2* | CJSA_0661[c] | N | flagellar basal-body rod protein | 308 | 829 | 4.1E-12 | 2.7 |
| *flgG* | CJSA_0662[c] | N | flagellar basal-body rod protein FlgG | 292 | 886 | 9.2E-15 | 3.0 |
| *hcrA* | CJSA_0713 | K | heat-inducible transcription repressor | 306 | 722 | 1.2E-08 | 2.4 |
| - | CJSA_0716 | R | hypothetical protein | 25 | 51 | 5.1E-02 | 2.0 |
| *flgS* | CJSA_0749 | T | sensor histidine kinase | 13 | 32 | 3.6E-03 | 2.5 |
| *flgL* | CJSA_0833 | N | flagellar hook-associated protein FlgL | 176 | 503 | 2.1E-08 | 2.9 |
| - | CJSA_0969 | S | putative lipoprotein | 820 | 1267 | 3.7E-02 | 1.5 |
| - | CJSA_1107[s] | - | hypothetical protein | 50 | 141 | 1.6E-09 | 2.8 |

**Table S6 continued**

| | | | | | | | |
|---|---|---|---|---|---|---|---|
| *omp50* | CJSA_1108[s] | - | 50 kda outer membrane protein precursor | 57 | 148 | 2.4E-06 | 2.6 |
| - | CJSA_1180 | - | hypothetical protein | 642 | 1553 | 2.1E-07 | 2.4 |
| *pseB* | CJSA_1231 | MG | UDP-GlcNAc-specific C4,6 dehydratase/C5 epimerase | 208 | 450 | 8.5E-06 | 2.2 |
| - | CJSA_1262[t] | J | putative methyltransferase | 18 | 31 | 3.7E-02 | 1.7 |
| *neuB2* | CJSA_1263[t] | M | N-acetylneuraminate synthase | 58 | 157 | 5.7E-04 | 2.7 |
| *neuC2* | CJSA_1264[t] | M | putative UDP-N-acetylglucosamine 2-epimerase | 7 | 25 | 7.9E-16 | 3.6 |
| *flgI* | CJSA_1386[f] | N | lagellar basal body P-ring protein | 167 | 546 | 3.2E-11 | 3.3 |
| - | CJSA_1387[f] | - | hypothetical protein | 139 | 354 | 2.4E-02 | 2.5 |
| *flgK* | CJSA_1390[f] | N | flagellar hook-associated protein FlgK | 160 | 302 | 8.1E-05 | 1.9 |
| - | CJSA_1562 | - | hypothetical protein | 43 | 126 | 4.3E-10 | 2.9 |
| *p19* | CJSA_1570 | P | periplasmic protein p19 | 16 | 44 | 1.5E-09 | 2.8 |
| - | CJSA_pVir0042 | - | hypothetical protein | 64 | 110 | 5.1E-02 | 1.7 |
| - | CJSA_pVir0013 | - | hypothetical protein | 35 | 88 | 1.1E-04 | 2.5 |

[a, b, c, etc] = denotes genes within the same operon as predicted by Rockhopper

Underlined = signifcantly different expression at both timepoints
Double underlined = significantly different expression at both timepoints, opposite direction
IG = intergenic

**Table S7.** Rockhopper comparison of gene expression in the flagellar modification locus of IA3902 between wild-type and ΔCjNC110

| | | | Exponential | | | Stationary | | |
|---|---|---|---|---|---|---|---|---|
| | | | Expression (RMPK) | | Fold Change | Expression (RMPK) | | Fold Change |
| **Pathway** | **Name[#]** | **Synonym** | **WT** | **ΔCjNC110** | | **WT** | **ΔCjNC110** | |
| PseAc | *pseB* | CJSA_1231 | 205 | 149 | -1.38 | 208 | 208 | 1.00 |
| PseAc | *pseC* | CJSA_1232 | 124 | 86 | -1.44 | 103 | 112 | 1.09 |
| | - | CJSA_1233 | 44 | 32 | -1.38 | 22 | 21 | -1.05 |
| | - | CJSA_1234 | 3 | 3 | -1.00 | 2 | 1 | -2.00[b] |
| | - | CJSA_1235 | 9 | 10 | 1.11 | 4 | 3 | -1.33 |
| | - | CJSA_1236 | 5 | 8 | 1.60[b] | 1 | 2 | 2.00[b] |
| | *acpP2* | CJSA_1237 | 66 | 100 | 1.52[b] | 130 | 114 | -1.14 |
| | - | CJSA_1238 | 73 | 67 | -1.09 | 73 | 71 | -1.03 |
| | - | CJSA_1239 | 98 | 89 | -1.10 | 74 | 82 | 1.11 |
| | - | CJSA_1240 | 29 | 27 | -1.07 | 13 | 13 | 1.00 |
| | *fabH2* | CJSA_1241 | 52 | 48 | -1.08 | 20 | 24 | 1.20 |
| | *acpP3* | CJSA_1242 | 142 | 129 | -1.10 | 43 | 56 | 1.30 |
| | - | CJSA_1243 | 10 | 17 | 1.70[b] | 4 | 8 | 2.00[b] |
| | - | CJSA_1244 | 34 | 23 | -1.48 | 30 | 18 | -1.67[b] |
| | - | CJSA_1245 | 27 | 28 | 1.04 | 42 | 40 | -1.05 |
| | - | CJSA_1246 | 35 | 84 | 2.40[b] | 88 | 128 | 1.45 |
| | - | CJSA_1247 | 115 | 114 | -1.01 | 72 | 78 | 1.08 |
| | - | CJSA_1248 | 60 | 49 | -1.22 | 45 | 41 | -1.10 |
| PseAc | *pseF* | CJSA_1249 | 44 | 37 | -1.19 | 87 | 73 | -1.19 |
| PseAc | *pseG* | CJSA_1250 | 36 | 29 | -1.24 | 57 | 54 | -1.06 |
| PseAc | *pseH* | CJSA_1251 | 5 | 4 | -1.25 | 11 | 8 | -1.38 |
| | *hisF* | CJSA_1252 | 84 | 30 | **-2.80** | 56 | 31 | **-1.81** |
| | *hisH* | CJSA_1253 | 81 | 39 | -2.08[b] | 120 | 85 | -1.41 |
| PseAm | *pseA* | CJSA_1254 | 129 | 50 | **-2.58** | 261 | 152 | -1.72[b] |
| PseAc | *pseI* | CJSA_1255 | 91 | 79 | -1.15 | 82 | 87 | 1.06 |
| | *maf1* | CJSA_1256 | 23 | 18 | -1.28 | 30 | 21 | -1.43 |
| | - | CJSA_1257 | 67 | 50 | -1.34 | 34 | 26 | -1.31[a] |
| | - | CJSA_1258 | 29 | 17 | -1.71[b] | 11 | 9 | -1.22 |
| | - | CJSA_1259 | 34 | 0* | -34.00[b] | 20 | 0* | -20.00[b] |
| | - | CJSA_1260 | 26 | 0* | -26.00[b] | 15 | 0* | -15.00[b] |
| LegAm | - *(ptmG)* | CJSA_1261 | 34 | 66 | **1.94** | 10 | 29 | **2.90** |
| LegAm | - *(ptmH)* | CJSA_1262 | 25 | 49 | 1.96[b] | 18 | 27 | 1.50[b] |
| LegAm | *neuB2 (ptmC)* | CJSA_1263 | 171 | 57 | **-3.00** | 58 | 31 | **-1.87** |
| LegAm | *neuC2 (ptmD)* | CJSA_1264 | 38 | 28 | -1.36 | 7 | 9 | 1.29 |
| LegAm | - *(ptmE)* | CJSA_1265 | 42 | 31 | -1.35 | 13 | 10 | -1.30 |
| LegAm | - *(ptmF)* | CJSA_1266 | 30 | 23 | -1.30 | 43 | 16 | -2.69[b] |
| LegAm | *ptmB* | CJSA_1267 | 93 | 58 | -1.60[b] | 158 | 42 | **-3.76** |

| | | | | | | | | |
|---|---|---|---|---|---|---|---|---|
| LegAm | *ptmA* | CJSA_1268 | 107 | 41 | **-2.61** | 102 | 24 | **-4.25** |
| PseAm | *pseD* | CJSA_1269 | 16 | 7 | -2.29[b] | 12 | 4 | -3.00[b] |
| | *maf3* | CJSA_1270 | 29 | 28 | -1.04 | 15 | 10 | -1.50[b] |
| | *maf4* | CJSA_1271 | 21 | 15 | -1.40 | 28 | 19 | -1.47 |
| PseAc | *pseE* | CJSA_1272 | 51 | 58 | 1.14 | 36 | 39 | 1.08 |
| | *flaB* | CJSA_1273 | 2228 | 1700 | -1.31 | 4221 | 5100 | 1.21 |
| | *flaA* | CJSA_1274 | 4060 | 3061 | -1.33 | 10448 | 11846 | 1.13 |
| | - | CJSA_1275 | 27 | 26 | -1.04 | 6 | 8 | 1.33 |
| | *maf6* | CJSA_1276 | 21 | 22 | -0.95 | 6 | 8 | 1.33 |
| | *maf7* | CJSA_1277 | 28 | 28 | 1.00 | 9 | 13 | 1.44 |

**Bold** indicates significant differential expression (Q value <0.05, fold change >1.5)
[a] = Q value <0.05 but fold change <1.5
[b] = fold change >1.5 but Q value >0.05
* = actual value 0, however, 1 was utilized to calculate a fold change
[#] = currently annotated gene names with alternative gene names in ( )
PseAc and PseAm = pseudaminic acid pathway and derivatives
LegAm = legionaminic acid pathway and derivatives

**Table S8.** Plasmids used in this study.

| Strain | Description | Source or Reference |
|---|---|---|
| pGEM-T | linearized vectors with T overhang and β-galactosidase screening | Promega, Madison, WI |
| pCjNC110::cat | pGEM plasmid carrying CjNC110 deletion construct with $Cm^R$ | This study |
| pRRK | *Campylobacter* plasmid containing the *rrs-rrl* 16S/23S operon with $Kan^R$ | Muraoka and Zhang, 2011 |
| pRRK::CjNC110 | pRRK plasmid carrying CjNC110 insertion | This study |

$Kan^R$ = kanamycin resistance cassette

$Cm^R$ = chloramphenicol resistance cassette

**Table S9.** Primers used in this study.

| Primers | Sequence | Target | Use | Reference |
|---|---|---|---|---|
| CjNC110F2 | 5'-TTTGATTTGCGTTTTTGCAT-3' | CjNC110 | Cloning | This study |
| CjNC110R2 | 5'-ATCAAGAGCTTGAGCGAAGG-3' | CjNC110 | Cloning | This study |
| Cj1198F1 | 5'-AACTACTTCAAACATAAAATTTCCTTG-3' | luxS/CjNC110 | Cloning | This Study |
| Cj1199R3 | 5'-CCATGCAAAACCGGTAAAAAA-3' | luxS/CjNC110 | Cloning | This Study |
| CjNC110c1F | 5'-GCAATCTAGATGCATTCTTTAGATGAAGCCA-3' | CjNC110 | Cloning | This Study |
| CjNC110c1R | 5'-GACTGTCTAGAAATTCTTTGCCAAGTTTGAA-3' | CjNC110 | Cloning | This Study |
| LuxRTPCR-F | 5'-CCTTAGAACATTTATTCGCAGGAT-3' | *luxS* | RT-PCR | This Study |
| LuxRTPCR-R | 5'-GACAACCCATAGGTGAAATATCAAT-3' | *luxS* | RT-PCR | This Study |
| 16S-rRNA-F | 5'-TACCTGGGCTTGATATCCTA-3' | 16s-rRNA | RT-PCR | Han et al., 2008 |
| 16S-rRNA-R | 5'-GGACTTAACCCAACATCTCA-3' | 16s-rRNA | RT-PCR | Han et al., 2008 |
| SA1356F | 5'-TCCCATTTGGATGTTGTTGA-3' | CjSA_1356 | RNAseq | Luo et al., 2012 |
| SA1356R | 5'-CAGAACCTGGCCACAAACTT-3' | CjSA_1356 | RNAseq | Luo et al., 2012 |
| CjNC110-LNA | /5'DigN/ GCACATCAGTTTCAT/3'Dig_N/ | CjNC110 | Northern | This study |
